# A marmoset brain cell census reveals influence of developmental origin and functional class on neuronal identity

**DOI:** 10.1101/2022.10.18.512442

**Authors:** Fenna M. Krienen, Kirsten M. Levandowski, Heather Zaniewski, Ricardo C.H. del Rosario, Margaret E. Schroeder, Melissa Goldman, Martin Wienisch, Alyssa Lutservitz, Victoria F. Beja-Glasser, Cindy Chen, Qiangge Zhang, Ken Y. Chan, Katelyn X. Li, Jitendra Sharma, Dana McCormack, Tay Won Shin, Andrew Harrahill, Eric Nyase, Gagandeep Mudhar, Abigail Mauermann, Alec Wysoker, James Nemesh, Seva Kashin, Josselyn Vergara, Gabriele Chelini, Jordane Dimidschstein, Sabina Berretta, Benjamin E. Deverman, Ed Boyden, Steven A. McCarroll, Guoping Feng

## Abstract

The mammalian brain is composed of many brain structures, each with its own ontogenetic and developmental history. Transcriptionally-based cell type taxonomies reveal cell type composition and similarity relationships within and across brain structures. We sampled over 2.4 million brain cells across 18 locations in the common marmoset, a New World monkey primed for genetic engineering, and used single-nucleus RNA sequencing to examine global gene expression patterns of cell types within and across brain structures. Our results indicate that there is generally a high degree of transcriptional similarity between GABAergic and glutamatergic neurons found in the same brain structure, and there are generally few shared molecular features between neurons that utilize the same neurotransmitter but reside in different brain structures. We also show that in many cases the transcriptional identities of cells are intrinsically retained from their birthplaces, even when they migrate beyond their cephalic compartments. Thus, the adult transcriptomic identity of most neuronal types appears to be shaped much more by their developmental identity than by their primary neurotransmitter signaling repertoire. Using quantitative mapping of single molecule FISH (smFISH) for markers for GABAergic interneurons, we found that the similar types (e.g. *PVALB*+ interneurons) have distinct biodistributions in the striatum and neocortex. Interneuron types follow medio-lateral gradients in striatum but form complex distributions across the neocortex that are not described by simple gradients. Lateral prefrontal areas in marmoset are distinguished by high relative proportions of *VIP*+ neurons. We further used cell-type-specific enhancer driven AAV-GFP to visualize the morphology of molecularly-resolved interneuron classes in neocortex and striatum, including the previously discovered novel primate-specific *TAC3+* striatal interneurons. Our comprehensive analyses highlight how lineage and functional class contribute to the transcriptional identity and biodistribution of primate brain cell types.

**One-Sentence Summary:** Adult primate neurons are imprinted by their region of origin, more so than by their functional identity.

## Main Text

The mammalian brain’s complex functional diversity stems from its vast cellular and molecular repertoire. To provide a more complete understanding of the cell types across major cortical and subcortical structures in a non-human primate brain, we conducted a census of cell types of the adult marmoset brain based on their transcriptional profiles. Previous single cell sequencing studies of the marmoset brain focused on single brain regions (*1, 2*), or on specific cell classes across regions (*3, 4*). However, inclusion of both closely and distantly related brain structures and cell types can yield insights into the developmental and ontological relationships between them (*5*). Comprehensive transcriptomic cell type atlases have been produced for the mouse (*6–9*). Complementing these, recent transcriptomic datasets that sample many regions across humans and nonhuman primates brains (*10–12*) offer powerful resources for comparative analysis of brain cell type features. We generated single-nucleus RNA sequencing (snRNA-seq; 10x Genomics 3’ v3.1) data from 2.4 million unsorted brain cell nuclei across 8 neocortical and 10 subcortical locations from 10 young adult marmosets (4 M, 6 F), and resolved clusters from all major neuronal and non-neuronal cell classes. snRNA-seq data were generated as part of the Brain Initiative Cell Census Network (BICCN, RRID:SCR_015820) and are available on the BICCN Data Center (RRID:SCR_022815; https://biccn.org/data) as well as the NeMO archive (<u>RRID:SCR_016152</u>; https://doi.org/10.1101/2022.10.18.512442).

All neuron-containing brain structures in the central nervous system possess both excitatory and inhibitory neuronal populations, though the proportions and degree of developmental relatedness between these two populations varies by structure. In the neocortex and other telencephalic structures of vertebrates, distinct populations of neurons are typically categorized based on their neurotransmitter status as either inhibitory (GABAergic) or excitatory (glutamatergic). In other brain structures, primary neurotransmission appears less essential to a cell type’s identity. We found that the transcriptional identities of excitatory and inhibitory neurons within telencephalic brain structures segregate strongly, consistent with previous studies in other mammalian species (*2, 6, 7, 13*). In contrast, there is much greater transcriptional similarity between GABAergic and glutamatergic neuronal types in non-telencephalic compartments. Moreover, few gene expression distinctions present in telencephalic glutamatergic neurons are shared with glutamatergic neurons in non-telencephalic brain regions. While primary neurotransmission did not drive transcriptional similarity between neurons, their brain structure of origin did; the adult transcriptomic identity of most neuronal types appear to be shaped much more by their developmental identity than by their neurotransmitter signaling repertoire.

In the mouse, overall interneuron/excitatory neuron ratios are thought to be largely invariant across the neocortex (*14*). Even so, relative proportions of interneuron subtypes do vary and underlie principles of functional organization in the mouse neocortex (*15*). The small size of the marmoset brain and its near-lissencephalic neocortex enables quantitative, cell-type-resolved mapping in a primate. We used smFISH to spatially profile major interneuron types across marmoset striatum and neocortex. In the striatum, major interneuron types are distributed as medial-lateral gradients. In contrast, the marmoset neocortex has a much more complex topography of interneuron concentrations that is not explained by a single spatial axis. Lateral prefrontal areas in particular, which have undergone expansion in primate evolution (*16*) are typified by higher proportions of *VIP*+ and *LAMP5/ID2*+ interneurons and lower proportions of *SST*+ interneurons relative to all other neocortical areas.

The development of viral tools to effectively deliver transgenes to specific primate brain cell types has many powerful applications (*17–20*). We used this approach to sparsely label and reconstruct the morphologies of selected types. We reconstructed a set of molecularly-characterized neurons that had received systemic delivery of an AAV carrying a reporter (AAV9-hDlx5/6-GFP-fGFP) under an interneuron-selective regulatory element(*19*). Reconstructions are available for download from the Brain Image Library (BIL; https://doi.org/10.35077/g.609). As AAV9-hDlx5/6-GFP-fGFP did not label a recently discovered, primate-specific striatal interneuron type (*3*) w(*3*)This enabled visualization(*3*).

Together, our census of major transcriptomically-defined brain cell types and quantitative mapping of interneuron biodistributions provides a key resource for the primate neuroscience community and for comparative studies of cell type evolution.

### A transcriptomic census of marmoset brain cell types

We acquired snRNA-seq data from 18 brain regions collected using 10x Genomics 3’ 3.1 chemistry across 10 young adult marmosets (ages 1.5-4; 6 F) as well as a small dataset from PFC of two aged animals (**Fig. 1, Fig. S1, Table S1**). The number of donors per brain structure varied (min = 2, max = 10; **Fig. S1**), as did the cell sampling rate per brain structure (**Fig. 1A; Fig. S1A, Table S2**). Neocortex was the most comprehensively sampled in terms of total numbers, donors, and regional dissections. We acquired data from: cerebellum, brainstem, hypothalamus, thalamus, amygdala, striatum (separate dissections for caudate, putamen and nucleus accumbens), hippocampus, basal forebrain and neocortex. Within the neocortex, we separately sampled 8 neocortical locations (prefrontal, temporal pole, S1, M1, A1, V2, V1, lateral parietal association), and within PFC, from four prefrontal subdivisions (**Fig. S1B-C**).

**Figure 1.**
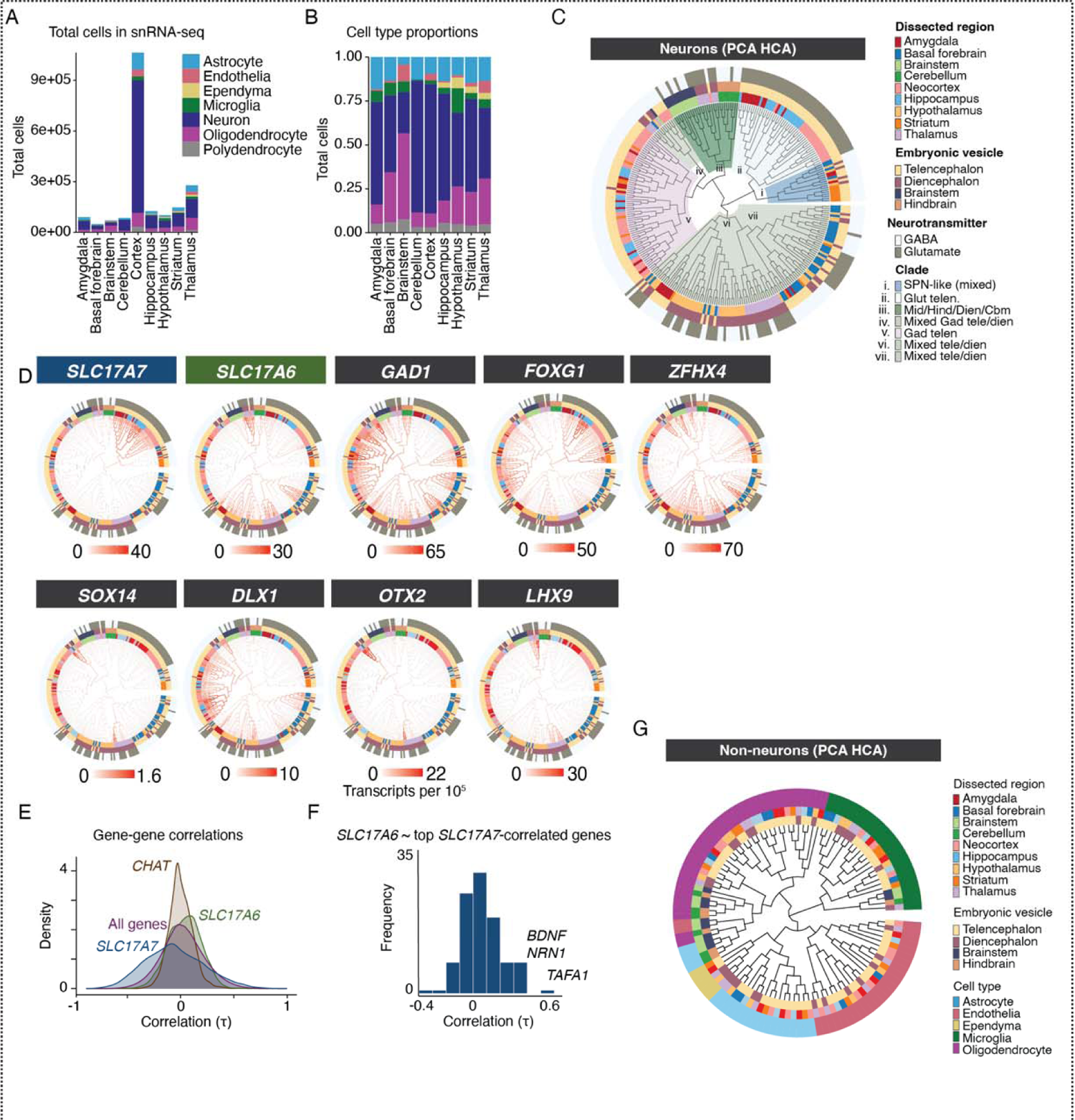
Single nucleus RNA sequencing of marmoset brain. **(A)** Number of nuclei per brain structure. Colors indicate cell classes. **(B)** Proportions of cell classes across brain structures. **(C)** Neurons in each dissected region are clustered separately, then pseudo-bulked “metacells” of all 288 clusters are arranged in the dendrogram using hierarchical clustering of top 100 PCA scores of expressed genes. Outer rings colored by dissected subregion (1st ring), cephalic vescicle (2nd ring) and major neurotransmitter usage (3rd ring). Seven major “clades” are colored and labeled according to cell type and regional composition of clade. **(D)** Expression of markers for glutamatergic neurons (*SLC17A7*, *SLC17A6*), GABAergic neurons (*GAD1*), and transcription factors (*FOXG1*, *ZFHX4*, *SOX14, DLX1, OTX2, LHX9*) are plotted as heatmaps on dendrogram shown in (C). These transcription factors are largely restricted to specific clades or are associated with particular cephalic vesicles. **(E)** Gene-gene correlation (Spearman tau) distributions across all neuron clusters in (C) for *SLC17A7*, *SLC17A6*, and *CHAT* (marker for cholinergic neurons) as well as background (all genes). The distribution of pairwise correlations to *CHAT* had a lower standard deviation (mean *r* = 0.002, std. dev = 0.116) relative to baseline gene-gene correlations (mean *r* = 0.02, std. dev = 0.199; F(5078, 17315809) = 0.33, *p*-adj < 1e-15). In contrast, pairwise correlations to *SLC17A7* were much broader relative to the background distribution of all gene-gene correlations. **(F)** Distribution of cross-cell-type correlation to *SLC17A6* of genes most correlated with *SLC17A7* (top 116 genes with Spearman tau > 0.5 to *SLC17A7*). **(G)** Same hierarchical clustering procedure as (C) but for non-neuronal cell types. Outer ring colors indicate major non-neuronal cell class. Inner ring colors indicate region dissection and vesicle; colors as in (C).

Nuclei from each brain structure were pooled across donors and analyzed to identify major cell types and their proportions by brain structure (**Fig. 1B-D**). Using linear discriminant analysis (scPred;(*21*)) trained on a supervised set of cell class labels, we identified and discarded low quality cells and doublets, and assigned each nucleus to its probable major type – astrocyte, endothelia, ependyma, microglia/macrophage, neuron, oligodendrocyte, or oligodendrocyte precursor cell (OPC).

After major cell type assignment, nuclei were clustered by brain structure to reveal subtype diversity within each major class. We used a previously described clustering pipeline (*6*) based on independent components analysis (fastICA). Each clustering analysis involved additional curation of doublets and outlier cells, followed by a second round of sub-clustering of major clusters. At each of these curation stages, independent components that loaded on individual donors or batches were excluded and reclustering was performed to attenuate donor and batch effects on clustering results. For each cluster, a “metacell” was generated by summing transcript counts of individual cells of each type together, followed by scaling and normalizing to 100,000 transcripts/nucleus. These regional and cell-type resolved metacells were the starting point for most cross-region analyses. The final snRNA-seq dataset size after curation contained 2.01 million cells (**Table S2)**.

### Hierarchical clustering of neurons

Cell types are identified based on factors including their function, developmental origin, lineage, and regional context (*22–24*). We studied how transcriptional profiles related to each other across 288 neuronal cell types (metacells) sampled across all 18 marmoset brain structures. Individual replicates and other variables (age, sex) were generally proportionately represented across resolved cell types with several exceptions that were likely mainly to differences in dissection across donors (**Fig. S1B-E, Fig S2**). We used hierarchical clustering to situate the neuronal types on a dendrogram computed using distances calculated from the top 100 PC scores across neuron types (**Fig. 1C**). Most telencephalic types (neocortex, amygdala, hippocampus, striatum) clustered distinctly from diencephalic and hindbrain types, indicating that developmental lineage continues to shape the transcriptional identity of adult primate brain cell types. However, of the 7 major clades, 4 contained mixtures of cell types at the terminal (leaf level) originating from distinct cephalic compartments. For example, basal forebrain neuron types intermingled with hypothalamic types, suggesting closer transcriptional similarity of two structures that occupy distinct cephalic compartments. The overall dendrogram configuration was broadly conserved when recomputed using other distance functions (**Fig. S2A**).

Transcription factors are master regulators that determine cell type identity in development through temporal patterning, suggesting that they may be a key class of genes that determine transcriptional identity in adults (*8*). Supporting this view, some transcription factors are associated with specific brain structures or cephalic compartments, such as *FOXG1* in the telencephalon (**Fig. 1D**) and *OTX2* in non-telencephalic structures **(Fig. 1D)**, show expression restricted to specific clades in the dendrogram.

Hierarchical clustering based only on transcription factor genes was highly similar to the original tree computed over all expressed genes. However, the tree ordering generated by transcription factors alone did not produce lower tree distances to the original tree than similarly sized sets of randomly selected expressed genes (**Fig. S2D**). This suggests that though transcription factors undoubtedly play central roles in determining cell identity in development, they do not determine global transcriptomic similarities across neuron types in adulthood.

To determine whether a broadly similar transcriptomically-defined dendrogram is conserved across species, we repeated the analysis using a prior single cell RNA sequencing dataset of regions sampled across the mouse brain (*6*) (**Fig. S2C,D**). We found a conserved tree structure in the mouse data, suggesting broad conservation of the features that drive transcriptomic similarities in neurons across adult primates and rodents. The relative ordering of the major clades was strikingly similar across species: for example, in both species, telencephalic glutamatergic neurons are most similar to a clade of GABAergic neurons that includes striatal spiny projection neurons (SPNs) as well as transcriptionally similar types in amygdala and hypothalamus (**Fig. S2C**). In both species, all telencephalic GABAergic interneurons formed a large single clade, and cerebellar types were most similar to a mixed clade containing thalamic and brainstem types.

Borrowing from ancestral state reconstruction methods typically used to estimate evolutionary divergences between genetic sequences or species (*25*), we applied a maximum likelihood-based approach (fastAnc) to the expression of all genes at each leaf (observed cell type), along with the branch lengths between adjacent leaves, in order to reconstruct the transcriptomic state (inferred expression values of genes) at internal nodes of the marmoset dendrogram. We then compared pairwise “ancestral” expression values of all genes in the parent nodes for each of the 7 major clades depicted in **Fig. 1C, Fig. S2E-F)**. Transcription factors (TFs) were overrepresented in comparisons of internal nodes that contained cell types stemming from developmentally related brain regions, but were not overrepresented when the leaf cell types stemmed from multiple regions (**Fig. S2E-F**). While transcription factor expression reflects the developmental origins of cell types, and while their expression alone recapitulates the tree structure seen by including all genes (**Fig. S2D** and (*8*)), some cell types are transcriptionally similar despite having distinct developmental origins. This may reflect convergence in adult transcriptional profiles (*5, 26*).

### Many adult neuronal types are imprinted by their developmental origin

We next inspected RNA expression patterns along with details about each cell type’s dissection region-of-origin to assess which brain structures tended to contain highly similar cell types and which had more dispersed transcriptional profiles. Some tissues, such as the neocortex, gave rise to cell types that exclusively clustered within their cephalic domain (**Fig. 1C**), a result unlikely to be driven by ambient RNA contamination since regionally variable genes were not over-enriched in ambient RNA estimates (**Fig. S5A**). However, within a given cephalic domain, cell types from distinct brain structures were often more similar to types sampled from other brain structures. For example, while hippocampal cell types were all found in telencephalic clades in the dendrogram, many individual hippocampal types were more similar to types in the amygdala or neocortex than they were to other hippocampal types (**Fig. 1C**).

Out of 62 neocortical neuron types, only two types joined clades outside of the major GABAergic and glutamatergic telencephalic branches: (1) the *MEIS2*+ prefrontal GABAergic types (**Fig. S1C; Fig. 1D; Fig. S3**), which formed a clade most similar to diencephalic and midbrain *OTX2+* types, and (2) Neocortical Cajal-Retzius (C-R) neurons, which were more similar to a clade of *LHX9*+ thalamic neurons despite indications that they predominantly originate from the cortical hem in primates (*27*) (**Fig. 1D**). As with hippocampal neurons, neocortical GABAergic neurons neighbored other GABAergic neurons from the striatum, amygdala and hippocampus that tended to express the same marker. For example, all *PVALB*+ types sampled across these telencephalic structures grouped together (**Fig. S3**). Unlike neocortical GABAergic interneurons, glutamatergic neocortical neurons were broadly most similar to other telencephalic glutamatergic types, but showed almost no mixing at the terminal leaf level (**Fig. 1C**). Cerebellar neuron types were entirely restricted to a single, unmixed clade in both mouse and marmoset (**Fig. S2C**), though this relative isolation could be due to the lack of other hindbrain structures in our datasets (**Fig. 1C**).

Consistent with previous work suggesting that mammalian thalamus contains both midbrain-derived and forebrain-derived GABAergic interneurons (*28*), we observed distinct clades of thalamic neurons that were most similar to diencephalic or midbrain populations. Interestingly, thalamic GABAergic neurons that were *OTX2*+ were distinct from other thalamic populations (**Fig. 1D, Fig. S3**), and formed a clade with other *OTX2*+ neurons sampled from brainstem, hypothalamus, and basal forebrain. These populations were *SOX14*+, while thalamic *SOX14*-populations joined mixed diencephalic-telencephalic clades (**Fig. 1D; Fig. S3**). The dispersion of thalamic neurons to distinct diencephalic and midbrain-dominated clades supports recent work suggesting multiple developmental origins for primate thalamic GABAergic neurons (*28*). We did not find thalamic GABAergic populations that were transcriptionally similar to telencephalic types (*29*), though we note that the next most proximal clade to the thalamic *OTX2*+ types were *MEIS2*+ GABAergic neocortical neurons **(Fig. 1D)**.

Amygdala neuron types were distributed to 5 of the 7 major clades (**Fig. 1C**). Consistent with this widely dispersed cellular profile, the amygdala is composed of loosely associated nuclei with diverse phylogenetic and developmental origins, as documented by previous studies (*30, 31*). Specifically, the basolateral amygdala, which has a high proportion of excitatory neurons, shares properties with cortical and claustrum neurons, whereas the intercalated nuclei of the amygdala contain inhibitory *FOXP2*+ projection neurons that share developmental origins with some striatal GABAergic populations such as striatal spiny projection neurons, SPNs (*5, 32, 33*). In our dataset, glutamatergic amygdalar neurons were most similar to neocortical and hippocampal glutamatergic neurons, while *FOXP2*+ GABAergic amygdalar neurons clustered with SPN types (**Fig. S3**), in line with recent lineage tracing studies in mice (*5*) and with analysis of fetal macaque types (*33*). Amygdala GABAergic interneuron types had highly divergent transcriptomic profiles; while some were most similar to telencephalic GABAergic neurons, others clustered with hypothalamic GABAergic neurons. These results underscore the diversity and developmental complexity of cell types comprising the amygdala nuclei.

### Neurotransmitter usage is not strongly associated with transcriptomic identity

The neurotransmitters used by neurons are central to their function, and neurons are often classified by their primary neurotransmitter. However, synthesis of the major neurotransmitters – glutamate (in mammals, excitatory) and GABA or glycine (inhibitory) – requires only a modest number of genes, and GABA itself is synthesized from glutamate (*34*). It is unclear whether many or few other genes are correlated with GABAergic (or glutamatergic) identity. For instance, whether neurotransmitter utilization is strongly associated to the identity of neuron types may differ by brain structure: transcriptomically-defined neuron types in the neocortex and other telencephalic structures are hierarchically grouped into GABAergic and glutamatergic types (*2, 7, 26*), which reflects both their distinct developmental origins and their distinct neurotransmitter repertoire. In other brain structures such as the hypothalamus, the distinction between neurons utilizing GABA or glutamate is much less clear at the transcriptional level (*8, 35*) (**Fig. 1C,E,F, Fig. S3**).

To determine whether neurotransmitter identity was associated with a general transcriptional identity across all neurons expressing the same neurotransmitter, we examined expression of genes encoding the most prevalent vesicular glutamate transporters (*SLC17A6, SLC17A7*) and GABA synthesis enzymes (*GAD1, GAD2*) (**Fig. 1C-F; Fig. S3**). If primary neurotransmitter usage was strongly associated with the expression of many other genes, we would expect that neurons belonging to the same neurotransmitter set (GABAergic or glutamatergic) would preferentially group together regardless of other factors (such as developmental origin). However, we did not find evidence for strong global transcriptomic identities of GABAergic and glutamatergic neurons. Neuronal types from each set were distributed across the tree, suggesting divergent transcriptomic identities of cell types that share a common neurotransmitter **(Fig. 1C).**

In telencephalic structures such as neocortex, hippocampus, and amygdala, glutamatergic neurons all express *SLC17A7* and segregate from GABAergic telencephalic neurons (**Fig 1C,D**). Telencephalic GABAergic neurons separated into distinct clades: one for GABAergic interneurons, and another for GABAergic projection neurons such as SPNs of the striatum (**Fig 1C, Fig S2)**. In non-telencephalic structures such as the hypothalamus, basal forebrain, thalamus, brainstem, and cerebellum, GABAergic and glutamatergic types were highly intermixed. This pattern held in a mouse scRNA-seq dataset, and when using different approaches for hierarchical clustering **(Fig. S2)**. Even within the telencephalon, neurotransmitter identity did not drive global transcriptional similarity between major clades. For instance, GABAergic projection neurons, such as SPNs of the striatum, were transcriptionally closer to glutamatergic neurons in the neocortex, amygdala, and hippocampus than they were to telencephalic GABAergic interneurons **(Fig. 1C; Fig. S3)**.

Although glutamatergic neurons from distinct cephalic origins do not cluster together, maintaining glutamatergic neurotransmission or associated function could require a common, core set of genes. To assess how neurotransmitter utilization relates to genome-wide RNA expression patterns, we examined distributions of gene-gene correlations across cell types (**Fig. 1E**). Surprisingly few genes are strongly positively correlated with both *SLC17A6* (VGLUT2) and *SLC17A7* (VGLUT1) expression, even those associated with glutamate synthesis and packaging **(Fig. S3)**. 116 genes had correlated expression to *SLC17A7* (Spearman’s tau > 0.5). The median correlation of *SLC17A6* to those 116 genes was centered at tau = 0.05 (**Fig. 1F**). Only a few genes correlated above 0.5 to both *SLC17A6* and *SLC17A7,* including *BDNF, NRN1*, and *TAFA1.* Moreover, only 10 genes (*ARPP21, BDNF, CACNA2D1, CHN1, CHST8, CPNE4, LDB2, NRN1, PTPRK, TAFA1*) are differentially expressed (> 2.5 fold change) in both *SLC17A6*+ glutamatergic neurons and *SLC17A7*+ glutamatergic neurons relative to *GAD1*+ neurons. We examined the principal component loadings to see if any were associated with primary neurotransmission. While individual PCs loaded strongly on specific parts of the tree (**Fig. S2B**), none of the top 20 PCs (accounting for 92% variance) distinguished cell types by their neurotransmission. The bulk of gene expression that distinguishes neuronal types from each other appears incidental to neurotransmission.

### Neocortical expression fingerprints differ between neurons and glia

The mammalian neocortex is partitioned into functionally, connectionally, and cytoarchitectonically distinct regions, called areas. We examined regional distinctions in proportions and gene expression of cell types from 8 cortical locations (**Fig. 2A**). Consistent with prior reports (*3, 6, 13, 36*), cell subtypes identified in one cortical region were generally present in all other cortical regions, though in different proportions (**Fig. 2B**). There were two main exceptions in neurons. GABAergic *MEIS2*+ cells (GABAergic cluster 6, **Fig. 2B**) were far overrepresented in PFC samples (specifically in dissections of medial and medio-orbital PFC, **Fig. S1C**), a compositional distinction not observed in mouse (*6, 36*). The second exception was a cluster of *RORB+, KCNH8+* glutamatergic neurons in V1 (and to a lesser extent V2) that diverged from *RORB*+ populations found in the other cortical regions (Glutamatergic cluster 2, **Fig. 2B-D**). The expansion and divergence of *RORB*+ populations in visual cortex is consistent with the elaboration and sub-specialization of primary V1 layer IV in primates (*37*). One astrocyte type, marked by high expression of *VCAN*, was highly enriched in V1 and V2 (Astrocyte cluster 3, **Fig. 2D**). Using smFISH, we validated higher co-localization of *VCAN* and *GFAP* in V1-adjacent white matter compared with PFC and V1 gray matter (**Fig. S4I**). *VCAN* is also expressed in OPCs. We did not observe higher *VCAN* expression in V1 OPCs, suggesting that the regional variation in *VCAN* expression is specific to astrocytes (**Fig. 2E**).

**Figure 2.**
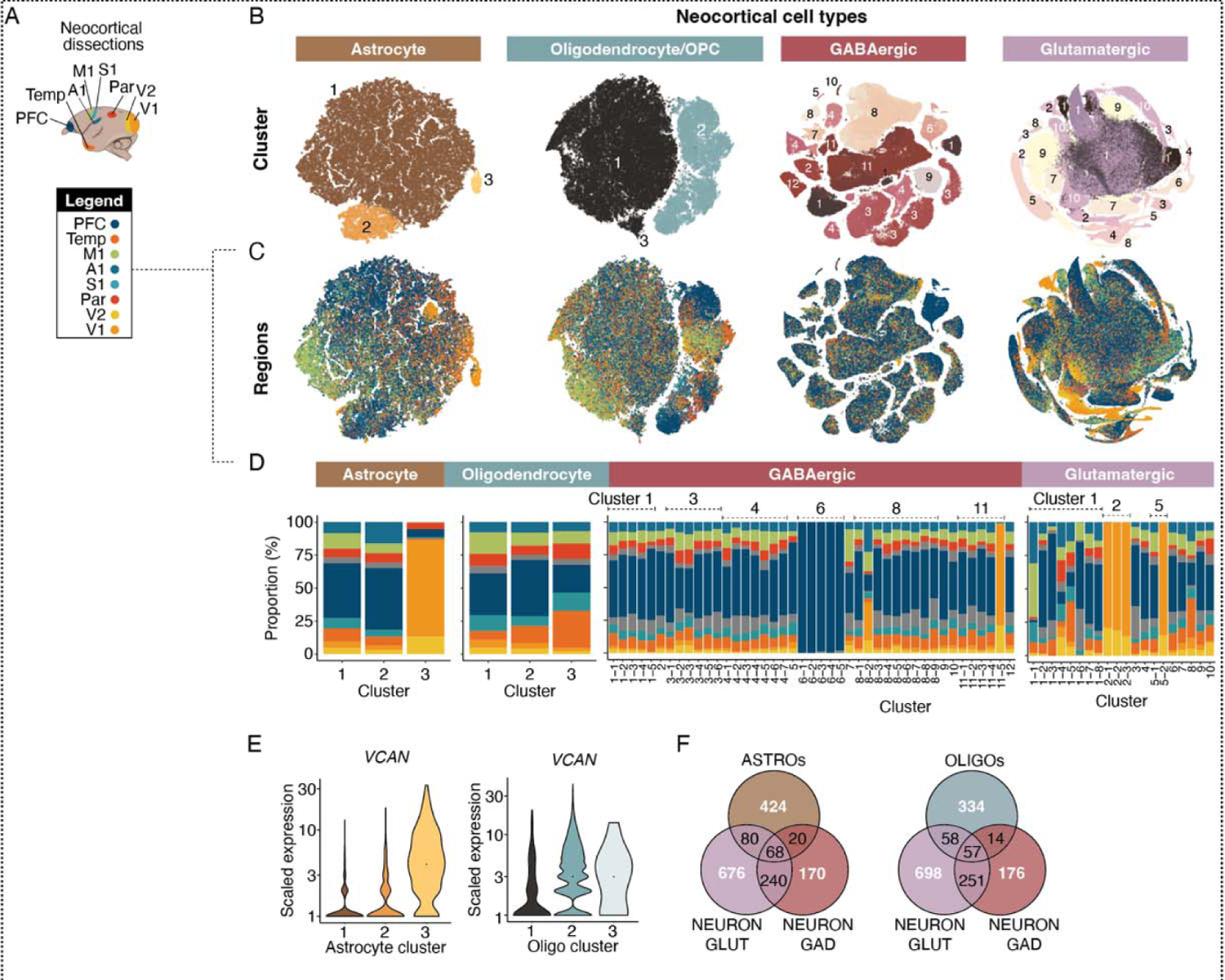
Regional variation in neocortical cell types and expression patterns. **(A)** Cortical regions sampled. **(B)** t-SNE embeddings of neocortical macroglia (astrocytes, oligodendrocyte lineage types) and neurons (GABAergic interneurons, glutamatergic neurons). Colors represent clusters (numbered). **(C)** Same as (B) but cells colored by neocortical dissection. **(D)** Regional proportions of each cluster, colors same as (A). **(E)** *VCAN* expression across astrocyte clusters and oligodendrocyte lineage clusters. Colors as in (B). **(F)** Venn diagrams showing overlap of neocortical regionally differentially expressed genes (rDEGs) across GABAergic neurons, glutamatergic neurons, astrocytes and oligodendrocyte lineage cells. rDEGs are defined as > 3 fold expression difference in homologous cell types across pairs of cortical regions.

Glutamatergic neurons sampled from different neocortical locations show regionally distinct gene expression (*36*). In primates, this is also true of neocortical GABAergic neurons (*3*). Astrocytes in mouse and primate brains are heterogeneous across major subdivisions (**Fig. S4D-H**), but the extent to which they are locally customized in distinct regions of neocortex is less well understood (*2, 7, 26, 38*)(*4*). Studies in mice have revealed layer-specific astrocyte subpopulations in the cortex (*39*) and variation between neocortical and hippocampal astrocytes (*40*).

To address whether cortical regional variation in gene expression is shared across cell types, we performed pairwise comparisons between major clusters of cortical excitatory neurons, inhibitory neurons, astrocytes and oligodendrocyte lineage types across all eight neocortical locations **(Fig. 2F, Table S3)**. Each of these cell classes displayed regionally differentially expressed genes (rDEGs) across neocortical regions (**Fig. 2F**), an effect that could not be attributed to ambient RNA contamination (**Fig. S5A**). Similar to what has been described in the mouse cortex (*39, 40*), astrocytes within the marmoset cortex exhibited regional transcriptional variation, but overall neurons had more rDEGs than macroglia (**Fig. 2F; Fig. S5B**). 62% of rDEGs in interneurons were also rDEGs in excitatory neurons. Interestingly, though cortical astrocytes and oligodendrocytes arise from a common lineage with cortical excitatory neurons (*41*), they shared a lower percentage of rDEGs in common with excitatory neurons as compared with interneurons (25% astrocytes, 25% oligodendrocyte lineage). Regionally differentially expressed genes within a cell class (glutamatergic, GABAergic, astrocyte, oligodendrocyte lineage) tend to be biased in the same regions across subtypes within that class. However, certain subtypes and regions accumulated more rDEGs than others. For example, across all cell types and particularly within neurons, higher-order temporal association cortex and prefrontal cortex tended to be most distinct from V1 and V2 (**Fig. S5B**). To determine the extent to which rDEGs are private to individual donors or represent shared features of variability within conserved cell types, we analyzed the consistency of neocortical rDEGs across the three donors that had been sampled at all eight neocortical locations (**Table S3**). Our findings revealed that 55% and 39% of glutamatergic and GABAergic rDEGs, respectively, were consistent among donors, while only 24% and 14% of astrocyte and oligodendrocyte rDEGs, respectively, were shared between donors. These results indicate lower inter-individual consistency in regional gene expression signatures in non-neuronal cells.

### A hypothalamic-origin neuron type in medial amygdala

In mammalian CNS, some cell types migrate long distances from proliferative zones to their mature destinations (*42, 43*). However, neurons generally tend to respect cephalic boundaries and remain within the same subdivision as their progenitors (*44*). This tight control over migration potential makes it difficult to disentangle the persistent influence of developmental origin from potential later influences arising from shared tissue context that might affect all neurons similarly in a given brain structure.

However, cephalic boundary crossings do exist (*28, 29, 45*). Though rare, such boundary crossing events can reveal whether cell types that embarked on cross-cephalic migration retain transcriptomic profiles more in common with their tissues of origin, or more in common with their final destinations.

We found a striking example of cross-cephalic migration in the amygdala. First, we observed an unexpected clustering pattern in our analysis, wherein three amygdala neuron types joined a clade not with other telencephalic neurons, but instead with *SLC17A6*+ hypothalamic and basal forebrain types (**Fig. 3A**). Moreover, despite expressing *SLC17A6* and lacking expression of *GAD1* and *GAD2,* these cells showed a closer association with GABAergic rather than glutamatergic types (**Fig. 3B**). They did not express other genes required for GABAergic transmission such as *SLC32A1* (VGAT) and also lacked the molecular machinery for non-canonical GABA reuptake or release observed in other cell populations (**Fig. S3**)(*46, 47*). These findings suggest that these particular amygdalar neurons exhibit a “cryptic” transcriptomic identity, in which they are glutamatergic but have transcriptomic profiles that are much more similar to GABAergic types, relative to other telencephalic neurons.

**Figure 3.**
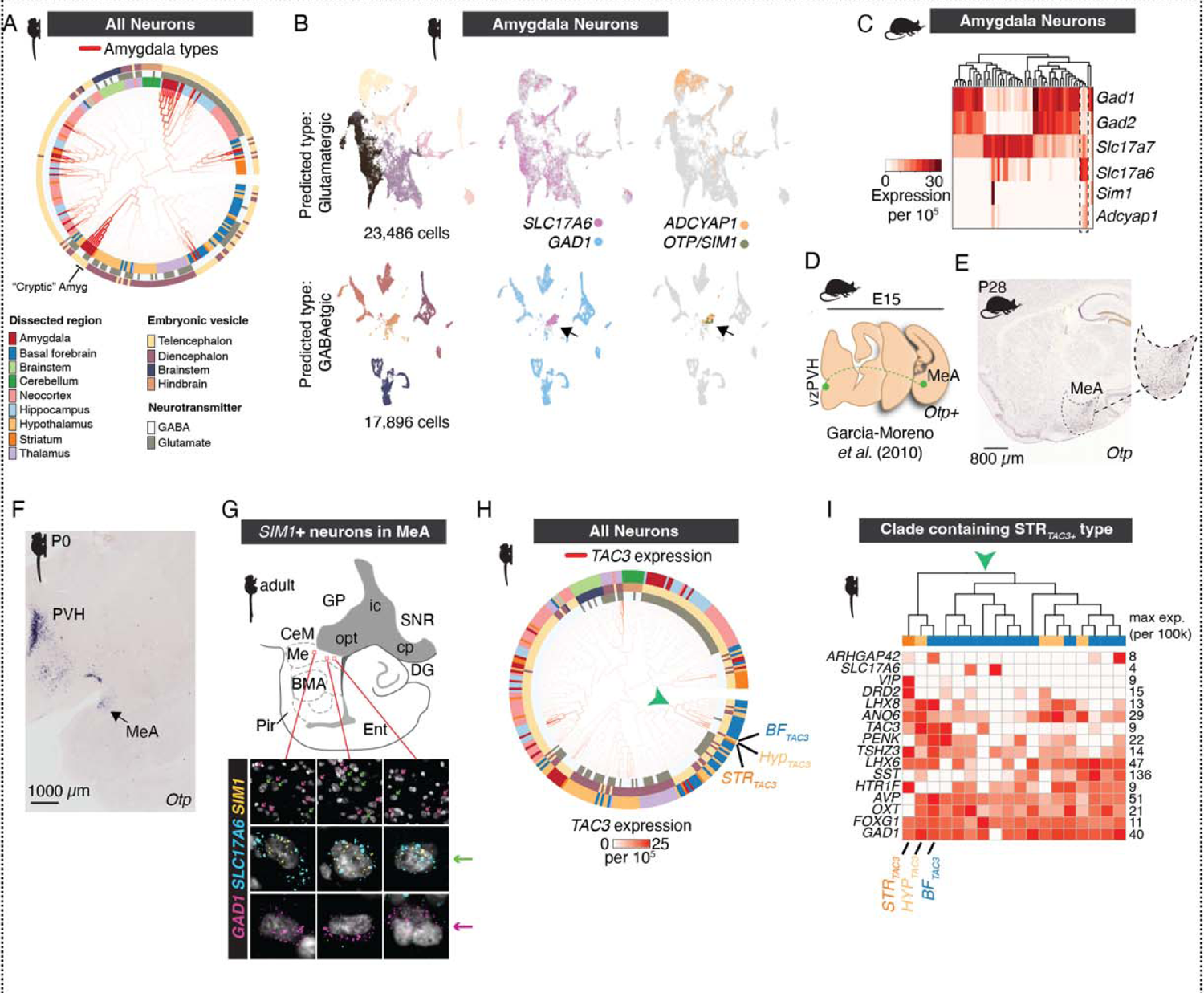
Examples suggesting cross-cephalic-boundary cell type migration. **(A)** Locations of amygdala clusters in the dendrogram from Fig. 1C. **(B)** Clustering of marmoset amygdala cells (n = 44,165 neurons) predicted to be glutamatergic (top row) or GABAergic (bottom row) based on a linear classifier (scPred) trained on supervised cell type models. t-SNEs for each class are colored by cluster (first column) or genes (*SLC17A6*, *GAD1, ADCYAP1, OTP, SIM1*). Arrowhead indicates cluster of neurons that are classified as GABAergic yet do not express *GAD1* and do express *SLC17A6*. **(C)** snRNA-seq from mouse amygdala neurons. Heatmap shows normalized, scaled expression of neurotransmitters in neurons from snRNA-seq of adult mouse amygdala (n = 25,930 nuclei, 3 replicates pooled). Dendrogram shows hierarchical clustering of neuron types. Dotted outline shows presence of 3 *Slc17a6*+ subtypes that preferentially cluster with GABAergic (*GAD1+/GAD2+*) subtypes and that express *Sim1* and/or *Adcyap1*. **(D)** Cartoon of embryonic migration of *Otp*+ cells that migrate from proliferative zones around the 3rd ventricle to periventricular hypothalamus (vzPZH) and medial amygdala (MeA), following data in Garcia-Moreno et al. (*45*). **(E)** ISH for *Otp* in sagittal section of P28 mouse brain. Dotted outline indicates borders of medial amygdala (MeA). **(F)** Marmoset P0 coronal section showing ISH staining for *OTP*. Data from (https://gene-atlas.brainminds.jp/; (Shimogori et al. 2018; Kita et al. 2021)). **(G)** Cartoon of marmoset amygdala the sagittal plane. FISH staining for *GAD1* (magenta), *SLC17A6* (cyan), and *SIM1* (yellow) in the medial amygdala (Me). Magenta arrows highlight *GAD1* expressing nuclei. Green arrows highlight *SLC17A6* and *SIM1* dual expressing nuclei. **(H)** Marmoset neuronal dendrogram shown in Fig. 1C with clades colored by *TAC3* expression. Unexpectedly, a novel primate-specific striatal *TAC3* population (*3*) clusters with hypothalamic and basal forebrain populations, and not with other telencephalic GABAergic interneurons. Red arrowhead indicates clade containing three *TAC3+* populations found in striatum, basal forebrain, and hypothalamus. **(I)** Heatmap of genes expressed in the clade indicated by red arrowhead in (H). Expression values of each gene is normalized to its max within the clade. The three *TAC3*+ populations are labeled and each show distinct expression differences.

The “cryptic” amygdalar subtypes display additional atypical gene expression features compared to other amygdala neuronal types, including expression of *OTP* and *SIM1* **(Fig. 3B)**. These transcription factors are typically expressed in neuronal lineages originating from proliferative zones around the third ventricle. In mice, there is a migratory stream of diencephalic neurons into the telencephalon (*45, 48*) into the medial amygdala, and this migration is dependent on *Otp* expression (*45*) (**Fig. 3D**). We generated mouse amygdala snRNA-seq data from 53,745 nuclei and verified the presence of a *Sim1*+ neuron type in mice that also clusters with GABAergic neurons and expresses *Slc17a6* but not *Gad1* or *Gad2* (**Fig. 3C**). In mice, these neurons constitute the majority population in the amygdala that express *Adcyap1* (encoding the protein PACAP) (**Fig. 3C**), a neuropeptide that is extensively (but not exclusively) expressed in hypothalamic populations (*49*) and associated with energy homeostasis (*50*), stress, anxiety (*51*), and immune responses (*52*). (In marmosets, *ADCYAP1* is additionally expressed in subsets of *SLC17A7*+ neurons that do not share the “cryptic” phenotype, indicating that *ADCYAP1* has a distinct distribution of expression in amygdala neurons between mice and marmosets.)

We further confirmed the expression of *Sim1+* cells in the medial amygdala in mice using the Allen Institute ISH atlas (**Fig. 3E**). To precisely locate the specific amygdalar nuclei housing the cryptic population, we investigated the expression of *SIM1* in neonatal marmosets (*53, 54*). *SIM1* expression was highly enriched in the neonatal marmoset medial amygdala (MeA) (https://gene-atlas.brainminds.jp; **Fig. 3F**), which mirrors the migration of *Sim1*+ neurons from the diencephalon in mice (28). We verified *SIM1* and *SLC17A6* but not *SIM1* and *GAD1* colocalization in adult marmoset medial amygdala using single-molecule fluorescence *in situ* hybridization (smFISH) (**Fig. 3G**). These results suggest that the cryptic amygdala neurons are a conserved population in both mice and primates that likely have diencephalic origins. Consistent with a “birthplace imprinting” effect, they retain transcriptional identities more similar to diencephalic types than to the telencephalic types with which they ultimately reside (**Fig. 3A**).

### Primate-specific striatal TAC3+ interneurons are similar to specific TAC3+ diencephalic types

Previously, we discovered a *TAC3+, LHX6*+ interneuron subtype in the striatum of humans, macaques, and marmosets that was absent in mice and ferrets (*3*). Compared with other striatal types, they are transcriptionally most similar to *PVALB*+ interneurons and, because they expressed transcription factor *LHX6*, we surmised that they likely also arose from the medial ganglion eminences (MGE) (*3*). Consistent with this inference, a recent study of fetal macaque single nucleus RNA-seq data from ventral telencephalic progenitor domains found that *TAC3*+ striatal interneurons likely arise from progenitors in the MGE, and diverge from an ancestral progenitor class that also gives rise to conserved *MAF*+ progenitors that produce *PVALB*+ and *TH*+ striatal interneurons (*33*).

The broader census of brain structures in the current dataset allowed us to compare the transcriptional identity of the striatal *TAC3*+ type to cell types outside of the striatum. Beyond expression in the striatal type, *TAC3* is expressed in 20 different neuron types in our dataset, including expression in cortical GABAergic neurons as well as several amygdala, basal forebrain, thalamic, and hippocampal types (**Fig. 3H; Fig. S3, Fig 6A,B**). *TAC3*+ types did not form a single clade in the dendrogram, and their transcriptional resemblance to other neuron types largely reflected their tissue or cephalic origin. For example, thalamic *TAC3*+ subtypes were found in a thalamic-only clade, while hippocampal *TAC3*+ neurons were most similar to other hippocampal and amygdala types. The thalamic types were *SLC17A6*+ while all other *TAC3*+ types were GABAergic.

Unexpectedly, the striatal *TAC3*+ type was most similar not to other striatal interneuron types as we previously concluded (*3*), but rather to two other *TAC3*+ GABAergic types found in basal forebrain and hypothalamus **(Fig. 3H**, tissue validation in **Fig. S6A**, **Data S2**). The broader clade containing these three types (depicted by green arrowhead in **Fig. 3H**) consisted entirely of basal forebrain and hypothalamic types, with the exception of the striatal *TAC3*+ type. Each of the similar *TAC3*+ populations had distinct gene expression (such as high expression of OXT and AVP in the hypothalamic type, and DRD2 expression exclusively in the striatal type), ruling out dissection contamination **(Fig. 3I)**. The 3 types remained direct neighbors when the dendrogram was recomputed using other distance metrics (e.g. correlation-based or PCA scores using transcription factor expression, see **Fig. S2**). As this clade assignment was unexpected, we omitted the basal forebrain and hypothalamic *TAC3*+ types and recomputed the dendrogram (retaining 286 of the original 288 neuronal types) to determine whether the striatal *TAC3*+ was broadly more similar to hypothalamic types than to telencephalic types **(Fig. 6C**). As striatal interneurons are generally divergent from hypothalamic/basal forebrain types, omitting these two cell types enabled us to test whether the *TAC3*+ striatal type is broadly more similar to hypothalamic neurons or to striatal neurons. The global structure of the dendrogram was essentially the same as the original, except that the striatal *TAC3*+ type now joined the major telencephalic GABAergic clade with other striatal interneurons (neighboring the striatal *PVALB*+ subtype) (**Fig. 6C**), recapitulating our original similarity assignment (*3*).

Considering their unexpected transcriptional similarity to both a telencephalic (basal forebrain) and a diencephalic (hypothalamus) type, the *TAC3*+ striatal type may either arise from a telencephalic progenitor (*33*) that also gives rise to sister diencephalic (hypothalamic) types, or else shows striking transcriptional convergence with diencephalic types that have distinct developmental origins. Favoring a telencephalic origin as suggested by previous report (*33*), the hypothalamic, basal forebrain and striatal *TAC3*+ types are all *FOXG1*+, a transcription factor associated with telencephalic origin (**Fig 3I**). They also express *LHX6*+ and *NKX2-1*+ (**Fig S3**), consistent with a medial ganglionic eminence origin, however we note that in mice some hypothalamic types also express *Nkx2-1* and *Lhx6* (*9, 55, 56*). Ultimately, lineage tracing of the striatal *TAC3*+ type in a primate would resolve whether a shared progenitor gives rise to both telencephalic and diencephalic types.

### Quantitative maps of interneuron distribution across primate neocortex and striatum

Our analysis of RNA expression in the single nucleus dataset indicates that developmentally-linked telencephalic GABAergic populations (*42, 43*) retain globally similar identities in adulthood. For example, *SST+* striatal interneurons are more similar to *SST*+ hippocampal and neocortical interneurons than they are to other striatal types (**Fig. S3**). Within each brain structure, developmentally linked populations become functionally specialized and follow distinct spatial rules for their allocation. In the mouse, quantitative imaging of molecularly identified interneuron types has revealed that densities of *Sst*+, *Vip*+, and *Pvalb*+ cortical interneurons vary across the cortical mantle; their relative proportions relate to unique functional and microcircuit properties of different cortical areas (*15*). In primates, overall neuron densities are more variable by cortical location than in mice: they vary by as much as 5 fold across the cortical sheet, with highest neuron proportions and densities found in occipital cortex and particularly in V1 (*57*). However, quantitative mapping of molecularly-defined subtypes of neurons have not been performed in a primate, and it is not known if they have conserved or distinct spatial distributions compared to mice.

We used smFISH to image the major cortical and striatal interneuron types across sagittal sections in marmoset (**Fig. 4** and **Fig. S7-9**). We quantified proportions and densities of each type relative to all cells (DAPI; **Fig. S7**). In the neocortex, we binned these using an areal parcellation of marmoset neocortex (https://doi.org/10.24475/bma.4520) to determine whether interneuron proportions varied systematically by brain area. In total, we quantified 377,554 neocortical interneurons across 30 sections by smFISH. In striatum, we quantified 6,848 interneurons across 32 sections. Each series sampled sagittal sections ∼160 µm apart, beginning 1,184-1,584 µm from the midline up to 6,384 µm laterally, including the majority of the striatum with the exception of the most-lateral portion of the putamen. We used a cell segmentation algorithm to count positive cells across sagittal sections and expressed interneuron proportions as a percent of all cells (DAPI+).

**Figure 4.**
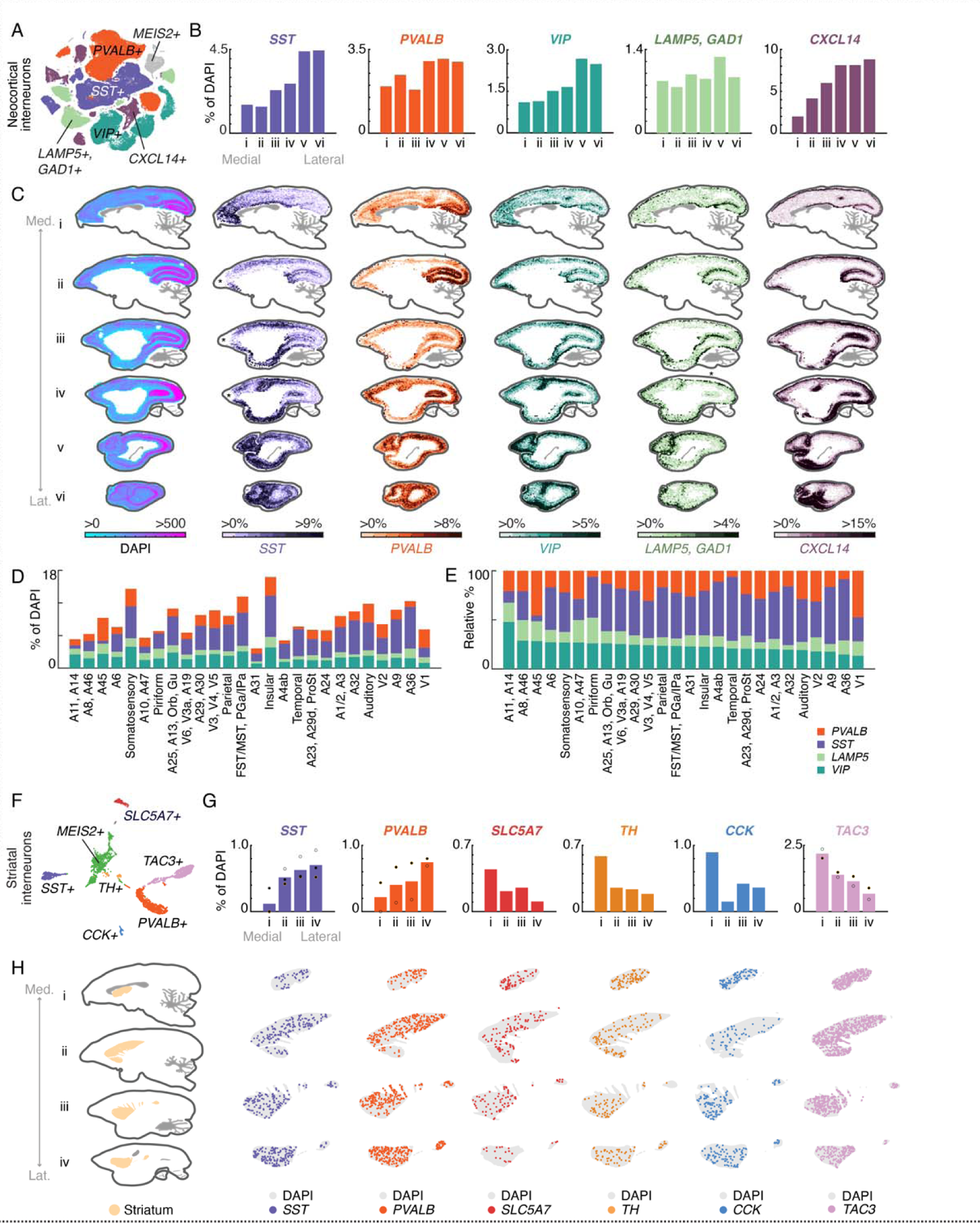
Cell-type-specific distributions of interneurons in primate telencephalon using quantitative smFISH. **(A)** t-SNE of GABAergic neocortical interneurons as in Fig. 2B, colored by subclass marker (*PVALB, SST, VIP, LAMP5, CXCL14*). Gray points are the *MEIS2+* population that is restricted to orbiomedial prefrontal cortex and was not spatially profiled. **(B)** Medial-lateral proportions of each major class as percentage of all cells (DAPI+). Barplots quantify positive cells as proportion of all (DAPI+) cells from medial to lateral sections of smFISH performed on 6 thin (16 µm) sagittal sections of marmoset neocortex, each section being 1600 µm apart and covering 9600 µm of neocortex. Colors as in (A). **(C)** smFISH for neocortical interneuron subclass markers showing locations of cells positive for each marker across 6 sagittal sections of the marmoset neocortex. First column shows density of all DAPI+ nuclei per unit area (approximately 387 µm per bin) profiled from one series. Heatmap scale in subsequent columns show percentage of marker-positive cells relative to DAPI+ cells. Average proportions across section shown in (B). Med. = medial, Lat. = Lateral. Asterisks in section ii and iii of the SST series and iv of the LAMP5 series denote regions where tissue or staining artifacts caused loss of signal. These can be seen in the DAPI-only series of these experiments, which show lower overall cell counts at these locations (**Fig S7**). **(D)** Quantitation of interneuron proportions by cortical area parcellated according to **Fig. S8B-C**. **(E)** Relative percentages of interneuron proportions by cortical area shown in Fig. S8B-C, sorted by max relative proportion of *VIP*+ interneurons. **(F)** t-SNE of striatal, cholinergic neurons (*SLC5A7*) and GABAergic interneurons (*SST*, *PVALB*, *TH*, *CCK*, *TAC3*). Green points correspond to the *MEIS2*+ population and were not spatially profiled. **(G)** Medial-lateral gradients of striatal cell type proportions across 4 sagittal sections. Dots = individual replicates. Colors as in (F). **(H)** smFISH for striatal cholinergic and interneuron subclass markers showing locations of cells positive for each subclass marker across 4 (16 µm) sagittal sections of the marmoset striatum, each 1600 µm apart covering 6400 µm in total. First column shows the anatomical context of the four sagittal striatal planes. Average proportions across each section are shown in (G).

Neocortical types were identified with probes for *SST, PVALB, CXCL14, VIP,* and *LAMP5* (**Table S4**), which collectively account for all major cortical interneuron populations **(Fig. 4A**). *CXCL14* is a marker for caudal ganglionic eminence derived cortical interneurons. It is expressed in most *VIP*+ and *LAMP5*+ cortical neurons, as well as a smaller population of *VIP*-, *LAMP5*-types, some of which are *PAX6*+ (and which correspond to the *SNCG*+ population in humans and mice (*2, 3*))). *VIP+* and *LAMP5+* interneurons are the two other major CGE-derived populations present in primates and mice. As *LAMP5* is also expressed in subsets of excitatory neurons, we performed dual labeling smFISH with *GAD1* to avoid counting glutamatergic types. Major striatal interneuron types were identified with probes for *SST, PVALB, CHAT/SLC5A7, TH, CCK,* and *TAC3* (**Table S4**). These together account for most major populations of non-SPN neurons in the striatum, with the exception of a population of *MEIS2*+ GABAergic striatal neurons that cluster together with non-SPN GABAergic neurons (**Fig. 1C, Fig. S3**) but are difficult to distinguish uniquely as several other markers are also expressed at variable levels in other striatal cell types.

### Neocortical interneuron types have highly focal biodistributions

Quantitative analysis of smFISH revealed highly focal and variable distributions of interneuron subtypes across marmoset neocortex (**Fig. 4B-E, Fig. S8A-D**). In both absolute numbers and relative proportions, *PVALB*+ interneurons were strongly enriched in the occipital lobe, particularly along the calcarine sulcus in the medial sections, as well as the occipital pole more laterally (**Fig. 4B-C; Fig. S8A, Data S1**). *SST+* interneurons in the neocortex increase medio-laterally (**Fig. 4B-C, Fig. S8A, Data S1**), but closer inspection revealed this is driven not by a spatial gradient so much as by highly focal enrichments around primary motor area (M1) and primary somatosensory cortex (S3, S1/2), the cingulate cortex, entorhinal cortex and medial prefrontal cortex (**Fig. 4C**). *CXCL14+* neurons are enriched along the calcarine sulcus medially, as well as in ventral aspects of the occipital cortex more laterally. There are higher proportions dorsomedially in the parietal cortex (**Fig. 4C**). *VIP*+ neurons were enriched in PFC and also increased laterally at or near somatosensory cortex and posterior parietal cortex (**Fig. 4C)**. *LAMP5+* interneurons showed a bias to the top of the cortical layers, consistent with dominant composition of this class as neurogliaform Layer 1 types (*3, 58, 59*), though this class also contains the *LAMP5/LHX6* type that is found in deeper layers (*3, 13*) (**Fig. 4C)**.

To better appreciate how these distributions relate to neocortical areas, we used a histologically-based marmoset neocortical parcellation (https://doi.org/10.24475/bma.4520; **Fig. S8B-C**) to bin smFISH interneuron proportions by cortical area (**Fig. 4D-E; Fig. S8B**). Small adjacent areas were merged, resulting in 26 areal groupings (**Fig. S8C**). Overall interneuron proportions relative to all cells varied by 4.5-fold across areal groupings (**Fig. 4D-E, Fig. S8B-C)**. As a fraction of all cells, A31 (dorsal posterior cingulate) had the lowest overall proportion of interneurons (3%), while insular cortex (In) had the highest (17%) (**Fig. 4D**).

In mice, quantitative mapping of interneuron densities showed higher *Sst*+ densities and lower *Pvalb*+ densities in neocortical areas involved in higher cognitive functions, such as medial frontal and lateral association areas (*15*). This local circuit feature follows cortico-cortical connectivity network topography in mouse (*60*). In contrast, lower *Sst/Pvalb* ratios were associated with mouse primary motor and sensory areas, which are associated with less distributed (and more local) cortical connectivity (*15*). To determine whether primate neocortex followed similar rules as mouse neocortex of interneuron allocation, we examined normalized proportions of the four largely mutually exclusive interneuron classes (*PVALB+, SST+, LAMP5+, VIP+*) within our marmoset areal groupings (**Fig. 4D-E; Fig. S8B**). Lateral temporal cortex, including A36, had the highest *SST/PVALB* ratio. Other areas with high *SST/PVALB* ratios included piriform cortex (Pir), M1 and several medial/orbitofrontal areas (A25, A13, A32). This suggests that some marmoset association areas, notably medial frontal area and lateral temporal cortex, have high *SST/PVALB* ratio composition consistent with high *Sst/Pvalb* ratios in higher order association network in mice (*15*) (**Fig S8B**).

Strikingly, polar and lateral prefrontal areas (including A8, A46, A10, A47, A45), which are thought to be the most divergent relative to prefrontal areas in rodents (*61*), and which in primates are characterized by long-range cortico-cortical connectivity to other association areas (*62–64*), do not exhibit high *SST/PVALB* ratios (**Fig. 4D-E; Fig. S8B**). Of all cortical areal groupings we measured, the lowest *SST/PVALB* ratio was found in A45, a higher-order lateral prefrontal area with extensive projections to all lobes of the neocortex (*64*); http://analysis.marmosetbrain.org/). Thus, unlike the high *Sst/Pvalb* ratios observed in frontal areas in mouse (*15*), primate lateral prefrontal areas are characterized not by exceptional *SST/PVALB* ratios but rather as having some of the highest *VIP*+ interneuron proportions (**Fig. 4D-E; Fig. S8B**). Lateral prefrontal areas also have the highest total fraction of *VIP*+ and *LAMP5/GAD1+* interneurons (all above 50% of all interneurons); both of these populations predominately arise from the caudal ganglionic eminence, a progenitor zone that has expanded in primate evolution (*65*). These results suggest that primate lateral prefrontal areas do not follow the same local cortical circuit organizing principles that typify frontal areas in the mouse (*15*), or medial frontal areas in marmoset (**Fig. 4D-E; Fig. S8B**).

### Interneuron proportions in marmoset striatum follow medio-lateral gradients

The striatum, a crucial brain region involved in motor control, reward, and decision-making, displays complex functional topography and connectivity. In humans, anterio-medial portions containing nucleus accumbens/ventral striatum are functionally coupled to limbic and higher order cognitive networks, while lateral subdivisions are more coupled to sensory and motor networks (*66*). Prior work relating bulk gene expression measurements in human and macaque striatum to cortico-striatal network organization found a relationship between functional domain (e.g. somato/motor vs limbic network) and gene expression, including enrichment of *PVALB* in lateral portions of the striatum that are coupled to somatosensory and motor cortex (*67*). These results suggest that differences in local cell type composition across striatum may underlie aspects of its functional topography.

In mice, striatal interneuron subtypes have different spatial distributions: cholinergic (*Chat*+) neuron proportions increase dorsally and anteriorly (*68*), *Pvalb*+ interneurons are more abundant in dorsolateral striatum than in dorsomedial striatum, and *Sst*+ interneurons are spatially homogeneous (*69*) (**Fig. S9A**). Primates retain the major populations of striatal interneurons found in mice (*6, 70*), and additionally have gained the novel type distinguished by *TAC3* expression (*3, 33*) described in previous sections (**Fig. 3H, 4F**). A systematic quantification of striatal interneuron types has not been performed comprehensively in a primate, and it is unknown if they follow similar or distinct spatial distributions observed in mice. We used single-molecule FISH (smFISH) to investigate distributions of the major types of conserved striatal interneurons (*SST*+, *PVALB*+, *SLC5A7+*/*CHAT*+, *TH*+, *CCK*+, *TAC3*+, probes in **Table S4**) in serial sagittal sections of marmoset striatum (**Fig. 4G-H**).

Each striatal interneuron population exhibited a non-uniform distribution across the marmoset striatum, particularly in the medial-lateral axis. Similar to mice, the proportion of striatal *PVALB+* interneurons increases in lateral sections, from ∼0% to 0.8% of all cells (**Fig. 4G, Data S1**). Unlike mice, marmoset *SST+* interneuron distribution is non-uniform, appearing sparse near the midline and increasing in proportion (0.1-0.7%; **Fig. 4G, Data S1**). Cholinergic neurons (*CHAT+*) show the opposite medial-lateral gradient (0.45%-0.1%; **Fig. 4G, Data S1**). Similar to *CHAT+* neurons, *TH+* striatal interneurons, which are transcriptionally similar to the *PVALB+* type, exhibited a decreasing medial-lateral gradient (**Fig. 4G, Data S1**). *CCK+* striatal interneurons, which are a minority population in marmoset (*3*) and mouse (*70*) are enriched close to the midline and become much sparser laterally (**Fig. 4G, Data S1**). The *TAC3+* interneurons showed an increasing medial-lateral gradient, similar to *CHAT+* neurons (2%-0.5%; **Fig. 4G, Data S1**). No striatal population exhibited anterior-posterior or dorsal-ventral gradients with the exception of *PVALB* interneurons, which showed a modest dorsal-ventral gradient. Unlike striatal interneurons, MSNs proportions were largely uniform across the major axes: *DRD1* and *DRD2*, which distinguish direct and indirect MSNs, respectively, were slightly enriched medially but otherwise exhibited largely uniform distributions across the striatum (**Fig. S9B**). Overall, these findings suggest that the spatial distribution of interneuron subtypes within the primate striatum may play a role in shaping its functional topography and connectivity. While mice and primates share similar striatal interneuron populations, there are also differences in their spatial distributions.

### Morphology of conserved neocortical and striatal interneuron types

The morphology of interneuron types relates essentially to their function and contributions to neural circuits. While methods such as biocytin filling and Golgi staining are the gold standard for morphological reconstructions, the administration of low-titer AAVs carrying membrane-bound fluorescent reporters can be used to sparsely label cells for morphological reconstruction. We utilized a reporter AAV under the control of the forebrain interneuron-specific mDlx enhancer (*19*) to label neocortical and striatal interneurons, and then performed smFISH (probes in **Table S4**) on thick sections (120 µm) to confirm the molecular identity of GFP+ cells. Reconstructions using viral labeling are more challenging than with single cell filling methods, because GFP expression from neighboring cells and passing fibers have to be distinguished from signal attributable to the target cell. In some cases, the GFP signal appears punctate, making it challenging to follow discontinuous processes. For these reasons, our reconstructions are conservative: as we aimed to avoid reconstructing false positive fibers (fibers originating from other cells), we may in some cases under-ascertain the full dendritic arborization of target cells.

We imaged 1,203 GFP+ neurons in the striatum and 4,374 GFP+ neurons in the neocortex. We used NeuTube to reconstruct the top 216 telencephalic neurons that had the best GFP signal, did not have other cells labeled in the field of view, and were positive for at least one smFISH probe **(Table S5; examples in Fig. S10)**. Raw image stacks and NeuTube reconstructions are available at https://doi.org/10.35077/g.609. Using combinations of 1-2 probes for marker genes of different types, we identified GFP+ neocortical and striatal interneurons, respectively, with smFISH based on their type markers (neocortex: *SST*, *PVALB*, *CXCL14*, *VIP*, *LAMP5*; striatum: *SST*, *PVALB*, *SLC5A7*, *TH*, *CCK*, **Table S4**) (**Fig. 5A-B, *top rows***). We used a second tracing and reconstruction method (Imaris) on the 41 most complete GFP+ cells. Of these, we discarded 5 due to incomplete soma or highly discontinuous processes in the image stack, retaining 36 cells (**Fig. 5A-B, *bottom rows*; Table S6**). Morphological parameters were measured using the Surface function, which detects surface area and volume based on the fluorescence of the mDlx-AAV-GFP expression, and the Filament Tracer function, which traces structural features starting from the soma to the terminal processes based on the diameter of the soma and the thinnest projection of the cell **(Table S6)**. To assess whether there are region-dependent morphological differences within molecularly similar types, we compared several parameters between a collection of neocortical and striatal *PVALB*+ GFP+ cells that were reconstructed from the same marmoset (Cj 17-154; **Fig. 5C-D**) to avoid variable tissue shrinkage arising from different tissue storage conditions across the animals (see *Methods*). Striatal *PVALB*+ GFP+ cells were larger than cortical *PVALB*+ GFP+ cells in terms of length, area, but not soma diameter or the volume of GFP+ fluorescence (**Fig. 5D**). While the number of branches issued by their somas also did not differ, the striatal *PVALB*+ GFP+ interneurons exhibited more dendritic branch points than their cortical counterparts (**Fig. 5D**), suggesting that striatal *PVALB*+ interneurons have greater arborization than *PVALB*+ cortical interneurons.

**Figure 5.**
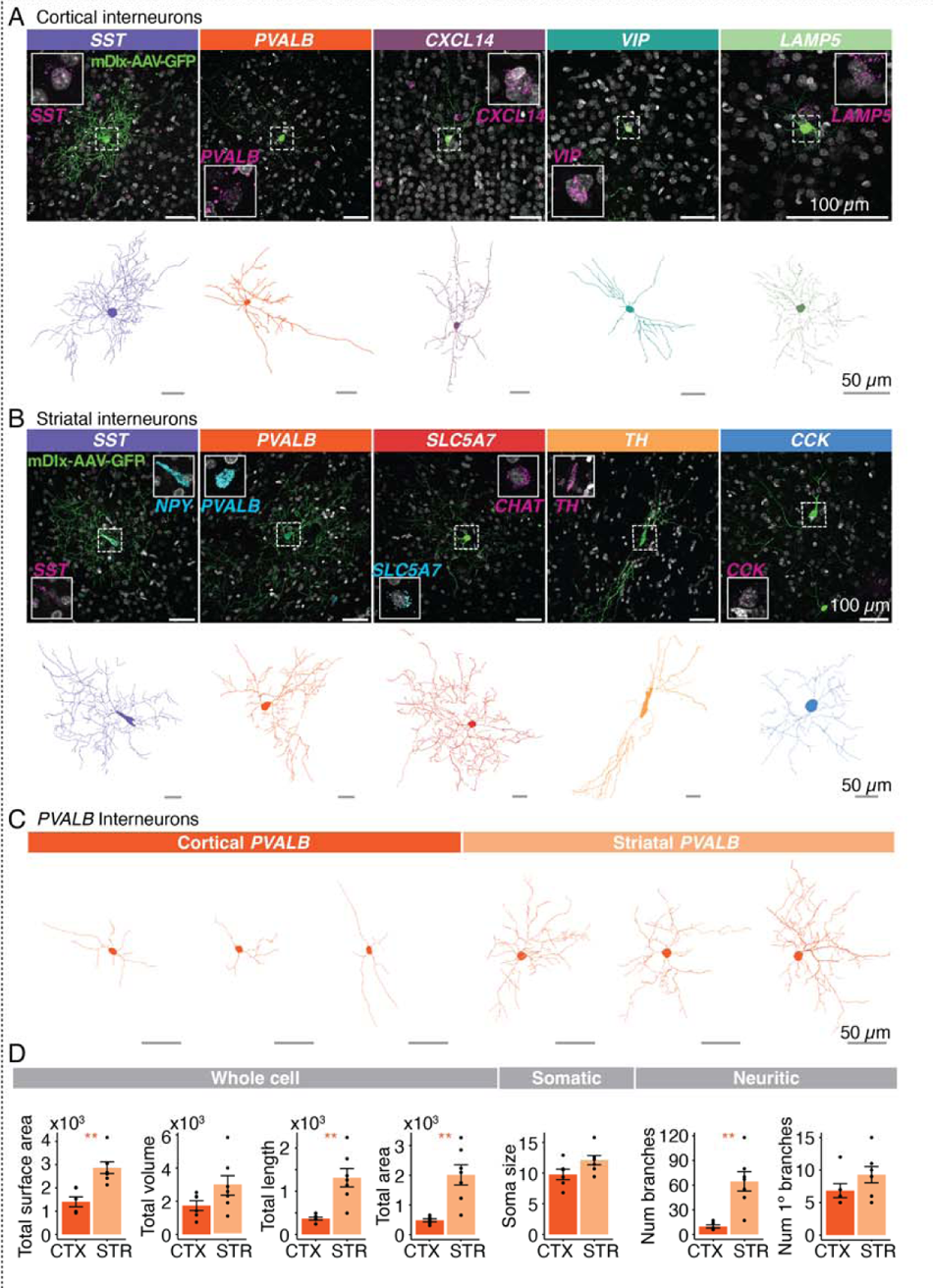
Morphological characteristics of marmoset cortical and striatal interneurons. **(A-B)** *Top rows*, Examples of AAV9-hDlx5/6-GFP-fGFP labeled neocortical **(A)** and striatal **(B)** interneurons. Insets show magnified cell nucleus of GFP+ cell along with smFISH staining for interneuron type marker to confirm molecular identity of labeled cell. Scale Bar = 100 µm. *Bottom rows,* Reconstructed skeletonized morphology (Imaris) of GFP+ cells depicted in *top rows*. Scale Bar = 50 µm. **(C)** Representative reconstructed neocortical and striatal *PVALB+* interneurons from one marmoset (Cj 17-154; **Table S1, S6**). Scale Bar = 50 µm. **(D)** Quantification of morphological characteristics of *PVALB*+ cells from Cj 17-154. Means ± SEM. \**P* < 0.05 and \*\**P* < 0.01. See the Methods section for detailed statistical information.

### Development of a novel TAC3-rAAV-GFP reporter

Given that the *TAC3*+ type comprises almost 30% of striatal interneurons in marmoset, we expected a sizable proportion of striatal GFP+ cells labeled by AAV9-hDlx5/6-GFP-fGFP (**Fig. 5**) to be *TAC3*+. However, while 55 GFP+ striatal cells were imaged across all smFISH *TAC3*-probe treated sections, we failed to find any colocalization with *TAC3* transcripts. In these experiments, which were double-labeled with *TAC3* and *PVALB* probes, 9/55 were *PVALB*+ (16% *PVALB*+, vs expected 21% of interneurons expected from snRNA-seq proportions), while 0/55 were *TAC3*+ (0% vs 28% expected from snRNA-seq).

The mDlx enhancer is a regulatory element specific to forebrain interneurons (*71*). Though the regulatory element, and the forebrain-interneuron expressed genes flanking it, *Dlx5* and *Dlx6*, are highly conserved in vertebrates, we wondered whether the lack of accessibility of the mDlx locus in the *TAC3* interneurons could explain our inability to find colocalization of *TAC3* expression and GFP. To assess this, we generated single nucleus ATAC-seq (snATAC-seq) data (69,808 nuclei) from fresh marmoset striatum (**Fig. 6A**). We used Signac (*72*) to integrate our previously annotated striatal snRNA-seq data and identify major striatal types. We then examined accessibility of the marmoset sequence homologous to the mDlx locus across interneuron types. While other striatal interneuron types (particularly the *SST*+ and *PVALB*+ types) showed accessibility (ATAC-seq peaks, reflecting chromatin accessibility) at the mDlx locus, the *TAC3*+ type did not (**Fig. 6B**).

**Figure 6.**
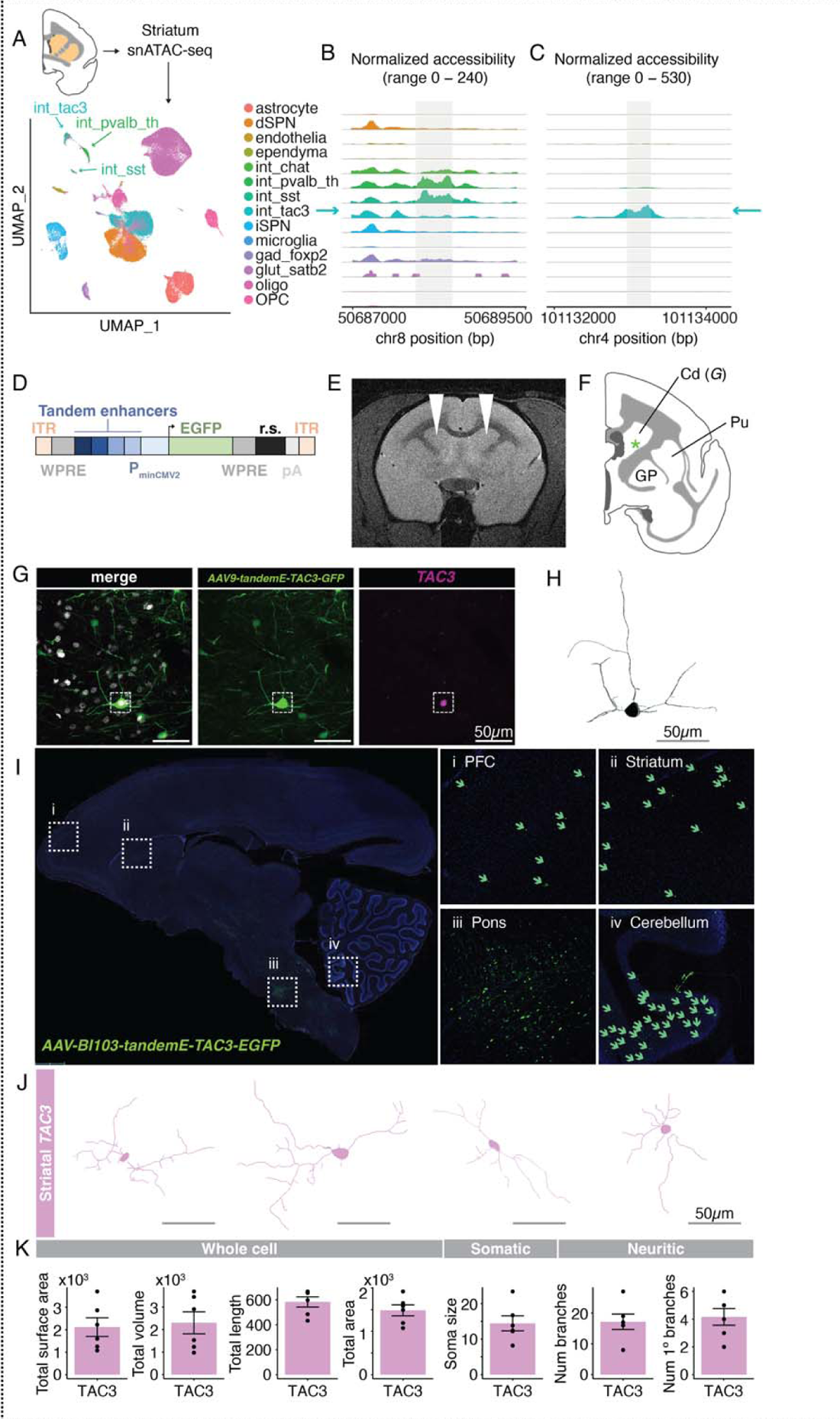
Development of a novel enhancer-AAV for *TAC3*+ primate-specific striatal interneurons. **(A)** snATAC-seq (69,808 nuclei) from fresh marmoset striatum (1 male, Cj 18-153). UMAP shows major clusters with labels transferred from striatal snRNA-seq data. **(B)** Chromatin accessibility at the locus corresponding to the mDlx sequence in marmoset shows read pileups in *PVALB*, *SST*, and *CHAT*+ neurons, but not in the *TAC3*+ type. **(C)** Chromatin accessibility of a candidate selective enhancer for the *TAC3*+ type. **(D)** Sequence construct design for *TAC3* interneuron specific AAV. Four *TAC3* interneuron-specific regulatory elements (example in (C)) are packaged in tandem upstream of a minimal promoter driving EGFP expression. ITR = inverted terminal repeats. WPRE = Woodchuck Hepatitis Virus Posttranscriptional Regulatory Element. PminCMV2 = minimal CMV2 promoter. EGFP = enhanced green fluorescent protein. R.S. = 300 bp random sequence. pA = polyadenylation sequence. **(E)** MRI showing injection location (white arrowheads) of the AAV9-tandemE-TAC3-EGFP virus into bilateral dorsal striatum (caudate) in one animal (Cj 19-207, **Table S1**). **(F)** Cartoon showing location of cell shown in (G). **(G)** Main: EGFP antibody-amplified confocal image of a labeled cell (position shown in (F)). Insets: smFISH for *TAC3* showing colocalization. Scale bar = 50 µm. **(H)** Morphological reconstruction (Imaris) of cell shown in (G). Scale bar = 50 µm. (**I**) Whole sagittal section (20x image) of adult marmoset showing cells transduced by the AAV-BI103-tandemE-TAC3-EGFP virus in marmoset Cj 20-214. GFP+/*TAC3*+ cells were detected sparsely in neocortex as well as striatum, cerebellum, substantia nigra, superior colliculus, and brainstem. (*i*) Prefrontal cortex, (*ii*) Striatum, (*iii*) Pons, (*iv*) Cerebellum. Green arrows highlight cells in areas with sparse expression. **(J)** Four reconstructions (Imaris) of striatal GFP+/*TAC3+* cells from similar section shown in (I). **(K)** Quantification of morphological parameters of reconstructed cells from striatal GFP+/*TAC3+* cells from the local injection in (E) as well as the systemic injection in (I) (**Table S6**).

To develop a viral tool that could transduce the striatal *TAC3+* cell type, we next nominated new candidate regulatory elements specific to the *TAC3*+ interneuron type. We used Signac to identify differentially accessible peaks in the *TAC3*+ cluster relative to all others, and filtered the set by fold change, percent accessibility across the target cell type population, and peak size (**Fig. 6C**). To maximize the likelihood of obtaining a functional reporter while minimizing the number of marmosets used, and because several of our top candidates were very short, we selected four top regulatory element candidates for the *TAC3*+ type for tandem packaging (**see *Methods***). The four candidates were on four different chromosomes and spanned between 94-215 bp. One site was in an exon of *CDH13*, one was in the first exon of *TAC3*, one was in an exon of *LOC108592287*, and one was intergenic (closest gene 200kb distance). These four elements were packaged in tandem in an AAV9 vector (AAV9-tandemE-TAC3-EGFP) containing a cytoplasmic GFP reporter (**Fig. 6D**). We delivered the virus via MRI-guided local injection into the anterior striatum of two marmosets (Cj 19-207 and 17-B111, **Fig. 6E and Fig. S11A, Table S1**), and imaged coronal sections of striatum for GFP positive cells after 6 & 10 weeks’ incubation time (**Fig. 6F and S11C**). smFISH confirmed colocalization of *TAC3* transcripts in GFP+ neurons in striatum (**Fig. 6G and Fig. S11B,D**). In one animal the virus diffused beyond the boundary of the striatum. In this animal we also detected strongly labeled GFP+ cells in a border zone between striatum, BNST, and the globus pallidus where the viral injection had diffused beyond striatum (**Fig. S11B-E**), and smFISH confirmed that these too were *TAC3+*.

To determine the broader biodistribution of cells transduced by the tandemE-TAC3 enhancer, we next packaged the same enhancers in a novel AAV capsid (BI103) capable of efficient transduction of brain cell types after systemic IV delivery in marmosets (Chan et al, in prep). We delivered this AAV (AAV-BI103-tandemE-TAC3-EGFP) to one marmoset (Cj 20-214, **Table S1**). After a 4 week incubation, the brain was perfused, sliced and stained with anti-GFP antibody to amplify GFP signal, and smFISH probes against *TAC3*+ to confirm colocalization. We found GFP+/*TAC3*+ colocalized cells in striatum as well as several extra-striatal locations including hypothalamus, substantia nigra, superior colliculus, brainstem, and neocortex (**Fig. 6I**). To assess the morphology of *TAC3*+ striatal interneurons produced by viral labeling, we reconstructed several of the most complete cells from both the local injections (AAV9-tandemE-TAC3-EGFP) and from the systemic injection (AAV-BI103-tandemE-TAC3-EGFP) using Imaris (**Figs. 6H,J,K** and **Table S6**). The cells tended to have 2-3 thick branches that extended from the soma, and which bifurcated close to the soma and became thinner thereafter. Reconstructed cells had median 13.1 µm (+/- 5.1 s.t.d) soma diameter, 2221.3 volume (µm^3; +/- 1192.8 s.t.d) and 16.5 (+/- 6.11) total dendritic branch points. These experiments show that whereas systemic injections of AAV9 under the mDlx enhancer could not transduce the striatal *TAC3*+ type, presumably due to loss of accessibility at the mDlx locus (**Fig. 6B**), local injections of AAV9 as well as systemic injections with alternative capsid with the cell type targeted enhancers were successful. Broadly, the use of cell type specific regulatory elements, coupled with viral engineering, enables a new horizon for the study of primate brain cell types.

## Discussion

In this study, we generated a molecular and cellular census of the adult marmoset brain using single-nucleus RNA sequencing, quantitative smFISH, and cell type specific AAV labeling. Our study reveals the complex repertoire of cell types in the marmoset brain. Our snRNA-seq dataset of over 2.4 million brain cells across 18 brain regions in the marmoset indicates that lineage is an important factor shaping adult transcriptomic identity of neuronal types, apparently more so than neurotransmitter utilization.

Using quantitative smFISH we revealed, for the first time in a primate, the spatial distributions of molecularly-resolved GABAergic interneuron types. Using GFP delivered by interneuron-specific AAVs, we generated morphological reconstructions of all major interneuron types in the neocortex and striatum. We generated a novel viral genetic tool under the control of cell type specific regulatory elements to transduce a previously described putative primate-specific striatal interneuron type. These datasets, generated as part of the BRAIN Inititive Cell Census Network, complement other recent and extensive cellular profiling studies in other species (*8–11, 73*) and will enable insights in cell type innovations and modifications in future comparative studies.

Telencephalic glutamatergic and GABAergic neurons strongly segregate in mammals as well as in homologous structures in amphibians (*26*), suggesting an evolutionarily conserved distinction. An initial atlas in mouse indicated that glutamatergic and GABAergic neurons from diencephalic and midbrain structures also partition almost perfectly by neurotransmitter usage (*7*), suggesting this could be a general rule. However, more recent transcriptomic censuses in mouse (*8, 9*) and human (*11*) indicate that across mammalian species, many non-telencephalic glutamatergic and GABAergic types tend to form highly intermixed clades that do not separate clearly by neurotransmitter identity. Our results in the adult marmoset (and mouse, **Fig S2**) concord with the notion that gene expression distinctions between telencephalic glutamatergic and GABAergic neurons do not hold for neurons in other brain structures (**Fig. 1; Fig. S2**). This has implications for generalizing transcriptomic associations to other phenotypes. For example, transcriptomic changes in glutamatergic or GABAergic neurons have been associated to diseases such as autism and schizophrenia (*74–76*). Such associations may not generalize to GABAergic or glutamatergic types outside of the sampled brain region (usually neocortex), consistent with observations that “global” changes to glutamatergic or GABAergic neurons in relation to disease actually often only surface in a few brain regions (*77*).

A previous analysis, based on shared patterns of several key transcription factors, proposed that telencephalic GABAergic neurons are developmentally and evolutionarily related to diencephalic GABAergic neurons (*78*). Our results indicate that when profiled at adulthood, only a limited number of telencephalic GABAergic types are transcriptionally similar to diencephalic types, some of which may arise from cephalic boundary crossings. Most neocortical, hippocampal, and some amygdalar and striatal GABAergic types are so distinct from diencephalic GABAergic types that they share more gene expression in common with telencephalic glutamatergic types (**Fig. 1C**).

Developmental origin or shared lineage plays a strong role in shaping the adult transcriptomic identity of neurons, but phenotypic convergence, whereby adult cell types converge on similar transcriptomic identities despite a non-shared developmental origin, may also drive apparent similarities amongst cell types. Recent advances in lineage tracing coupled with single cell RNA sequencing demonstrates that phenotypic convergence is surprisingly common between transcriptomically defined types (*5, 79*). As such, it is difficult to disambiguate between these possibilities in absence of data that confirm lineage directly (*5, 26*). For example, in our data the clade of GABAergic projection neurons that contains striatal SPNs and amygdala and basal forebrain *GAD1+, FOXP2+* neurons also contained several subtypes of hypothalamic *GAD1+, FOXP2+* neurons (**Fig. 1C; Fig. S2E, Fig. S3**). Either these hypothalamic *FOXP2+* subtypes have a convergent expression identity with long-range GABAergic projection neurons of the telencephalon, or else they arise from a common lineage.

Our data support known instances of cross-cephalic migrations in thalamus and amygdala, and also suggest new ones (**Fig. 3**). The similarity of the primate-specific *TAC3*+ striatal type to hypothalamic and basal forebrain *TAC3*+ types in particular is unexpected (**Fig. 3H**) and suggests a potential example of cross-cephalic vesicle migration. Recent work (*33*) proposes that the striatal *TAC3*+ type has a ventral ventricular telencephalic origin, similar to most other GABAergic interneuron types destined for striatum. While phenotypic convergence remains possible, another possibility is that a ventral telencephalic progenitor gives rise to both telencephalic and diencephalic types. This is supported by the expression of *FOXG1*, a transcription factor necessary for ventral telencephalic fate (*80*), in all three transcriptionally similar *TAC3*+ populations.

Developmentally linked interneuron populations in striatum and neocortex (e.g *PVALB*+ types in both structures) displayed distinct spatial distributions measured by cell counting of smFISH (**Fig. 4)**. While all interneuron subtype distributions in the striatum followed a gradient along a medial-lateral axis, the interneurons in the neocortex followed much more complex distributions that for the most part were not captured by simple gradients. While in the mouse, *Sst+/Pvalb+* ratios are a hallmark of higher order cortex (*15*), we found a different pattern of relative interneuron proportions across much of the higher-order association cortex in marmosets. Areas in the lateral prefrontal cortex were unique in their relatively high proportions of *VIP*+ neurons (and had unexceptional or even low ratios of *SST*+/*PVALB*+ neurons). These results suggest that primate lateral prefrontal areas may not follow the same local cortical circuit organizing principles of association cortex in the mouse.

Morphological characterization of the striatal and neocortical interneuron populations suggests variation in overall size and dendritic arborization amongst subtypes. To avoid interindividual and technical variability arising from different sample storage conditions (see Methods), we analyzed several morphological parameters in a subset of cortical and striatal *PVALB+* interneurons reconstructed from a single marmoset (**Figs. 5C-D**). Striatal *PVALB*+ cells were found to be larger in length, volume, and surface area and consisted of more dendritic branch points than the cortical cells. Our data altogether suggest that the striatal *PVALB+* interneurons exhibit higher dendritic complexity compared to their respective counterparts in the neocortex. These morphological differences could result in differing electrophysiological properties. For example, it is possible that *PVALB*+ cells in the striatum have higher capacitance and receive more synaptic inputs onto them, thereby impacting local signal integration and computation, than cortical *PVALB*+ interneurons. The functional significance of morphological differences across the interneuron subtypes identified in our study will ultimately require studying how these cells affect the circuits in which they reside (i.e., cellular/subcellular targeting biases and functional properties)(*81–83*).

The development of virally-based tools enables cell-type-specific access in nonhuman primates (*2, 18*– *20*). To maximize translatability across species, most approaches nominate candidates using evolutionary conservation (usually between mice and humans) as a selection criterion. However, the evolutionarily conserved mDlx enhancer did not drive expression in the novel striatal *TAC3+* type **(Fig. 6B)**. In general, we observed that the mDlx enhancer selectively under-ascertained several interneuron populations in marmoset in these experiments. For example, it systematically under-labeled *VIP*+ and *SST*+ types in the neocortex (expected 22% and 26% of interneurons, obtained 2% and 3%, respectively), as well as *SST*+ interneurons in the striatum (14%, obtained 1.8%). This underlabeling could in part be due to the titer and systemic delivery approach we adopted, which was necessary in order to achieve sparse labeling for morphology. The overall low efficiency in our ability to molecularly characterize GFP positive cells suggests a need for further optimization for this species and application.

The lack of transduction of *TAC3+* striatal interneurons using the AAV-mDlx virus prompted us to develop a novel AAV under the control of *TAC3*+ interneuron striatal regulatory elements (**Fig. 6D-H**). The novel reporter virus (AAV9-tandemE-TAC3-EGFP) that we developed to study the striatal *TAC3*+ type was driven by striatal *TAC3*+ enhancer elements. When delivered systemically, the virus also labeled *TAC3*+ neurons elsewhere in the brain, including in the hypothalamus, neocortex, substantia nigra. As neocortical *TAC3*+ cells bear little resemblance to *TAC3*+ striatal interneurons in terms of their global gene expression profiles (**Fig. 3H**), the widespread transduction of *TAC3*+ neurons suggests that the regulatory elements that we identified in the *TAC3*+ striatal type (defined only relative to other striatal cell types) may endogenously regulate the *TAC3* gene itself, or else a gene whose expression is highly correlated to *TAC3*.

Our study reveals the complex landscape of transcriptionally-defined cell types in the marmoset brain. Compared to mouse or other outgroup species, few primate lineage-gained cell types have emerged; likely more common is the redirection, repurposing, or elaboration of conserved types (*2, 13, 33, 84, 85*). The persistent fingerprint of developmental origin present in neuronal gene expression underscores important roles for novel neurogenic niches, developmental patterning and altered mechanisms guiding proliferation and cell migration in primate brain evolution that could be the subject of future study. For example, *in vivo* lineage tracing in primates could reveal the developmental origin of the striatal TAC3+ type and its relationship to other TAC3+ populations in other brain structures. We report the largest transcriptomic cell census of the marmoset brain and uncover the complex biodistribution of telencephalic interneurons by quantitative smFISH mapping. We present a compendium of morphological reconstructions of marmoset telencephalic interneurons, and describe new viral reagents for targeting and visualizing a primate-specific striatal interneuron type. Together, these resources will enable further comparative studies of the evolution and development of brain cell types.

## METHODS

### Animals used for study

#### Marmosets

Marmosets were pair-housed in spacious holding rooms with environmental control of temperature (23–28°C), humidity (40–72%), and 12 hr light/dark cycle. Their cages were equipped with a variety of perches and enrichment devices, and they received regular health checks and behavioral assessment from MIT DCM veterinary staff and researchers. All animal procedures were conducted with prior approval by the MIT Committee for Animal Care (CAC) and following veterinary guidelines.

#### Mice

Experimental mice were purchased from The Jackson Laboratory company and housed at the Mclean hospital animal facility (3–5 mice per cage) on a 12:12 hr light-dark cycle in a temperature-controlled colony room with unrestricted access to food and water. All procedures were conducted in accordance with policy guidelines set by the National Institutes of Health and were approved by the McLean Institutional Animal Care and Use Committee (IACUC).

### Tissue processing for single nucleus sequencing and smFISH

#### Marmoset specimens for snRNA-seq

Marmoset experiments were approved by and in accordance with Massachusetts Institute of Technology CAC protocol number 051705020. Adult marmosets (1.5–14.5 years old, 12 individuals; **Table S1**) were deeply sedated by intramuscular injection of ketamine (20–40 mg kg−1) or alfaxalone (5–10 mg kg−1), followed by intravenous injection of sodium pentobarbital (10– 30 mg kg−1). When the pedal with-drawal reflex was eliminated and/or the respiratory rate was diminished, animals were trans-cardially perfused with ice-cold sucrose-HEPES buffer (*3, 6*). Whole brains were rapidly extracted into fresh buffer on ice. Sixteen 2-mm coronal blocking cuts were rapidly made using a custom-designed marmoset brain matrix. Slabs were transferred to a dish with ice-cold dissection buffer (*3, 6*). All regions were dissected using a marmoset atlas as reference(*86*), and were snap-frozen in liquid nitrogen or dry ice-cooled isopentane, and stored in individual microcentrifuge tubes at −80 °C.

#### Marmoset specimens for snATAC-seq

Tissue from 1 marmoset (female, 1.5 y.o., **Table S1**) was used for both snRNA-seq and snATAC-seq (**Table S1**). Fresh tissue was dissected from anterior striatum (including caudate and putamen) and used immediately for snATAC-seq.

#### Mouse specimens for snRNA-seq

Three adult (P80-90) male wild-type mice were deeply anesthetized with isoflurane and sacrificed by decapitation. Brains were quickly excised, washed in ice-cold sterile 0.1M phosphate buffer saline (PBS) and dissected onto an ice-cold glass surface. Amygdala nuclei were identified and isolated using “The Allen mouse brain atlas” (https://mouse.brain-map.org/static/atlas, **Table S7**) as a reference for anatomical landmarks. The basolateral amygdaloid nucleus was exposed by performing two coronal cuts using the borders of *primary somatosensory cortex* and *primary visual cortex* as landmarks. Dissected specimens were collected in 1.5ml micro-centrifuge tubes, snap-frozen on dry ice, and stored at −80 °C until used.

#### Marmoset specimens for smFISH

Two marmosets were deeply sedated by intramuscular injection of alfaxalone (5–10 mg kg−1) (**Table S1**), followed by intravenous overdose of sodium pentobarbital (10– 30 mg kg−1). When the pedal with-drawal reflex was eliminated and/or the respiratory rate was diminished, animals were trans-cardially perfused with ice-cold saline. The brain was immediately removed, embedded in Optimal Cutting Temperature (OCT) freezing medium, and flash frozen in an isopropyl ethanol-dry ice bath. Samples were cut on a cryostat (Leica CM 1850) at a thickness of 16µm, adhered to SuperFrost Plus microscope slides (VWR, 48311-703), and stored at −80 °C until use. Portions of the brain that were not cut were recoated in OCT and stored again for future use. Samples were immediately fixed in 4% paraformaldehyde and stained on the slide according to the Molecular Instruments HCR generic sample in solution RNA-FISH protocol (Molecular Instruments, https://files.molecularinstruments.com/MI-Protocol-RNAFISH-GenericSolution-Rev7.pdf) or the Advanced Cell Diagnostics RNAscope Multiplex Fluorescent Reagent Kit v2 Assay (ACD, 323100, https://acdbio.com/sites/default/files/USM-323100%20Multiplex%20Fluorescent%20v2%20User%20Manual_10282019_0.pdf) protocol (**Table S7)**.

### Single nucleus RNA-seq, library preparation, sequencing

*10x RNA-seq.* Unsorted single-nucleus suspensions from frozen marmoset and mouse samples were generated as in (*87*). GEM generation and library preparation followed the manufacturer’s protocol (10X Chromium single-cell 3′ v.3, protocol version #CG000183_ ChromiumSingleCell3′_v3_UG_Rev-A).

Raw sequencing reads were aligned to the NCBI CJ1700 reference (marmoset) or GRCm38 (mouse). Reads that mapped to exons or introns were assigned to annotated genes. Libraries were sequenced to a median read depth of 8 reads per Unique Molecular Identifier (UMI, or transcript), obtaining a median 7,262 UMIs per cell.

### RNA sequencing data processing, curation, clustering

Processing and alignment steps follow those outlined in: https://github.com/broadinstitute/Drop-seq/ (**Table S7)**. Raw BCL files were processed using IlluminaBasecallsToSam, and reads with a barcode quality score below Q10 were discarded. Cell barcodes (CBCs) were filtered using the 10X CBC whitelist, followed by TSO and polyA trimming. Reads were aligned using STAR, and tagged with their gene mapping (exonic, intronic, or UTR) and function (strand). The reads were then processed through GATK BQSR, and tabulated in a digital gene expression matrix (DGE) containing all CBCs with at least 20 transcripts aligning to coding, UTR, or intronic regions. Cell selection was performed based on CellBender remove-background non-empties (*88*), % intronic (% of a CBC’s reads that are intronic), and number of UMIs for a CBC. A new filtered DGE containing only these selected CBCs was then generated. Finally, a gene-metagene DGE was created by merging the selected CBCs DGE with a metagene DGE (made by identifying reads from the selected CBCs that have a secondary alignment mapping to a different gene than its primary alignment).

Cell type classification models were trained using our annotations and scPred R package version 1.9.2 (*21*). Detection of cell-cell doublets was performed using a two-step process based on the R package DoubletFinder (*89*). DoubletFinder implements a nearest-neighbors approach to doublet detection. First, artificial doublets are simulated from input single-cell data and are co-clustered with true libraries. True doublet libraries are identified by their relative fraction of artificial doublet nearest-neighbors in gene expression space. In our workflow this process is run twice, once with high stringency to identify and remove clear doublets, and again with a lower threshold to identify remaining, subtler doublet libraries. We used the following parameters for round 1: PN = 0.4, PK = 10 ^ seq(−4, −1, length.out=50), NUM_PCS = 5. For the second round: PN = 0.45, NUM_PCS = 10, PK = 10 ^ seq(−4, −1, length.out=50). Because doublet libraries are initially categorized as true libraries in the nearest-neighbor search, we find this two-step process improves the sensitivity and accuracy of doublet detection.

Clustering was performed using independent component analysis (ICA; R package fastICA) dimensionality reduction followed by Louvain. Cells assigned to one of the major glial types (oligodendrocyte lineage, astrocytes, vascular/endothelia, microglia/macrophages) by scPred were collected across all brain regions and clustered together. Neurons from most telencephalic structures (neocortex, hippocampus, striatum, amygdala) confidently assigned to the categories “GABAergic” and “glutamatergic” and so were clustered separately by neurotransmitter usage for each brain structure.

Striatal neuron categories were “SPN” and “GABAergic interneuron”. Neurons in non-telencephalic brain structures were clustered separately by brain structure. All clusterings were performed in two stages: first-round clustering was based on the top 60 independent components (ICs) and a default resolution (res) of 0.1, nearest neighbors (nn) = 25. Following the process outlined in(*6*), each was manually reviewed for skew and kurtosis of gene loadings on factors and cells to identify ICs that loaded on outliers, doublets, or artifactual signals. These were discarded and reclustering was performed on the remaining ICs. Each resulting cluster was then subjected to second-round clustering, during which ICs were again curated.

Second-round clustering explored a range of parameters: nn=10,20,30; res=0.01,0.05,0.1,0.2,0.3,0.4,0.5,1.0. Final parameter values were chosen to optimize concordance, when possible, between the final number of clusters and the number of included ICs, such that each cluster was defined by one primary IC. Metacells for each cluster are generated by summing transcript counts for each cell across all cells in the cluster, normalizing by total number of transcripts, and scaling to counts per 100,000.

### Identification of regionally differentially expressed genes (rDEGs)

We computed rDEGs for neocortical gluatmatergic neurons, GABAergic neurons, astrocytes, and oligodendrocyte lineage types. We used individual cell level cluster assignments from the initial neocortical clustering (which contained all regions together) to create per-region metacells for each cluster. Normalized metacells were log10 transformed and pairwise differences across regions within the same cluster (cell type) were examined. Genes with > 3 fold difference in the same cluster between two regions were considered rDEGs. For this analysis we omitted from comparisons any region-cluster metacell generated from fewer than 50 cells, but retain comparisons between regions that had > 50 cells in that cluster. Genes that were consistently rDEGs in at least 3 individuals for each cluster-region pair are reported in **Table S3**.

### Ancestral State Reconstruction

Ancestral state reconstruction (ASR) is a method to infer hidden ancestral traits from extant observations. For example, given a phylogenetic tree of species and genomic sequences thought to be homologous across those species, the reconstruction takes into account branch lengths to reconstruct the most likely ancestral sequence. We applied a maximum likelihood-based ASR approach (R package: fastAnc) to the scaled, normalized metacells of cell types and the dendrogram of their similarity to produce estimates of expression of each gene at the internal nodes of the tree. This enabled comparisons of leaf nodes to internal nodes as well as internal nodes to each other. For example, compared to the parent node of amygdala, basal forebrain, and hypothalamic SPN-like GABAergic projection neurons, the reconstructed parent node of striatal SPNs had higher expression of known markers of striatal projection neurons such as *DACH1.* We used these reconstructions of internal node gene expression to compare major clades of the tree, and used a Chi-square test to determine whether transcription factors were overrepresented among differentially expressed genes (threshold: 3-fold difference) between pairs of internal nodes.

### Spatial smFISH experiments

All probes are listed in **Table S4**. All smFISH validation experiments were carried out on distinct biological replicates from those used for snRNA-seq or single-cell ATAC-seq experiments.

#### smFISH tissue processing and quantification

Two marmosets (Cj 18-134 and 19-212) were euthanized and perfused with PBS (**Table S1**). The brain was removed, embedded rapidly in OCT, and stored in the −80C freezer. Tissue was then cut to 16µm on a cryostat and stored in the −80C until needed. *In situ* hybridization was performed for genes of interest (see **Table S4**) with HCR or ACD antisense probes, incubated with TrueBlack Lipofuscin Autofluorescence Quencher (Biotium, 23007) for 10 seconds at room temperature to eliminate confounding lipofuscin autofluorescence present in the tissue. Samples were then coverslipped with ProLong Diamond Antifade Mountant (Invitrogen, P36970). Z-stack serial images were taken through the whole depth across striatum, hypothalamus and basal forebrain regions, and several regions of neocortex, on the TissueGnostic TissueFAXS SL slide-scanning, spinning disk confocal microscope (Hamamatsu Orca Flash 4.0 v3) using a 20×/0.8 NA air objective for ACD stains or a 40×/1.2 NA water-immersion objective for HCR stains.

Series images were segmented using StrataQuest, a software package from TissueGnostics, which enables the quantification of signals within segmented images (similar to CellProfiler). Nuclei objects were generated using the DAPI channel, and artifacts were removed based on size and intensity. Exclusion ROIs were manually drawn to avoid areas of white matter, large artifacts, and autofluorescence before computing intensity and other statistical and morphological measurements (20 parameters) for each channel. Specifically, 50 cells were hand-labeled as positive or negative for the markers of interest in order to identify the appropriate threshold for feature selection using the parameters that best discriminated this binary. Parameters included mean intensity, maximum intensity, standard deviation of intensity, and range of intensity, and equivalent diameter. These filters were then applied to the unlabeled data to identify positive cells.

Segmented cells were further analyzed using in-house code (https://github.com/klevando/BICCN-StrataQuest-Script, **Table S7**). Spatial locations of the cells were visualized by plotting the x-y coordinates associated with each nuclei. These were then binned into 2-D histograms across the x and y axis (corresponding to the rostrocaudal plane and to the dorsoventral plane respectively). A size of 100 bins was chosen for the first (medial-most) slice of a series across the x axis and the calculated bin size was then used across the y axis of the first slice and across other slices in the series. Positive events for a gene in a given bin were either normalized to the number of detected nuclei in that bin (DAPI) and plotted in 2-D as a relative heatmap, or simply plotted as a density heatmap without being normalized to DAPI. Whole slice normalizations across the mediolateral axis (all positive events for a gene in a given slice relative to DAPI) were plotted as bar graphs. DAPI and marker of interest counts are available in **Data** S1-S2.

#### Neocortical Areal Proportions

In addition to the mediolateral subdivisions of marmoset neocortex, we further parcellated the neocortex into subregions referencing the Brain/MINDS 3D Digital Marmoset Atlas (https://doi.org/10.24475/bma.4520). Slice selection was based upon visual recognition of prominent landmarks (white matter, striatal boundary as well as DAPI nuclei staining) within our tissue and matched with the nearest atlas slice. ROIs for each parcellated region were created using the StrataQuest software in the anterior-posterior and dorsal-ventral axes (**Fig. S8C**). The corresponding feature selection that was previously set within the full neocortical sections was carried across gene and slice. The parameters used were mean intensity, maximum intensity, standard deviation of intensity, and range of intensity, and equivalent diameter. These filters were then applied to the unlabeled data to identify positive cells within each individual ROI. The parcellated neocortical areas were then analyzed using in-house code (https://github.com/klevando/BICCN-StrataQuest-Script, **Table S7**). As above, positive events for a gene in a given ROI were normalized to DAPI and plotted in a (stacked) bar graph. Relative percentages between genes were calculated and plotted stacked.

### Morphology and smFISH experiments

AAV9-hDlx5/6-GFP-fGFP virus was generated as in (*19*). Virus was systemically IV injected (400ul-700ul at 1.7-2.4^10^ titer) in 5 marmosets. The virus was allowed to incubate for approximately 2 months. After systemic IV injection with AAV9-hDlx5/6-GFP-fGFP, marmosets were euthanized and perfused with saline followed by 4% paraformaldehyde (PFA). Brains were removed and 120µm sections were cut on a vibratome into PBS-Azide and stored at 4C or moved into 70% ethanol for storage at −20C. 70% ethanol storage prevents RNA degradation at this temperature without significant tissue shrinkage for short storage times. Due to lab shutdowns during the pandemic, sections from two marmosets were stored in 70% ethanol for approximately 4 months. These samples exhibited significant shrinkage, measured by DAPI-stained nuclei diameters (**Table S1**), therefore we only compared morphology parameters within-donor. Sections were taken as needed and *in situ* hybridization was performed with HCR antisense probes, following the generic sample in solution HCR protocol with a 2-fold increase in concentration of probe to hybridization buffer (Molecular Instruments, https://files.molecularinstruments.com/MI-Protocol-RNAFISH-GenericSolution-Rev8.pdf), for markers of interest (**Table S4**) that corresponded with RNA-seq defined clusters. The sections were then stained with anti-GFP antibody (**Table S4**) and a secondary antibody (**Table S4**) to amplify the endogenous GFP signal (https://www.protocols.io/view/marmoset-nhp-free-floating-anti-gfp-antibody-stain-3byl47nb2lo5/v1, **Table S4**). Sections were incubated in TrueBlack (Biotium, 23007) for 3-5 minutes in order to mask confounding lipofuscin autofluorescence throughout the section. Sections were then mounted onto a slide and coverslipped with ProLong Diamond Antifade Mountant (Invitrogen, P36970) for imaging.

#### Imaging for morphology

Sections prepared for morphology were imaged on a Nikon Ti Eclipse inverted microscope with an Andor CSU-W1 confocal spinning disc unit and Andor DU-888 EMCCD using a 40×/1.15 NA water-immersion objective, and later on a TissueGnostic TissueFAXS SL slide-scanning, spinning disk confocal microscope (with Hamamatsu Orca Flash 4.0 v3) using a 40×/1.2 NA water-immersion objective. With the TissueFAXS, overview images were taken in order to select GFP+ cells for imaging at 40× and to highlight the exact location of the cell. GFP+ cells were imaged for stained markers of interest. Selected sections were imaged on an upright confocal laser scanning microscope (Olympus Fluoview FV3000) using a 40x/0.95 NA air objective or a 60x/1.50 NA oil-immersion objective and cooled GaAsP PMTs.

#### Morphological reconstruction and feature quantification - Imaris

Without pre-processing the confocal images, three-dimensional (3D) reconstruction and surface rendering of striatal and neocortical interneurons were performed using Imarisx64 9.9 software (Oxford Instruments) based on GFP+ signal. Surface-rendered images were used to determine the soma diameter, total volume, and total surface area for each z-stack image. 3D-skeleton diagrams (**Figs. 5A-C** and **6H**), corresponding to each surface-rendered image (data not shown), were generated using the Filament Tracing wizard in Imaris and then pseudo-colored in Adobe Illustrator. The total number of primary dendritic branches, dendritic branch points, area, volume, and length of the 3D-skeleton diagrams were calculated using the AutoPath (no loops) algorithm in the filament tracing wizard in Imaris. The total number of primary dendritic branches for each cell is defined by the number of dendrite branches in the filament trace 1 distance value away from the soma. The distance value is calculated automatically by the AutoPath (no loops) algorithm based on the diameter of the soma and the diameter of the thinnest cellular process. All data were exported to CSV, and data collected from exemplar cells (**Fig. 5A-B** and **6G)** were reported in **Table S6**.

A separate dataset containing GFP+ interneurons from one animal (Cj 17-154) was first blinded using a custom Python script, then reconstructed with Imaris (**Figs. 5C-D**). All results are presented as mean ± SEM. Comparisons between soma diameter, surface area, surface volume, and the number of primary branches were carried out using unpaired *t-*tests. Unpaired Mann-Whitney *U* tests were used to statistically compare the total length, area, volume, and total number of dendritic branch points measurements. Statistically significant analyses were denoted as follows: \**p* < 0.05; \*\**p* < 0.01.

### Morphological reconstruction and feature quantification - NeuTube

Automatic or semi-automatic tracing algorithms are challenged by some data, perhaps due to the low SNR of a given image. To overcome this, we manually reconstructed the sparse neurons via Neutube (*90*) tracing software. With the software, we 1) Create 3D volume rendering of the GFP-AAV marmoset neuron, 2) use the signal transfer function (e.g., histogram equalization) for overall intensity and opacity values to optimize the signal-to-noise ratio by manually examining the clearest visualization of the dendrites, 3) Build the neuron skeleton over the 3D volume by tracing the processes, 4) Scan through the 3D volume to make sure no parts of the neuron are missed, 5) Double check the raw 2D images to see if any of the branches were not presented well in 3D due to volume rendering artifacts, 6) Label axon, dendrites and soma parts of the skeleton model. Reconstructed neurons were saved as SWC format.

### Single nucleus ATAC-seq, library preparation, sequencing

Single nucleus ATAC-seq (snATAC-seq; 10x Genomics Single Cell ATAC v1) was conducted on fresh marmoset tissue (1 female, 1.5 y.o.) dissected from anterior striatum. Nuclei suspensions were generated following the 10x recommended protocol (CG000212 Rev B). Library preparation followed 10x Genomics Single Cell ATAC v1 Guide (CG000168 Rev D). Library was sequenced on an Illumina NovaSeq (RRID:SCR_016387) using 100 bp paired-end reads to a median per-cell fragment depth of 25,472. Alignment and fragment counting was conducted using Cell Ranger (RRID:SCR_01734), aligned to marmoset genome cj1700 (https://www.ncbi.nlm.nih.gov/assembly/GCF_009663435.1/; GCA_009663445.2). snATAC-seq data were integrated with marmoset striatal snRNA-seq data from two independent animasl (bi005, bi006) using Signac (RRID:SCR_021158) with the following parameters: integration method: cca; weight.reduction = lsi. Differentially accessible peaks for each cluster were calculated with a 3-fold change cutoff and the minimum fraction of expressed cells (in target cluster) = 0.2. Enhancer locations were as follows (CJ1700 coordinates): Eh14 = chr1-122696345-122696560 (215 bp); Eh15 = chr17-32919427-32919564 (137 bp); Eh16 = chr20-38770509-38770619 (110 bp); Eh17 = chr9-60258694-60258788 (94 bp).

### TAC3-AAV viral design and production

AAVs were produced in accordance with the protocol (*91*). 5.7 μg/150 mm dish of construct DNA was transfected with 22.8 μg/150 mm dish of pAAV9 capsid plasmid and 11.45 μg/150 mm dish of pAdDeltaF6 helper plasmid using polyethylenimine 25K MW (Polysciences, 23966-1). Collection of cells and media for AAV harvesting began 72 hours following transfection before iodixanol gradient ultracentrifugation purification using a Type 70 Ti Fixed-Angle Titanium Rotor (Beckman-Coulter, 337922). Titer was calculated using digital droplet PCR (ddPCR) according to the protocol “<u>ddPCR</u> <u>Titration of AAV Vectors</u>” (https://www.addgene.org/protocols/aav-ddpcr-titration/) using the QX200 AutoDG Droplet Digital PCR System (BioRad, 1864100). gBlock fragments containing the tandem *TAC3* enhancers (E14E15E16E17) were synthesized by Integrated DNA Technologies (IDT). The enhancers were cloned into AAV plasmid vector backbones containing a minimal CMV promoter, the reporter and a barcode unique to each enhancer sequence using the NEBuilder® HiFi DNA Assembly Cloning Kit (NEB-E5520S), following standard protocol. 2 uL of Gibson assembly product was used to transform 50 uL of home-made Stbl3 cells following a standard transformation protocol. Mini-preparation of plasmids was carried out using the NucleoSpin mini kit (Macherey-Nagel). Positive clones were identified by restriction enzyme digestion and sequencing. The positive clones were grown in 300 mL LB cultures and the plasmids were extracted using the NucleoBond Xtra Midi EF kit (Macherey-Nagel). The plasmids were sent out for sequencing again before being packaged into AAVs.

### TAC3-AAV local injection procedure

#### Structural MRI Scanning

In preparation for MR imaging under anesthesia, an animal (2 marmosets, Cj 19-207 and 17-B111, **Table S1**) checked for robust health was fasted overnight with *ad libitum* access to water. Prior to scanning, the animal was sedated with Alphaxalone (5-10 mg/kg) or a combination of Alphaxalone (4-8 mg/kg) and Ketamine (5-10 mg/Kg) given intramuscularly. It was then transferred to a custom designed MRI cradle equipped with a 3D printed nose cone for delivering anesthetic gases. The animal’s head was securely held to minimize motion artifacts and stereotaxically align the rostral-caudal axis with the MR scanner bore using contrast filled MR compatible ear bars, an adjustable palette bar and eye bars. During scanning, anesthesia was maintained at 1-2% isoflurane in a mixture of oxygen. A temperature controlled water circulating heat pad was used for thermal support. Heart rate and blood oxygen saturation levels were continuously monitored and recorded every 10 min with an MR compatible monitoring system (Nonin, MN). MR scans consisted of high-resolution 3D T1 mapping using a Magnetization-Prepared Rapid Gradient-Echo (MPRAGE) sequence (Liu et al., 2011) with repetition time (TR) = 6000 ms, TE (echo time) = 4.6 ms; flip angle (FA) =12 degrees; FOV = 80, matrix size = 256×256. A high-resolution T2-weighted anatomical scan was also obtained using a RARE pulse sequence with effective TE = 35.5 ms, TR = 2500 ms, RARE factor = 8, spatial resolution = 156 µm × 156 µm × 0.5 mm, and matrix size = 256 × 256, RARE factor = 4, 80 mm FOV.

#### Virus injection surgery

Following sedation with Alphaxalone (5-10 mg/kg), the animal was intubated and maintained at 1.5-2% isoflurane in a mixture of oxygen throughout the rest of the procedure. An intravenous (IV) catheter was placed in the saphenous vein for infusion of fluids and drugs during the surgery and the recovery period. Heart rate, SpO2, ECG, end-tidal CO_2_ and rectal temperature were continuously monitored and logged every 5 minutes throughout. Once the animal acquired a stable plane of anesthesia, it was placed in a stereotaxic apparatus (Narishige, Japan). A thin layer of sterile eye lubricant was applied to protect against corneal drying. Animal was also provided with a single bolus dose of intravenous dexamethasone (0.4 mg/kg) to guard against brain swelling. The scalp and fascia were removed in layers via blunt dissection to expose the injection sites. Using MRI atlas guidance, bilateral craniotomies were performed to expose the medial anterior caudate and ventral posterior putamen region. The tandem enhancer virus AAV9-tandemE-TAC3-EGFP was injected in left Caudate (high titer, 10^13 low volume, 0.5 ul) and left Putamen (low titer, 10^11, high volume 1.5 ul). Each injection was delivered with a controlled syringe pump (100 nL/minute). After each injection the needle was kept in place for 10 minutes then slowly retracted over 1-2 minutes to minimize injection backflow. Following withdrawal, the cranial openings were covered with bonewax, fascia and scalp were sutured in layer wise manner with 3-0 vicryl sutures. The suture site was cleaned with warm sterile saline and covered with hypafix. For recovery the animal was transferred in an incubator and monitored closely till it was able to move effortlessly and accepted treats before returning to its home cage. Post-op medications were provided under veterinary advice for pain and inflammation control until full recovery. The animals were euthanized 6-10 weeks after viral injection and were perfused transcardially with 4% PFA. The brain was extracted for histology, smFISH, and confocal microscopy.

### TAC3-AAV-BI103 systemic injection procedure

One 2-year old female marmoset (**Table S1**) was sedated with Alphaxalone (5-10 mg/kg) given intramuscularly. A catheter was placed into the tail vein and 200 μl of purified viral particles were injected at a dose of 7.85×10^13^ vg/kg, followed by injection of 1.5 ml saline to flush the line. The animal was euthanized 4 weeks after viral injection and was perfused transcardially with 4% PFA. The brain was extracted for histology, smFISH, and confocal microscopy.

### TAC3-AAV histology

For AAV9-tandemE-TAC3-EGFP experiments, coronal sections around the injection sites were cut at 100 μm on a vibratome (Leica, VT1000S) and then stored in PBS-NaN3 at 4 °C until use. For AAV-BI103-tandemE-TAC3-EGFP experiments, coronal and sagittal sections were processed. Sections were stained for *TAC3* (Molecular Instruments, PRC843, **Table S4**) using the HCR v.3.0 protocol (Molecular Instruments) described above and a rabbit anti-GFP antibody (Invitrogen, A11122, **Table S4**) with subsequent secondary goat anti-rabbit conjugated with AF488 (ThermoFIsher, A-11008). Stained sections were then incubated 1-3 minutes in TrueBlack to eliminate confounding lipofuscin autofluorescence. The striatum was imaged for each section using the TissueFAXS SL with a 40×/1.2 NA water-immersion lens through the whole thickness of the tissue.

## Acknowledgements

We thank Tim Blosser for his early involvement in developing the spatial transcriptomics workflows. We thank Atsushi Takahashi for assistance with MRI scanning for animals receiving local viral injections. We thank the MIT veterinarian staff for animal husbandry and for their assistance with surgical procedures. We thank Monika Burns and Yuanyuan Hou for assistance with AAV IV injections and animal perfusions.

## Funding

National Institutes of Health grant U01MH114819 (GF, SAM, EB)

The National Institute of Neurological Disorders and Stroke grant UG3NS111689 (BED) NSF GRFP # 1745302 (MES)

MathWorks Science Fellowship (MES) Collamore-Rogers Fellowship at MIT (MES) NSF GRFP # 1122374 (TWS)

Broad Institute’s Stanley Center for Psychiatric Research (SAM, GF, BED) Dean’s Innovation Award (Harvard Medical School) (SAM)

Hock E. Tan and K. Lisa Yang Center for Autism Research at MIT (GF) Poitras Center for Psychiatric Disorders Research at MIT (GF) McGovern Institute for Brain Research at MIT (GF)

## Author Contributions

RNA/ATAC Data Generation: FMK, MG, AL Spatial Data Generation: KML, HZ

Data Analysis: FMK, KML, HZ, RCHdR, MES, MG, AL, KXL, VFB-G, TWS, SAM

Data Interpretation: FMK, KML, HZ, RCHdR, MES, MG, KXL, VFB-G, SB, EB, SAM, GF

Tissue Samples/Tissue Processing: FMK, KML, HZ, MG, AL, QZ, GC, SB AAV design, generation & experiments: MW, CC, QZ, JS, SJV, JD, KC, BED Morphology Data generation/analysis: KML, HZ, VFB-G, TWS, AM, EB Software/Data management: FMK, RCHdR, MG, AW, JN, SK

Writing: FMK, KML, HZ, MES, VFB-G, SAM, GF

Competing interests: Authors declare that they have no competing interests.

## Data and materials availability

Raw sequence data were produced as part of the BRAIN Initiative Cell Census Network are available for download from the Neuroscience Multi-omics Archive (https://assets.nemoarchive.org/dat-1je0mn3) and the Brain Cell Data Center (https://biccn.org/data). Morphological reconstructions and single molecule FISH of interneuron types are available for download through the Brain Image Library (https://submit.brainimagelibrary.org/search?grant_number=1-U01-MH114819-01). The AAV-BI103 rep-cap plasmid will be made available through Addgene upon publication of the characterization of this capsid and through direct requests to Ben Deverman.

## Supplementary Materials

**Figure S1.**
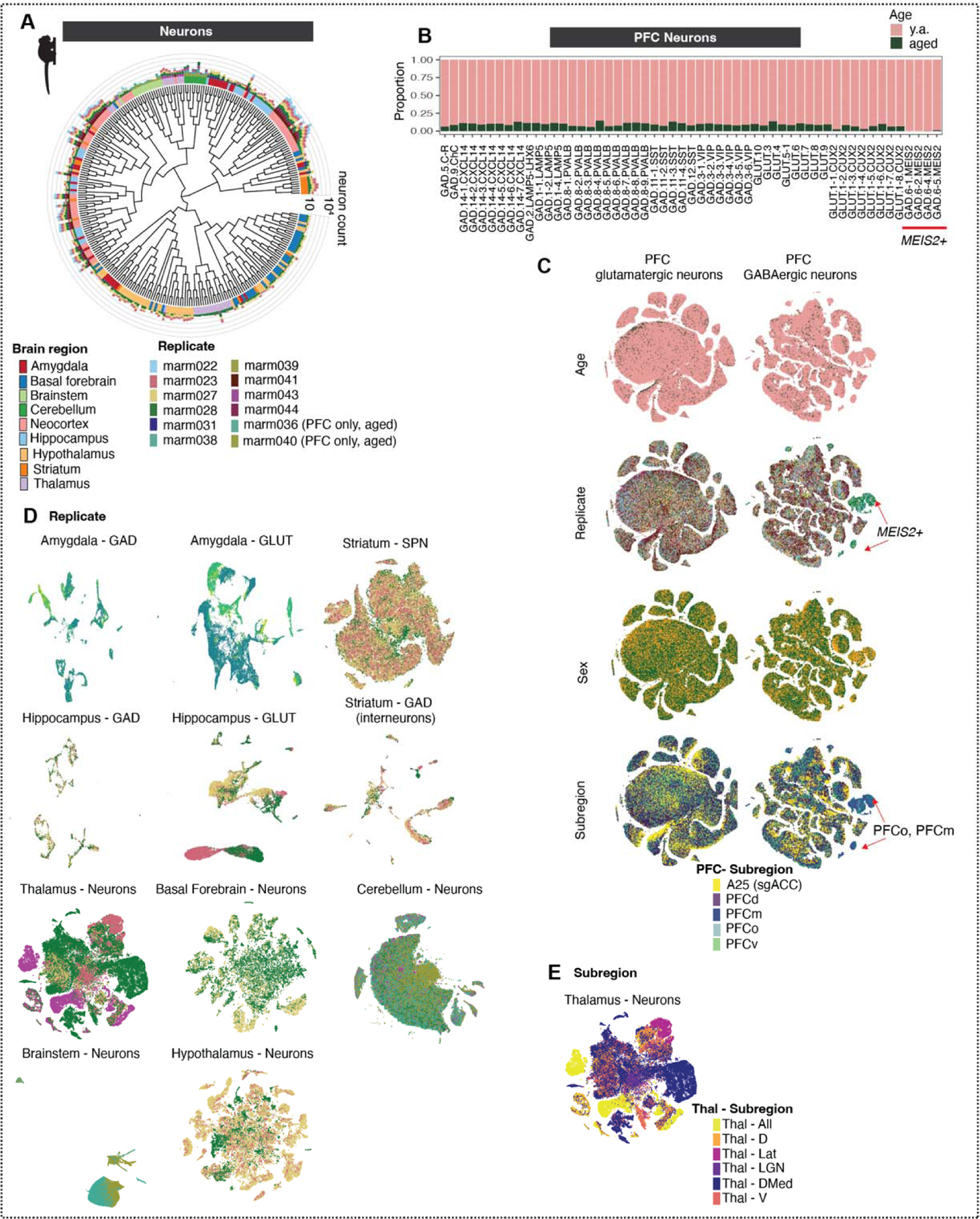
Neuron counts by donor across brain regions. **(A)** Neuronal dendrogram as in Fig. 1C, with outer barplots depicting number of nuclei per cell type and replicate. Ring colors are brain regions, colors in barplots correspond to replicates. **(B)** Proportional per-cluster representation of PFC neurons between young adult donors and aged (n=2) donors. While our snRNA-seq collection focused on post-sexual maturity young adults, we acquired an additional dataset of PFC sampled from 2 aged animals (1 M, 11y0m; 1F, 14y4m, 37,260 cells total; **Table S1**). Individual replicates contributed similar proportions of neurons to each prefrontal neuron subtype, and clusters generally had proportional representation across young adults and aged animals, as well as across males and females (**Fig. S1B-C**), suggesting that these variables do not dramatically impact neuronal ensembles and identities in prefrontal cortex. **(C)** t-SNE embeddings of PFC neurons (*top row*, GABAergic; *bottom row*, glutamatergic) with colors representing different metadata: age (young vs aged), replicate, PFC subregion. There was notable enrichment of *MEIS2+* GABAergic neurons in medial prefrontal and orbital prefrontal dissections **(Fig. S1C)**. Based on their gene expression profiles, these cells likely correspond to the recently described population of LGE-derived *MEIS2*+ neurons that populate the olfactory bulb in mice, and which are instead directed to medial prefrontal cortex in macaques and humans (*33*). **(D)** t-SNEs of neurons in each brain structure, with cells colored by replicate (colors as in (A)). Telencephalic neurons are plotted separately by class: GABAergic and glutamatergic classes (neocortex, hippocampus, amygdala), or GABAergic interneurons and spiny projection neurons (striatum). Compared with neocortex, greater cross-donor variability was observed in some subcortical structures such as hypothalamus and thalamus, though this was likely driven more by dissection variability than by donor variability, as the donor-specific clusters tended to be from subregions that were only sampled in one individual (see E). **(E)** t-SNEs of thalamic neurons with cells colored by thalamic subdivision.

**Figure S2.**
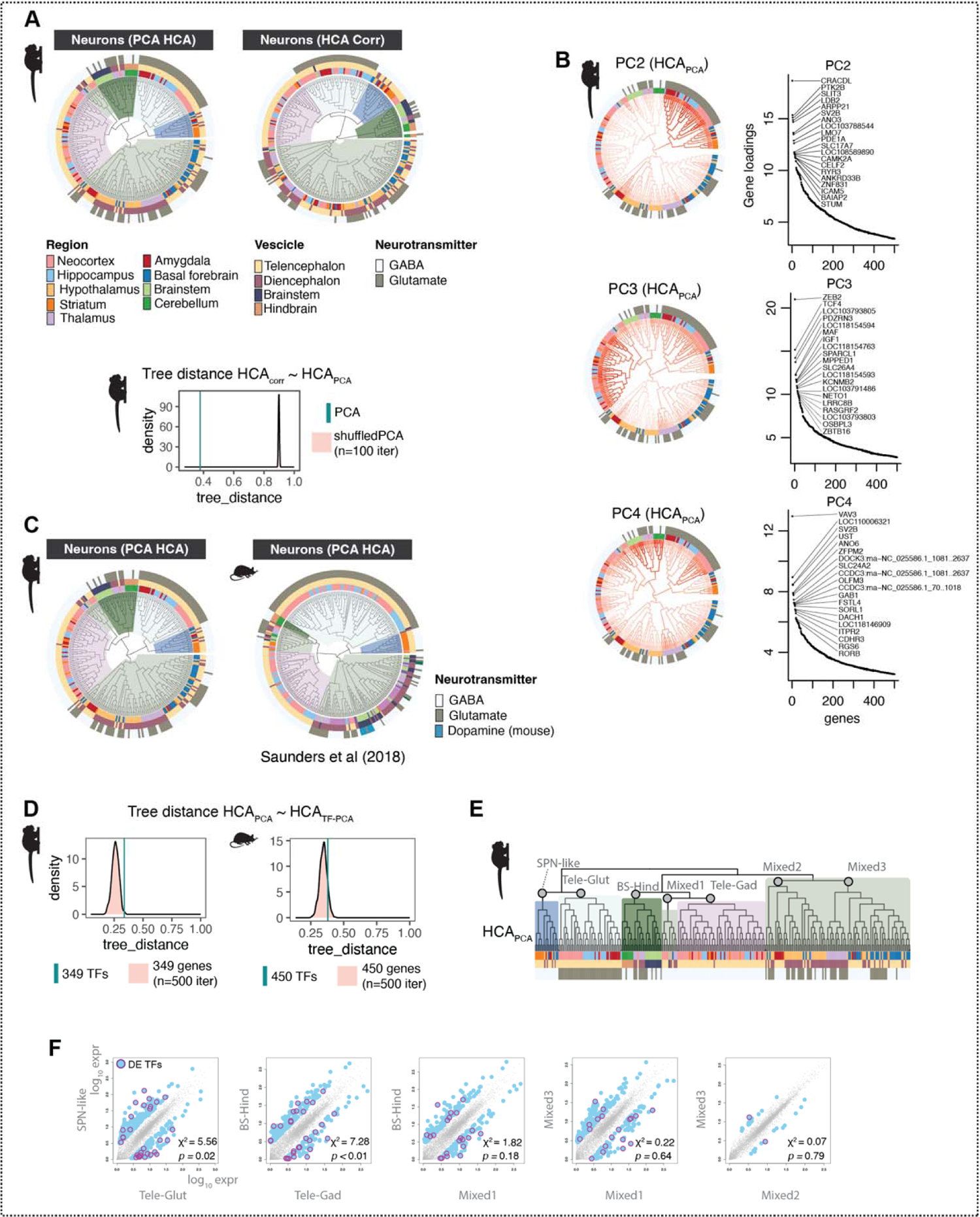
Conservation of neuronal hierarchy across species and clustering methods. (A) Comparison of marmoset and mouse neurons (mouse atlas data from (*6*)) using three different distance calculations for hierarchical clustering: HCA_corr_ distance = gene expression correlations of top 5904 genes in marmoset or 3528 in mouse (mouse genes include all genes with 1:1 orthologs to the 5904 marmoset genes and that are expressed in at least 10 transcripts per 100,000 in at least one mouse neuron type); HCA_PCA_ distance = top 100 PCA scores across the same genes; HCA_TF-PCA_ distance = top 100 PCA scores using expressed transcription factors only (marmoset = 349 TFs, mouse = 450 TFs). **(B)** Distance between hierarchical clustering (HC) dendrogram trees computed using different methods. Cyan line = tree distance (R package TreeDist) between hierarchical clustering using distance = HCA_corr_ and using distance = HCA_PCA_. Pink distribution is tree distance scores between HCA_corr_ and shuffled PCA scores (n = 100 shuffling iterations). Lower values of tree_distance (x-axis) mean higher agreement between dendrogram tree structures. **(C)** PCA loadings and top genes for PC2-PC4. PC scores are plotted on the HCA_PCA_ dendrogram. Ranked gene loading plots show top 20 genes per PC. **(D)** Tree distances computed as in (B) between HCA_PCA_ and HCA_TF-PCA_. The tree distances between these two trees is low, but not different from distributions of random, same-sized sets of genes. **(E)** Marmoset dendrogram in Fig. 1C (HCA_PCA_) indicating major clades compared in (F). **(F)** Ancestral reconstruction (AR; R package phytools) of gene expression profiles of major clades of marmoset neuron types from dendrogram in (E). Maximum likelihood estimates of gene expression (fastAnc) were computed for 7 major internal nodes (gray circles) of the HCA_PCA_ dendrogram. Scatterplots show pairwise comparisons between AR of internal nodes of major clades. Blue dots = genes with >3 foldchange difference between the two ARs. Magenta circles = differentially expressed transcription factors (DE-TFs). Chi-square and p-values describe whether TFs are significantly differentially enriched between the AR pairs.

**Figure S3.**
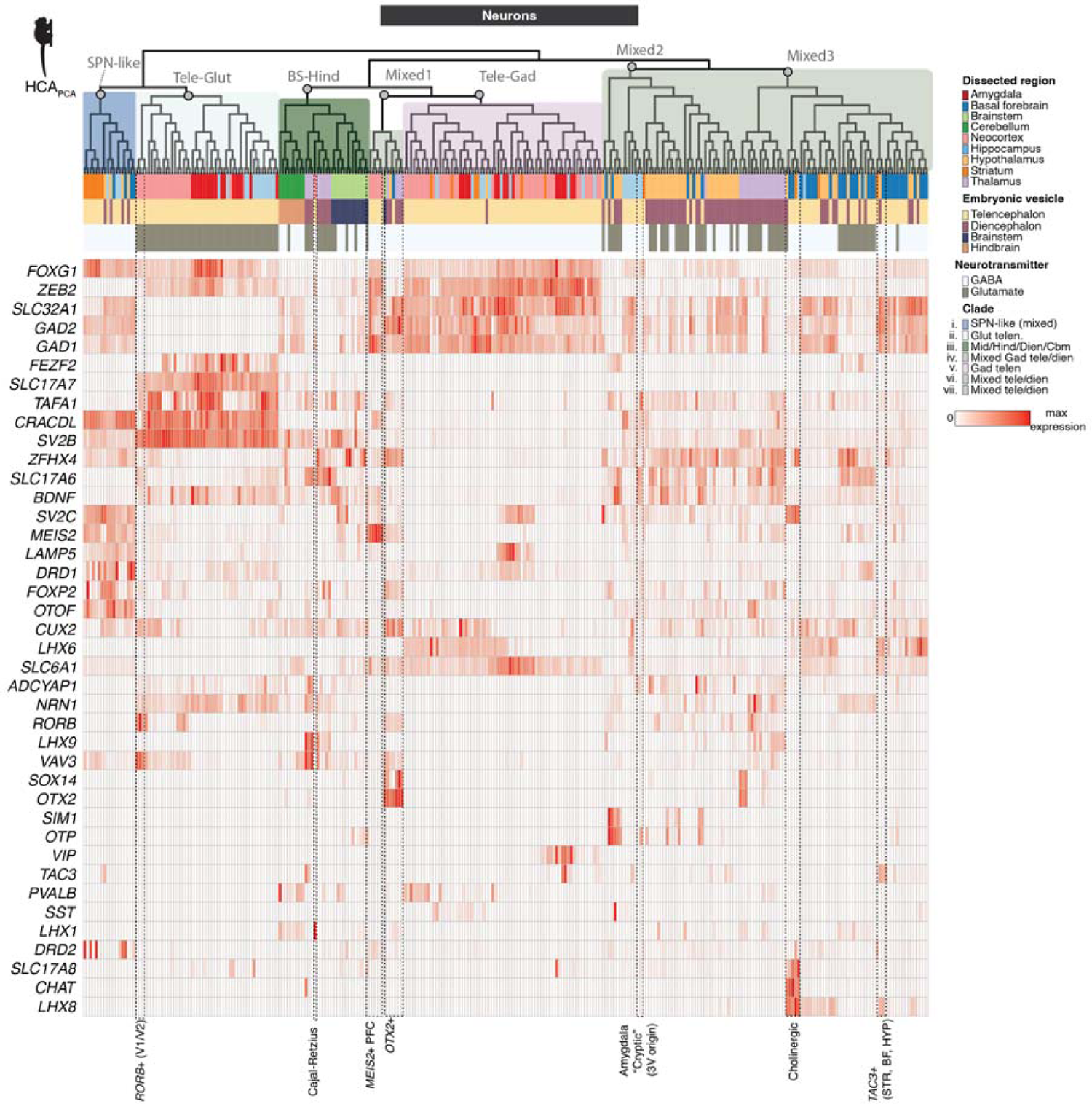
Gene expression across neural populations. Expression of broad class marker genes and other genes of interest across all neurons sampled by snRNA-seq. Heatmap colors are scaled to max normalized expression for each row (gene). Dendrogram ordering and metadata colors as in Fig. 1C. Cell types discussed in the main text are labeled at bottom.

**Figure S4.**
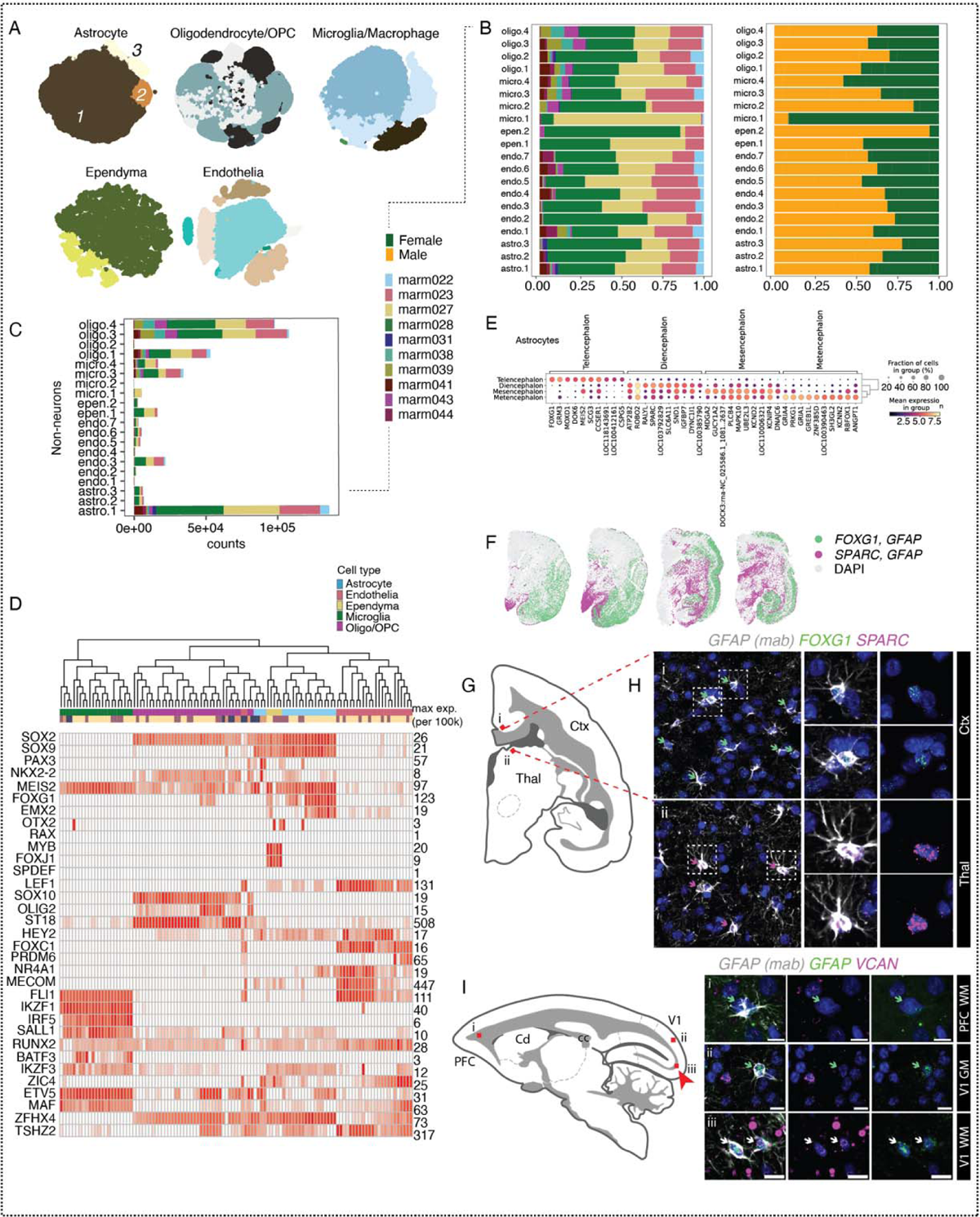
Glia diversity across regions. (A) t-SNE embeddings of major non-neuronal types colored by cluster. **(B)** Barplots of glial proportions colored by donor and by sex. **(C)** Non-neuronal nuclei counts. Colors indicate donor, same as (B). **(D)** Expression of marker genes in non-neurons. Genes as in (*8*). Heatmap colors are scaled to max normalized expression for each row (gene). Dendrogram and metadata colors as in Fig. 1G. **(E)** Differentially expressed genes in astrocytes across cephalic compartments. **(F)** Tissue validation for astrocyte differentially expressed genes (*FOXG1, SPARC*) in coronal sections of marmoset brain. Green dots indicate locations of cells that stain positive for *GFAP* (IHC, mAb **Table S4**) and *FOXG1* (smFISH). Magenta dots indicate cell positions for *GFAP* (IHC) and *SPARC* (smFISH). **(G)** Cartoon of coronal section imaged; Red boxes (*i-ii*) correspond to tissue validation in (H). **(H)** (*Left)* Fields of view from neocortex and thalamus stained for *GFAP* antibody (gray), *FOXG1* (green), and *SPARC* (magenta). Green arrows highlight *GFAP* cells colocalized with *FOXG1*, magenta arrows highlight *GFAP* cells colocalized with *SPARC.* (*Right*) Magnified examples of double positive cells in neocortex and thalamus. Ctx = cortex, Thal = thalamus. **(I)** (*Left*) Cartoon of sagittal section imaged; red boxes (*i-iii*) correspond to (*Right*) tissue validation of increased abundance of *VCAN*+ astrocytes in adult marmoset V1-adjacent white matter (*iii*) compared with PFC-adjacent white matter (*i*) and V1 gray matter (*ii*). GFAP antibody (gray) combined with smFISH probes against *VCAN* (magenta) and *GFAP* (green). Green arrows correspond to *GFAP*+ (antibody), *GFAP*+ (smFISH) cells. White arrows correspond to *GFAP*+ (antibody), *GFAP*+ (smFISH), and *VCAN*+ cells. V1 = visual cortex V1, PFC = prefrontal cortex, GM = gray matter, WM = white matter. Red arrow highlights locale of *VCAN+ GFAP+* images. Scale bar = 10 µm.

**Figure S5.**
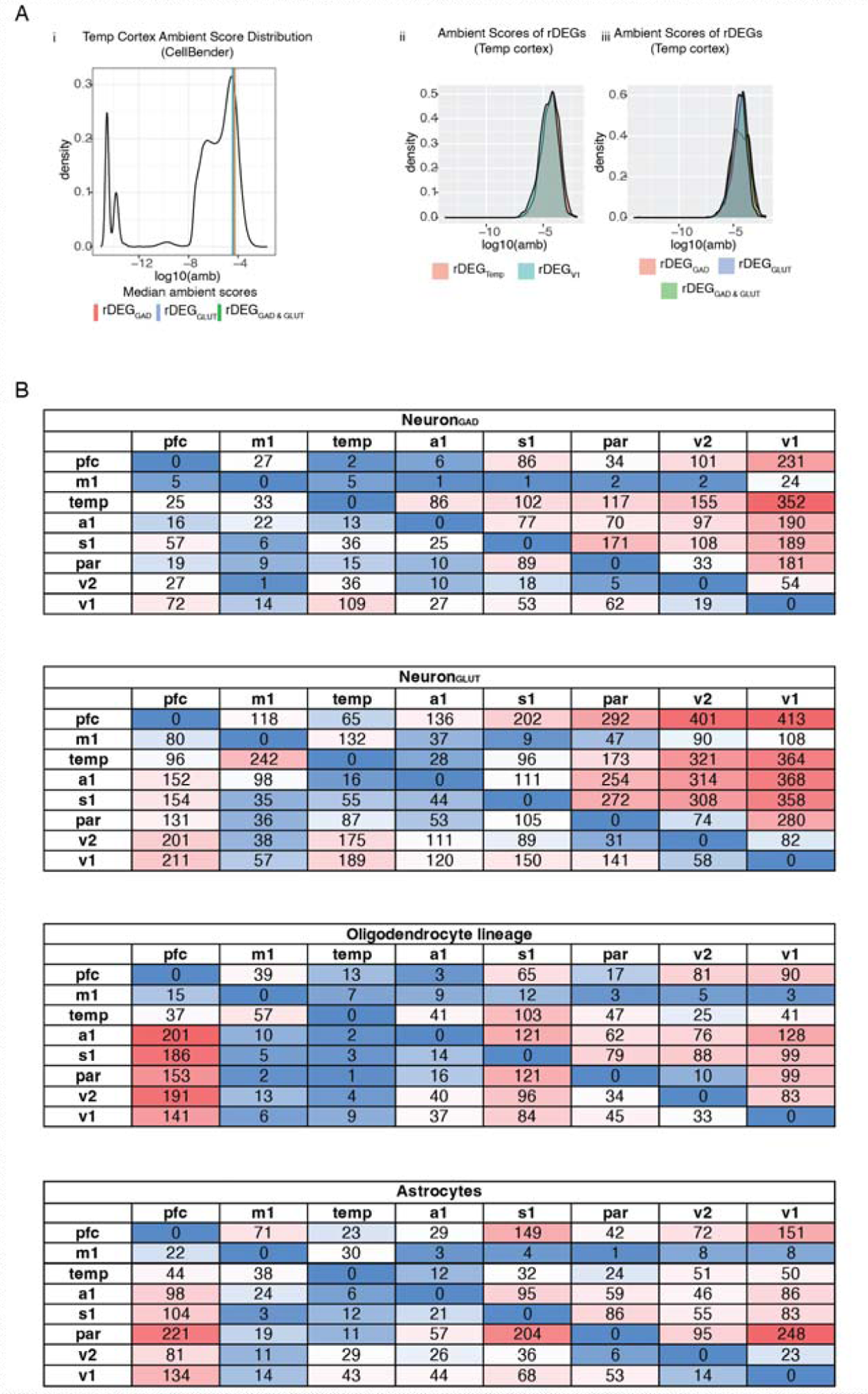
Cortical rDEGs are defined across cell types and do not reflect ambient RNA contamination. **(A) (***i)* rDEG ambient score distributions (from CellBender) in temporal cortex samples, which had amongst the highest numbers of rDEGs compared to other neocortical regions. Despite glutamatergic neurons being more numerous and having more expressed genes/transcripts per cell, the median ambient contamination scores for glutamatergic rDEGs were not higher than median contamination scores for GABAergic rDEGs. rDEGs shared between glutamatergic and GABAergic neurons had indistinguishable scores compared with rDEGs private to one neuronal class. *(ii)* Ambient scores in temporal cortex of temporal cortex rDEGs are indistinguishable from V1 rDEGs. *(iii)* Distributions of temporal cortex rDEG ambient scores by neuron class, again showing no difference between rDEGs that are shared or private to a neuronal class. **(B)** Numbers of regionally differentially expressed genes (rDEGs) between pairs of cortical regions for neurons, astrocytes, and oligodendrocyte lineage types.

**Figure S6.**
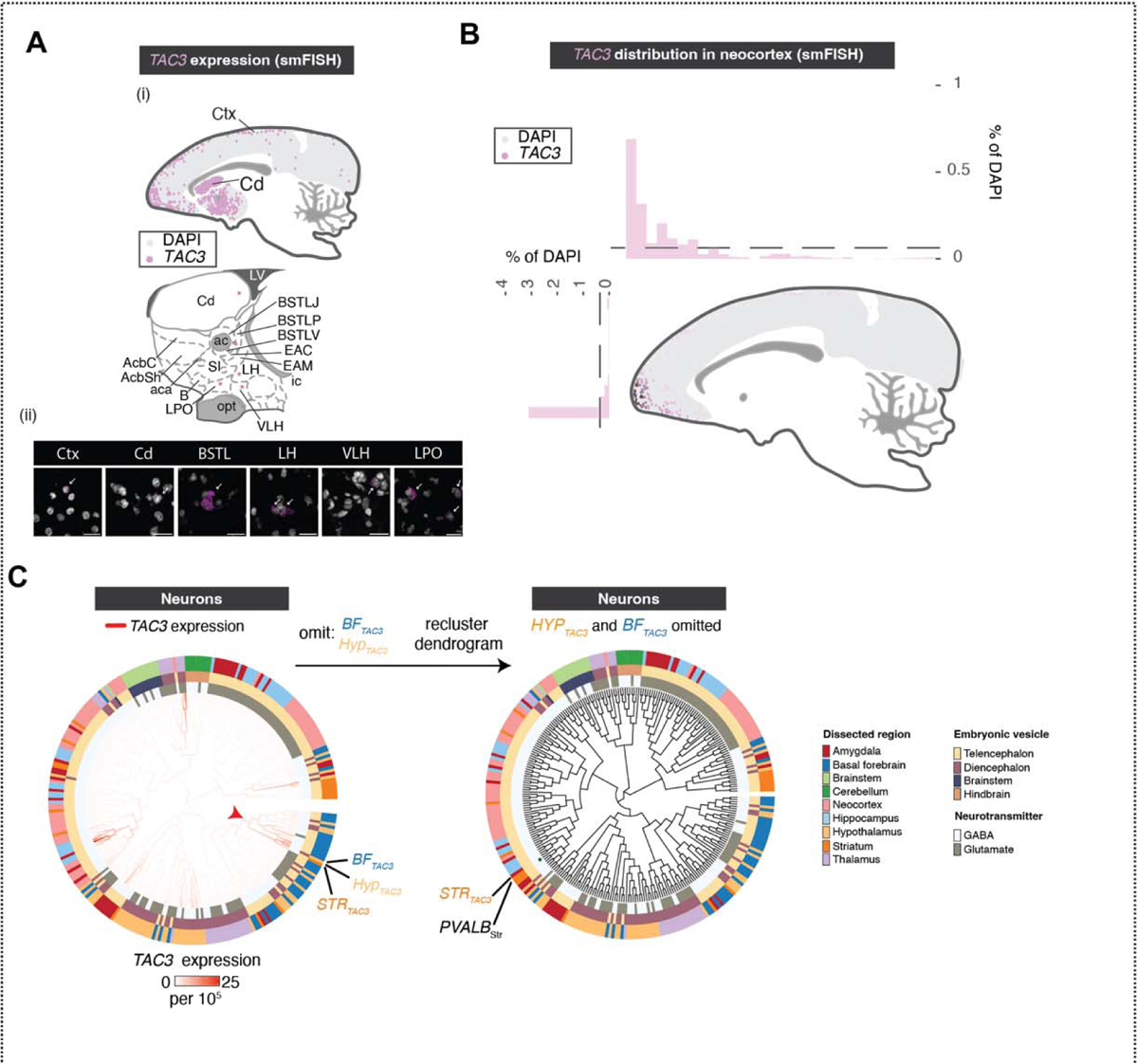
Locations of *TAC3*+ cells in marmoset forebrain. **(A)** smFISH reveals anatomical locations and expression levels *TAC3+* types in different brain regions. *(i)* Schematic of *TAC3*+ cells imaged across cortex, dorsal striatum. Ctx = neocortex, Cd = Caudate. *(ii)* Cartoon close-up of nuclei in striatum, basal forebrain and hypothalamus. Magenta stars = locations of *TAC3*+ cells in lower image panel. Cd = Caudate, AcbC = nucleus accumbens core, AcbSh = nucleus accumbens shell, SI = Substantia innominata, B = basal nucleus of Meynert, EAM = extended amygdala, medial, EAC = extended amygdala, central, ac = anterior commissure, BSTLP = bed nuc st, lateral posterior, BSTLJ = bed nuc st, juxtacap, BSTLV = bed nuc st, lateral ventral, LH = lateral hypothalamus, VLH = ventrolateral hypothalamus, LPO = lateral preoptic area. **(B)** Density and location of *TAC3*+ cells as proportion of all DAPI+ cells. Barplots show percentages in bins (approximately 1,290 µm per bin) taken across the anterior-posterior (top) and dorsal-ventral (left side) axes. **(C)** Effect on placement of the *TAC3+* striatal type on the neuronal dendrogram when omitting the two *TAC3*+ types in hypothalamus and basal forebrain. When these types are omitted and hierarchical clustering is repeated (using HCA_PCA_), the *TAC3*+ striatal type is most similar to *PVALB*+ striatal interneurons, consistent with previous reports that only compared telencephalic interneurons (*3*).

**Figure S7.**
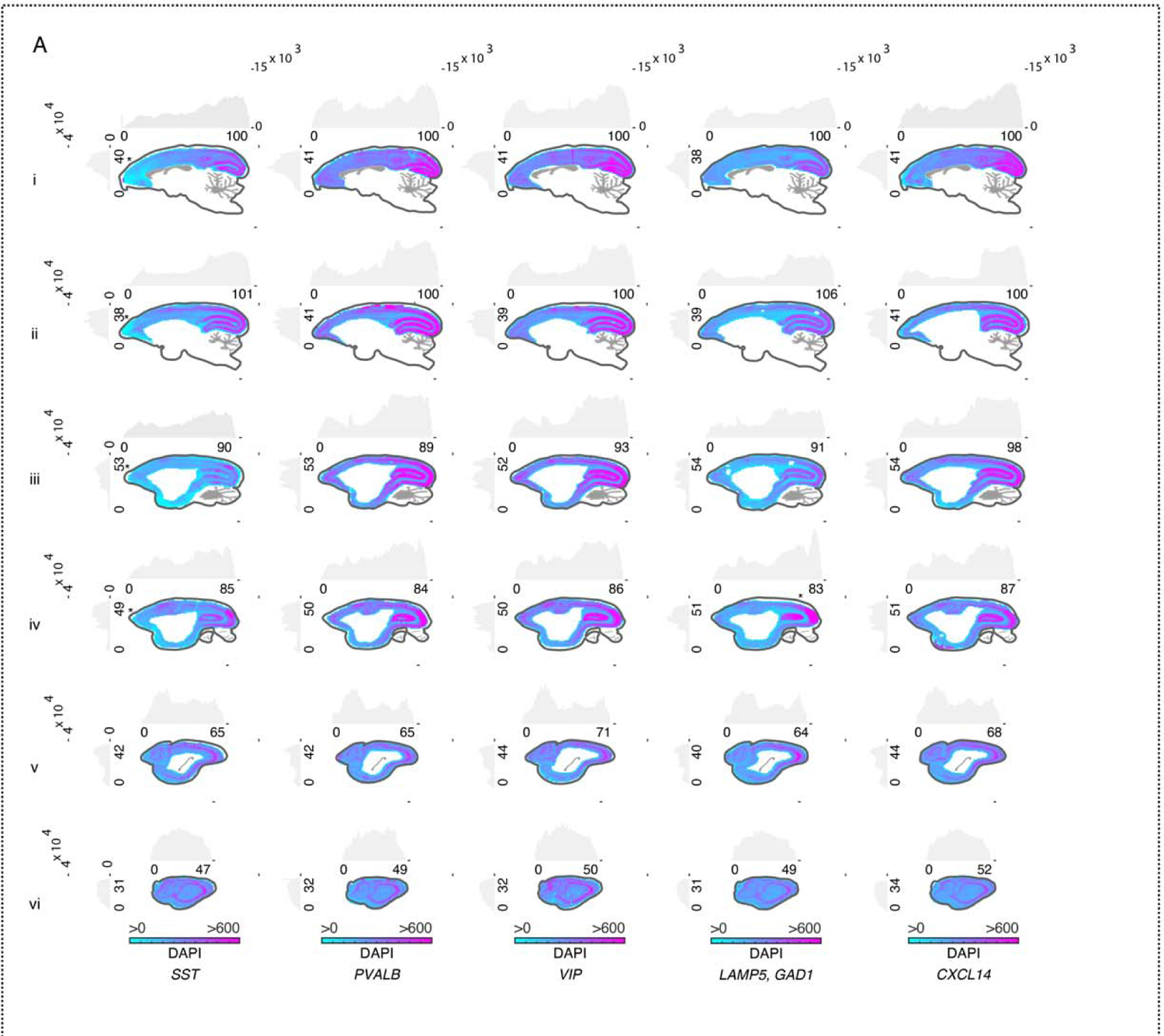
Total cell numbers across marmoset neocortex. **(A)** Total numbers of DAPI+ cells per unit area (approximately 387 µm per bin) for each of the sections shown in Fig. 4C.

**Figure S8.**
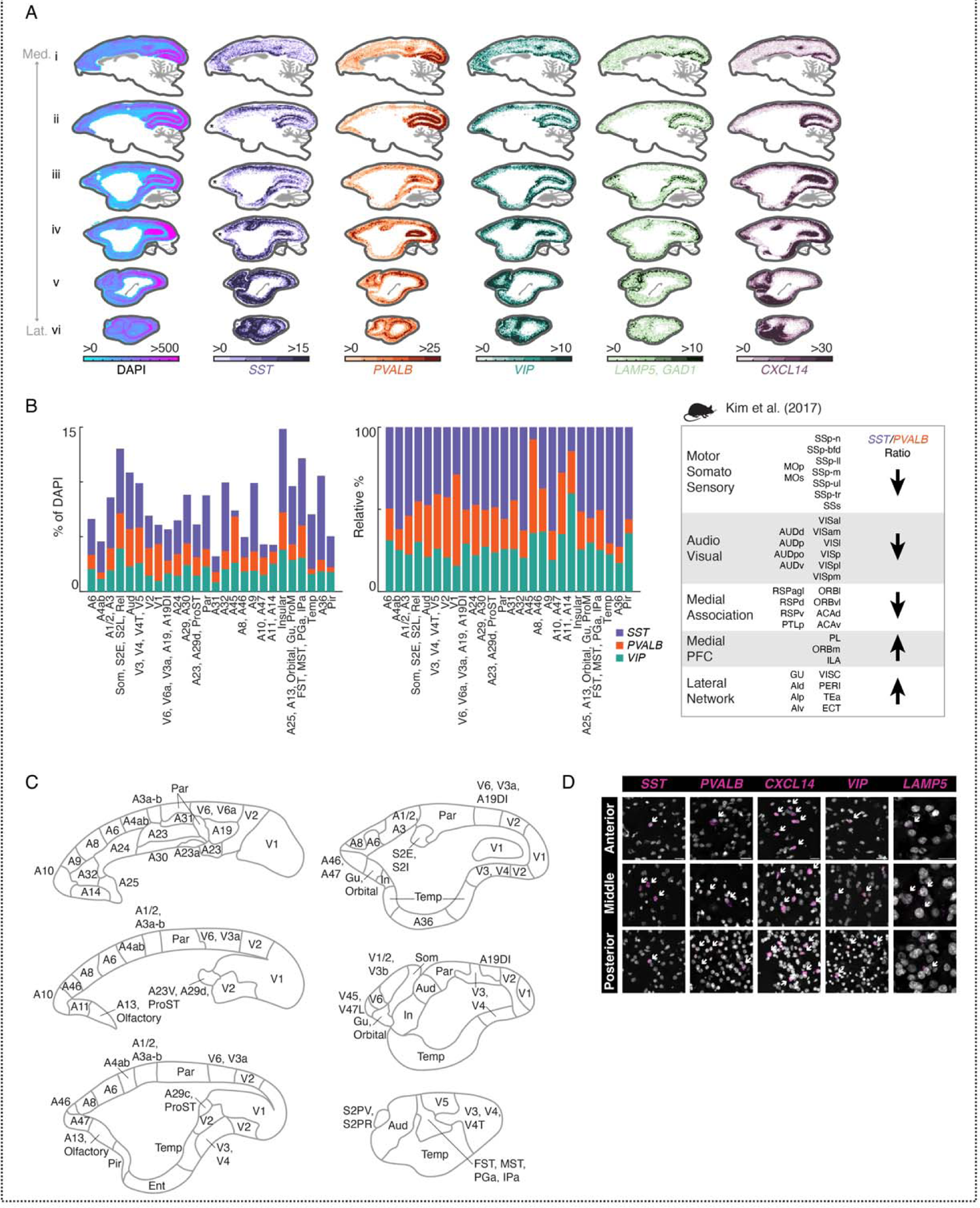
Interneuron numbers across marmoset neocortex. (A) smFISH for neocortical interneuron subclass markers showing locations of cells positive for each marker across 6 sagittal sections of the marmoset neocortex. Heatmap scale shows absolute density per unit area (approximately 387 µm per bin). First column shows DAPI and area profiled. **(B)** (*Left, middle)* Quantitation of interneuron proportions by cortical area in marmoset parcellated according to Fig. S8C. *(Right)* Quantitation of interneuron proportions by cortical area in mouse reproduced from Kim et al. (2017). (*Left*) Absolute percentages of *SST, PVALB,* and *VIP* populations. (*Middle*) Same as (*left),* but scaled as proportions to 100%. (*Right*) Schematic describing ratios of major interneuron types (*Sst+, Pvalb+*) in mouse from Kim et al. (2017). **(C)** Cartoons of cortical areas and areal groupings used to bin smFISH neocortical interneuron proportions in (B) and Fig. 4D-E Neocortical parcellation from https://doi.org/10.24475/bma.4520. **(D)** Examples of smFISH images quantitated in Fig. 4B-E. Panels for each marker show example positive cells in anterior, middle, and posterior locations across neocortex. White arrows indicate positive cells. Scale bar = 20 µm.

**Figure S9.**
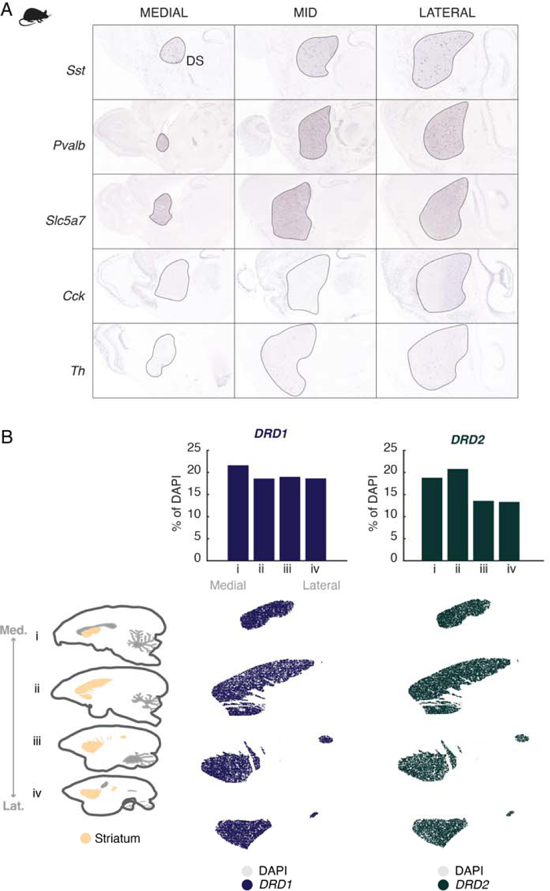
Interneuron numbers across marmoset striatum. (A) Mouse striatal *in situ* from Allen Brain Atlas for *Sst, Pvalb, Slc5a7, Cck, Th*. **(B)** Cartoon of marmoset striatum illustrates area profiled, medial to lateral. Proportions of *DRD1*+ (dark blue) and *DRD2*+ (dark green) cells across primate striatum calculated as in Fig. 4H.

**Figure S10.**
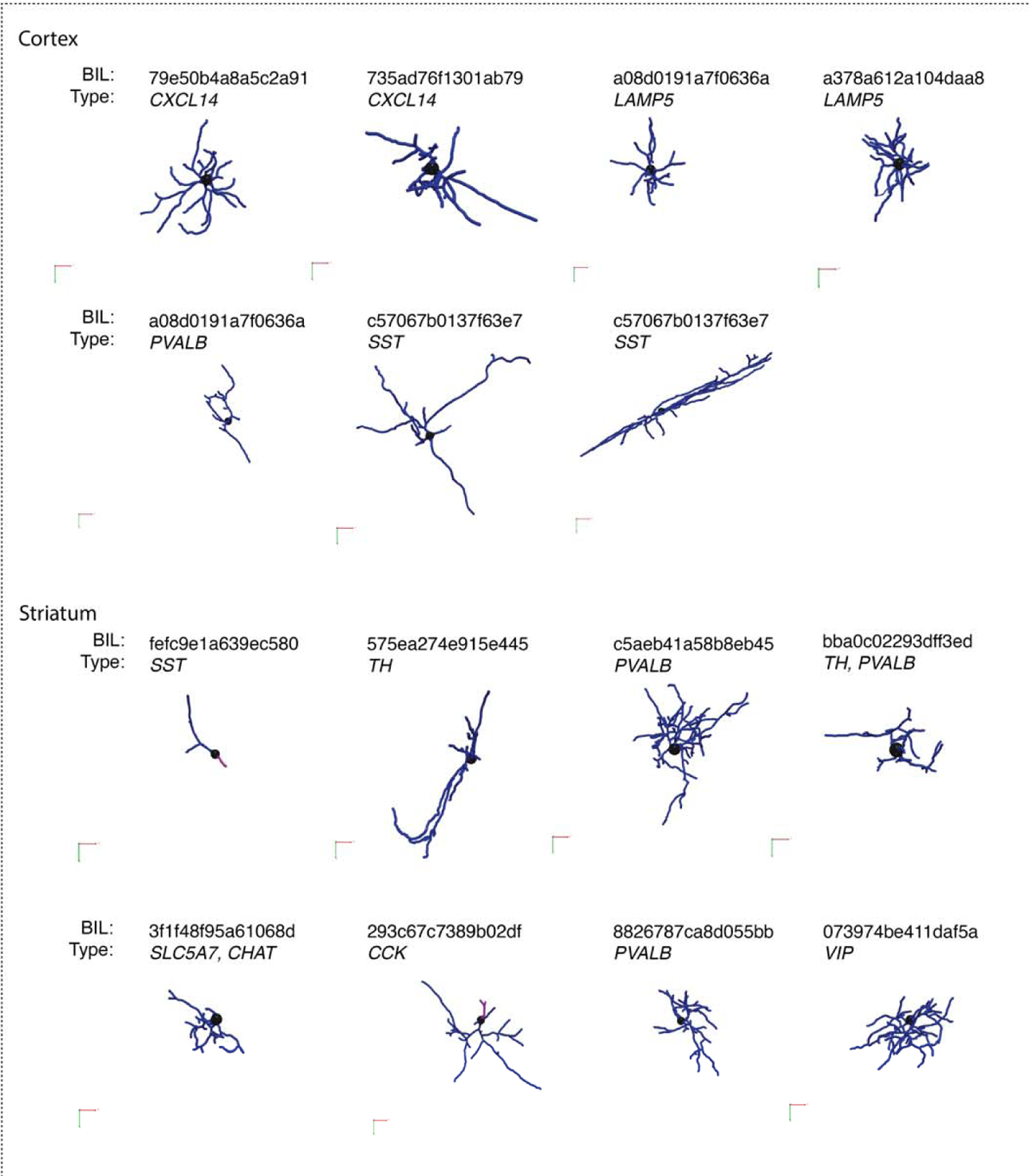
Morphology examples using NeuTube reconstructions. (A) Example morphological reconstructions of striatal and neocortical interneurons using the NeuTube pipeline. Each cell, along with associated smFISH staining, is available for download at https://doi.org/10.35077/g.609.

**Figure S11.**
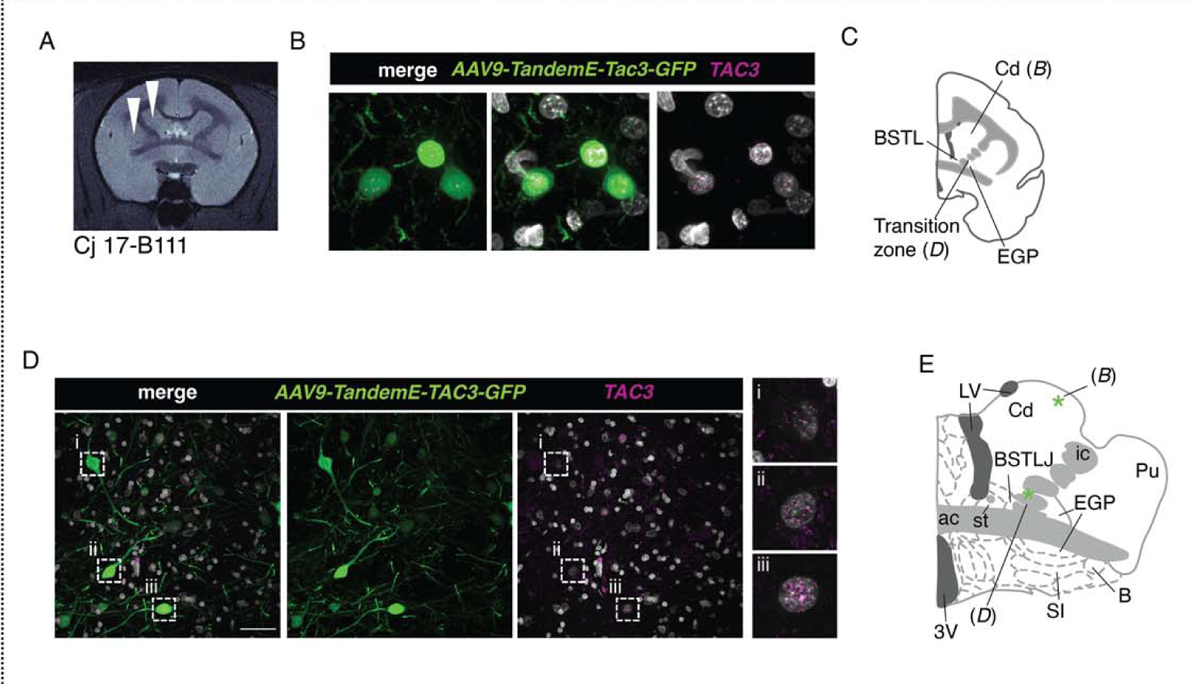
Examples of striatal and peri-striatal *TAC3*+ neurons labeled by AAV-tandemE-TAC3-EGFP. (A) MRI showing injection location (white arrowheads) of virus into bilateral dorsal striatum (caudate) in one animal (Cj 17-B111). **(B)** EGFP antibody-amplified confocal image of a labeled cell (position shown in (C) with smFISH for *TAC3* showing colocalization). **(C)** Cartoon showing location of positive cells in (B) as well as labeled cells in transition zone (D). **(D)** Examples of extra-striatal labeled cells from injections in (A). **(E)** Position of cells in (D). Cd = Caudate, SI = Substantia innominata, B = basal nucleus of Meynert, ac = anterior commissure, BSTLP = bed nuc st, lateral posterior, BSTLJ = bed nuc st, juxtacap, 3V = third ventricle, LV = lateral ventricle.

**Table S1. Marmoset sample information.** Table of animals, metadata, and experimental information.

Table S2. **snRNA-seq dataset by donor and brain area.** Tables of the number of cells per brain structure and samples per donor.

**Table S3. Neocortical rDEGs across three donors.** Neocortical rDEGs across three donors with pairwise comparisons between neocortical locations for major clusters of cortical excitatory neurons, inhibitory neurons, astrocytes and oligodendrocyte lineage types.

Table S4. **Fluorescent *in situ* hybridization probes and antibodies used.** A list of all FISH probes (RNA-Scope and Molecular Instruments) and antibodies used for validation of gene and protein expression *in situ*.

**Table S5. Morphological reconstructions performed with Neutube.** A list of all cells reconstructed with Neutube with their corresponding morphological measurements.

**Table S6. Morphological reconstructions performed with Imaris.** A list of all cells reconstructed with Imaris with their corresponding morphological measurements.

**Table S7. Links and DOIs.** A list of all links and DOIs referenced.

**Data S1. Striatal and cortical medio-lateral quantification.** Tabular data for striatal and cortical medio-lateral histograms/bar plots.

**Data S2. *TAC3* medio-lateral quantification.** Tabular data for *TAC3* medio-lateral histograms and bar plots.

## References

1. S. Ma, M. Skarica, Q. Li, C. Xu, R. D. Risgaard, A. T. N. Tebbenkamp, X. Mato-Blanco, R. Kovner, Ž. Krsnik, X. de Martin, V. Luria, X. Martí-Pérez, D. Liang, A. Karger, D. K. Schmidt, Z. Gomez-Sanchez, C. Qi, K. T. Gobeske, S. Pochareddy, A. Debnath, C. J. Hottman, J. Spurrier, L. Teo, A. G. Boghdadi, J. Homman-Ludiye, J. J. Ely, E. W. Daadi, D. Mi, M. Daadi, O. Marín, P. R. Hof, M.-R. Rasin, J. Bourne, C. C. Sherwood, G. Santpere, M. J. Girgenti, S. M. Strittmatter, A. M. M. Sousa, N. Sestan, Molecular and cellular evolution of the primate dorsolateral prefrontal cortex.*Science*, eabo7257 (2022).

2. T. E. Bakken, N. L. Jorstad, Q. Hu, B. B. Lake, W. Tian, B. E. Kalmbach, M. Crow, R. D. Hodge, F. M. Krienen, S. A. Sorensen, J. Eggermont, Z. Yao, B. D. Aevermann, A. I. Aldridge, A. Bartlett, D. Bertagnolli, T. Casper, R. G. Castanon, K. Crichton, T. L. Daigle, R. Dalley, N. Dee, N. Dembrow, D. Diep, S.-L. Ding, W. Dong, R. Fang, S. Fischer, M. Goldman, J. Goldy, L. T. Graybuck, B. R. Herb, X. Hou, J. Kancherla, M. Kroll, K. Lathia, B. van Lew, Y. E. Li, C. S. Liu, H. Liu, J. D. Lucero, A. Mahurkar, D. McMillen, J. A. Miller, M. Moussa, J. R. Nery, P. R. Nicovich, S.-Y. Niu, J. Orvis, J. K. Osteen, S. Owen, C. R. Palmer, T. Pham, N. Plongthongkum, O. Poirion, N. M. Reed, C. Rimorin, A. Rivkin, W. J. Romanow, A. E. Sedeño-Cortés, K. Siletti, S. Somasundaram, J. Sulc, M. Tieu, A. Torkelson, H. Tung, X. Wang, F. Xie, A. M. Yanny, R. Zhang, S. A. Ament, M. M. Behrens, H. C. Bravo, J. Chun, A. Dobin, J. Gillis, R. Hertzano, P. R. Hof, T. Höllt, G. D. Horwitz, C. D. Keene, P. V. Kharchenko, A. L. Ko, B. P. Lelieveldt, C. Luo, E. A. Mukamel, A. Pinto-Duarte, S. Preissl, A. Regev, B. Ren, R. H. Scheuermann, K. Smith, W. J. Spain, O. R. White, C. Koch, M. Hawrylycz, B. Tasic, E. Z. Macosko, S. A. McCarroll, J. T. Ting, H. Zeng, K. Zhang, G. Feng, J. R. Ecker, S. Linnarsson, E. S. Lein, Comparative cellular analysis of motor cortex in human, marmoset and mouse. Nature. 598, 111–119 (2021).

3. F. M. Krienen, M. Goldman, Q. Zhang, R. C. H. del Rosario, M. Florio, R. Machold, A. Saunders, K. Levandowski, H. Zaniewski, B. Schuman, C. Wu, A. Lutservitz, C. D. Mullally, N. Reed, E. Bien, L. Bortolin, M. Fernandez-Otero, J. D. Lin, A. Wysoker, J. Nemesh, D. Kulp, M. Burns, V. Tkachev, R. Smith, C. A. Walsh, J. Dimidschstein, B. Rudy, L. S. Kean, S. Berretta, G. Fishell, G. Feng, S. A. McCarroll, Innovations present in the primate interneuron repertoire. Nature. 586, 262– 269 (2020).

4. J.-P. Lin, H. M. Kelly, Y. Song, R. Kawaguchi, D. H. Geschwind, S. Jacobson, D. S. Reich, Microenvironment Impacts the Molecular Architecture and Interactivity of Resident Cells in Marmoset Brain. *bioRxiv* (2021), p. 2021.01.25.426385.

5. R. C. Bandler, I. Vitali, R. N. Delgado, M. C. Ho, E. Dvoretskova, J. S. Ibarra Molinas, P. W. Frazel, M. Mohammadkhani, R. Machold, S. Maedler, S. A. Liddelow, T. J. Nowakowski, G. Fishell, C. Mayer, Single-cell delineation of lineage and genetic identity in the mouse brain. Nature. 601, 404– 409 (2022).

6. A. Saunders, E. Z. Macosko, A. Wysoker, M. Goldman, F. M. Krienen, H. de Rivera, E. Bien, M. Baum, L. Bortolin, S. Wang, A. Goeva, J. Nemesh, N. Kamitaki, S. Brumbaugh, D. Kulp, S. A. McCarroll, Molecular Diversity and Specializations among the Cells of the Adult Mouse Brain. Cell. 174, 1015–1030.e16 (2018).

7. A. Zeisel, H. Hochgerner, P. Lönnerberg, A. Johnsson, F. Memic, J. van der Zwan, M. Häring, E. Braun, L. E. Borm, G. La Manno, S. Codeluppi, A. Furlan, K. Lee, N. Skene, K. D. Harris, J. Hjerling-Leffler, E. Arenas, P. Ernfors, U. Marklund, S. Linnarsson, Molecular Architecture of the Mouse Nervous System. Cell. 174, 999–1014.e22 (2018).

8. Z. Yao, C. T. J. van Velthoven, M. Kunst, M. Zhang, D. McMillen, C. Lee, W. Jung, J. Goldy, A. Abdelhak, P. Baker, E. Barkan, D. Bertagnolli, J. Campos, D. Carey, T. Casper, A. B. Chakka, R. Chakrabarty, S. Chavan, M. Chen, M. Clark, J. Close, K. Crichton, S. Daniel, T. Dolbeare, L. Ellingwood, J. Gee, A. Glandon, J. Gloe, J. Gould, J. Gray, N. Guilford, J. Guzman, D. Hirschstein, W. Ho, K. Jin, M. Kroll, K. Lathia, A. Leon, B. Long, Z. Maltzer, N. Martin, R. McCue, E. Meyerdierks, T. N. Nguyen, T. Pham, C. Rimorin, A. Ruiz, N. Shapovalova, C. Slaughterbeck, J. Sulc, M. Tieu, A. Torkelson, H. Tung, N. V. Cuevas, K. Wadhwani, K. Ward, B. Levi, C. Farrell, C. L. Thompson, S. Mufti, C. M. Pagan, L. Kruse, N. Dee, S. M. Sunkin, L. Esposito, M. J. Hawrylycz, J. Waters, L. Ng, K. A. Smith, B. Tasic, X. Zhuang, H. Zeng, A high-resolution transcriptomic and spatial atlas of cell types in the whole mouse brain. bioRxiv (2023), p. 2023.03.06.531121.

9. J. Langlieb, N. Sachdev, K. Balderrama, N. Nadaf, M. Raj, E. Murray, J. Webber, C. Vanderburg, V. Gazestani, D. Tward, C. Mezias, X. Li, D. Cable, T. Norton, P. P. Mitra, F. Chen, E. Macosko, The cell type composition of the adult mouse brain revealed by single cell and spatial genomics. bioRxiv (2023), doi:10.1101/2023.03.06.531307.

10. K. L. Chiou, X. Huang, M. O. Bohlen, S. Tremblay, D. R. O’Day, C. H. Spurrell, A. A. Gogate, T. M. Zintel, Cayo Biobank Research Unit, M. G. Andrews, M. I. Martinez, L. M. Starita, M. J. Montague, M. L. Platt, J. Shendure, N. Snyder-Mackler, A single-cell multi-omic atlas spanning the adult rhesus macaque brain. bioRxiv (2022), p. 2022.09.30.510346.

11. K. Siletti, R. Hodge, A. M. Albiach, L. Hu, K. W. Lee, P. Lönnerberg, T. Bakken, S.-L. Ding, M. Clark, T. Casper, N. Dee, J. Gloe, C. Dirk Keene, J. Nyhus, H. Tung, A. M. Yanny, E. Arenas, E. S. Lein, S. Linnarsson, Transcriptomic diversity of cell types across the adult human brain. bioRxiv (2022), p. 2022.10.12.511898.

12. J.-P. Lin, H. M. Kelly, Y. Song, R. Kawaguchi, D. H. Geschwind, S. Jacobson, D. S. Reich, Transcriptomic architecture of nuclei in the marmoset CNS. Nat. Commun. 13, 5531 (2022).

13. R. D. Hodge, T. E. Bakken, J. A. Miller, K. A. Smith, E. R. Barkan, L. T. Graybuck, J. L. Close, B. Long, N. Johansen, O. Penn, Z. Yao, J. Eggermont, T. Hollt, B. P. Levi, S. I. Shehata, B. Aevermann, A. Beller, D. Bertagnolli, K. Brouner, T. Casper, C. Cobbs, R. Dalley, N. Dee, S.-L. Ding, R. G. Ellenbogen, O. Fong, E. Garren, J. Goldy, R. P. Gwinn, D. Hirschstein, C. D. Keene, M. Keshk, A. L. Ko, K. Lathia, A. Mahfouz, Z. Maltzer, M. McGraw, T. N. Nguyen, J. Nyhus, J. G. Ojemann, A. Oldre, S. Parry, S. Reynolds, C. Rimorin, N. V. Shapovalova, S. Somasundaram, A. Szafer, E. R. Thomsen, M. Tieu, G. Quon, R. H. Scheuermann, R. Yuste, S. M. Sunkin, B. Lelieveldt, D. Feng, L. Ng, A. Bernard, M. Hawrylycz, J. W. Phillips, B. Tasic, H. Zeng, A. R. Jones, C. Koch, E. S. Lein, Conserved cell types with divergent features in human versus mouse cortex. Nature. 573, 1–38 (2019).

14. S. Sahara, Y. Yanagawa, D. D. M. O’Leary, C. F. Stevens, The Fraction of Cortical GABAergic Neurons Is Constant from Near the Start of Cortical Neurogenesis to Adulthood. Journal of Neuroscience. 32, 4755–4761 (2012).

15. Y. Kim, G. R. Yang, K. Pradhan, K. U. Venkataraju, M. Bota, L. C. G. del Molino, G. Fitzgerald, K. Ram, M. He, J. M. Levine, P. Mitra, Z. J. Huang, X.-J. Wang, P. Osten, Brain-wide Maps Reveal Stereotyped Cell-Type-Based Cortical Architecture and Subcortical Sexual Dimorphism. Cell. 171, 456–460.e22 (2017).

16. J. Hill, T. Inder, J. Neil, D. Dierker, J. Harwell, D. Van Essen, Similar patterns of cortical expansion during human development and evolution. Proceedings of the National Academy of Sciences. 107, 13135–13140 (2010).

17. D. Goertsen, N. C. Flytzanis, N. Goeden, M. R. Chuapoco, A. Cummins, Y. Chen, Y. Fan, Q. Zhang, J. Sharma, Y. Duan, L. Wang, G. Feng, Y. Chen, N. Y. Ip, J. Pickel, V. Gradinaru, AAV capsid variants with brain-wide transgene expression and decreased liver targeting after intravenous delivery in mouse and marmoset. Nat. Neurosci. 25, 106–115 (2022).

18. J. K. Mich, L. T. Graybuck, E. E. Hess, J. T. Mahoney, Y. Kojima, Y. Ding, S. Somasundaram, J. A. Miller, B. E. Kalmbach, C. Radaelli, B. B. Gore, N. Weed, V. Omstead, Y. Bishaw, N. V. Shapovalova, R. A. Martinez, O. Fong, S. Yao, M. Mortrud, P. Chong, L. Loftus, D. Bertagnolli, J. Goldy, T. Casper, N. Dee, X. Opitz-Araya, A. Cetin, K. A. Smith, R. P. Gwinn, C. Cobbs, A. L. Ko, J. G. Ojemann, C. D. Keene, D. L. Silbergeld, S. M. Sunkin, V. Gradinaru, G. D. Horwitz, H. Zeng, B. Tasic, E. S. Lein, J. T. Ting, B. P. Levi, Functional enhancer elements drive subclass-selective expression from mouse to primate neocortex. Cell Rep. 34, 108754 (2021).

19. J. Dimidschstein, Q. Chen, R. Tremblay, S. L. Rogers, G.-A. Saldi, L. Guo, Q. Xu, R. Liu, C. Lu, J. Chu, J. S. Grimley, A.-R. Krostag, A. Kaykas, M. C. Avery, M. S. Rashid, M. Baek, A. L. Jacob, G. B. Smith, D. E. Wilson, G. Kosche, I. Kruglikov, T. Rusielewicz, V. C. Kotak, T. M. Mowery, S. A. Anderson, E. M. Callaway, J. S. Dasen, D. Fitzpatrick, V. Fossati, M. A. Long, S. Noggle, J. H. Reynolds, D. H. Sanes, B. Rudy, G. Feng, G. Fishell, A viral strategy for targeting and manipulating interneurons across vertebrate species. Nat. Neurosci. 19, 1743–1749 (2016).

20. D. Vormstein-Schneider, J. D. Lin, K. A. Pelkey, R. Chittajallu, B. Guo, M. A. Arias-Garcia, K. Allaway, S. Sakopoulos, G. Schneider, O. Stevenson, J. Vergara, J. Sharma, Q. Zhang, T. P. Franken, J. Smith, L. A. Ibrahim, K. J. M. Astro, E. Sabri, S. Huang, E. Favuzzi, T. Burbridge, Q. Xu, L. Guo, I. Vogel, V. Sanchez, G. A. Saldi, B. L. Gorissen, X. Yuan, K. A. Zaghloul, O. Devinsky, B. L. Sabatini, R. Batista-Brito, J. Reynolds, G. Feng, Z. Fu, C. J. McBain, G. Fishell, J. Dimidschstein, Viral manipulation of functionally distinct interneurons in mice, non-human primates and humans. Nat. Neurosci. 23, 1–21 (2020).

21. J. Alquicira-Hernandez, A. Sathe, H. P. Ji, Q. Nguyen, J. E. Powell, scPred: accurate supervised method for cell-type classification from single-cell RNA-seq data. Genome Biol. 20, 264 (2019).

22. H. Zeng, J. R. Sanes, Neuronal cell-type classification: challenges, opportunities and the path forward. Nat. Rev. Neurosci. 18, 530–546 (2017).

23. D. Arendt, J. M. Musser, C. V. H. Baker, A. Bergman, C. Cepko, D. H. Erwin, M. Pavlicev, G. Schlosser, S. Widder, M. D. Laubichler, G. P. Wagner, The origin and evolution of cell types. Nat. Rev. Genet. 17, 744–757 (2016).

24. H. Zeng, What is a cell type and how to define it? Cell (2022) (available at https://www.sciencedirect.com/science/article/pii/S0092867422007838).

25. L. J. Revell phytools: an R package for phylogenetic comparative biology (and other things). Methods in Ecology and Evolution. 3, 217–223 (2012).

26. J. Woych, A. Ortega Gurrola, A. Deryckere, E. C. B. Jaeger, E. Gumnit, G. Merello, J. Gu, A. Joven Araus, N. D. Leigh, M. Yun, A. Simon, M. A. Tosches, Cell-type profiling in salamanders identifies innovations in vertebrate forebrain evolution. Science. 377, eabp9186 (2022).

27. G. Meyer, Building a human cortex: the evolutionary differentiation of Cajal-Retzius cells and the cortical hem. J. Anat. 217, 334–343 (2010).

28. P. Jager, G. Moore, P. Calpin, X. Durmishi, I. Salgarella, L. Menage, Y. Kita, Y. Wang, D. W. Kim, S. Blackshaw, S. R. Schultz, S. Brickley, T. Shimogori, A. Delogu, Dual midbrain and forebrain origins of thalamic inhibitory interneurons. Elife. 10 (2021), doi:10.7554/eLife.59272.

29. K. Letinic, P. Rakic, Telencephalic origin of human thalamic GABAergic neurons. Nat. Neurosci. 4, 931–936 (2001).

30. L. W. Swanson, G. D. Petrovich, What is the amygdala? Trends Neurosci. 21, 323–331 (1998).

31. T. Hirata, P. Li, G. M. Lanuza, L. A. Cocas, M. M. Huntsman, J. G. Corbin, Identification of distinct telencephalic progenitor pools for neuronal diversity in the amygdala. Nat. Neurosci. 12, 141–149 (2009).

32. T. Kaoru, F. C. Liu, M. Ishida, T. Oishi, M. Hayashi, M. Kitagawa, K. Shimoda, H. Takahashi, Molecular characterization of the intercalated cell masses of the amygdala: implications for the relationship with the striatum. NSC. 166, 220–230 (2010).

33. M. T. Schmitz, K. Sandoval, C. P. Chen, M. A. Mostajo-Radji, W. W. Seeley, T. J. Nowakowski, C. J. Ye, M. F. Paredes, A. A. Pollen, The development and evolution of inhibitory neurons in primate cerebrum. Nature. 603, 871–877 (2022).

34. D. L. Kaufman, C. R. Houser, A. J. Tobin, Two forms of the gamma-aminobutyric acid synthetic enzyme glutamate decarboxylase have distinct intraneuronal distributions and cofactor interactions. J. Neurochem. 56, 720–723 (1991).

35. J. R. Moffitt, D. Bambah-Mukku, S. W. Eichhorn, E. Vaughn, K. Shekhar, J. D. Perez, N. D. Rubinstein, J. Hao, A. Regev, C. Dulac, X. Zhuang, Molecular, spatial, and functional single-cell profiling of the hypothalamic preoptic region. Science. 362, eaau5324–80 (2018).

36. B. Tasic, Z. Yao, L. T. Graybuck, K. A. Smith, T. N. Nguyen, D. Bertagnolli, J. Goldy, E. Garren, M. N. Economo, S. Viswanathan, O. Penn, T. Bakken, V. Menon, J. Miller, O. Fong, K. E. Hirokawa, K. Lathia, C. Rimorin, M. Tieu, R. Larsen, T. Casper, E. Barkan, M. Kroll, S. Parry, N. V. Shapovalova, D. Hirschstein, J. Pendergraft, H. A. Sullivan, T. K. Kim, A. Szafer, N. Dee, P. Groblewski, I. Wickersham, A. Cetin, J. A. Harris, B. P. Levi, S. M. Sunkin, L. Madisen, T. L. Daigle, L. Looger, A. Bernard, J. Phillips, E. Lein, M. Hawrylycz, K. Svoboda, A. R. Jones, C. Koch, H. Zeng, Shared and distinct transcriptomic cell types across neocortical areas. Nature. 563, 1–41 (2018).

37. P. Balaram, J. H. Kaas, Towards a unified scheme of cortical lamination for primary visual cortex across primates: insights from NeuN and VGLUT2 immunoreactivity. Front. Neuroanat. 8, 81 (2014).

38. O. A. Bayraktar, L. C. Fuentealba, A. Alvarez-Buylla, D. H. Rowitch, Astrocyte development and heterogeneity. Cold Spring Harb. Perspect. Biol. 7, a020362 (2014).

39. O. A. Bayraktar, T. Bartels, S. Holmqvist, V. Kleshchevnikov, A. Martirosyan, D. Polioudakis, L. Ben Haim, A. M. H. Young, M. Y. Batiuk, K. Prakash, A. Brown, K. Roberts, M. F. Paredes, R. Kawaguchi, J. H. Stockley, K. Sabeur, S. M. Chang, E. Huang, P. Hutchinson, E. M. Ullian, M. Hemberg, G. Coppola, M. G. Holt, D. H. Geschwind, D. H. Rowitch, Astrocyte layers in the mammalian cerebral cortex revealed by a single-cell in situ transcriptomic map. Nat. Neurosci. 23, 500–509 (2020).

40. M. Y. Batiuk, A. Martirosyan, J. Wahis, F. de Vin, C. Marneffe, C. Kusserow, J. Koeppen, J. F. Viana, J. F. Oliveira, T. Voet, C. P. Ponting, T. G. Belgard, M. G. Holt, Identification of region-specific astrocyte subtypes at single cell resolution. Nat. Commun. 11, 1220 (2020).

41. A. Kriegstein, A. Alvarez-Buylla, The glial nature of embryonic and adult neural stem cells. Annu. Rev. Neurosci. 32, 149–184 (2009).

42. C. C. Harwell, L. C. Fuentealba, A. Gonzalez-Cerrillo, P. R. L. Parker, C. C. Gertz, E. Mazzola, M. T. Garcia, A. Alvarez-Buylla, C. L. Cepko, A. R. Kriegstein, Wide Dispersion and Diversity of Clonally Related Inhibitory Interneurons. Neuron. 87, 999–1007 (2015).

43. C. Mayer, C. Hafemeister, R. C. Bandler, R. Machold, R. B. Brito, X. Jaglin, K. Allaway, A. Butler, G. Fishell, R. Satija, Developmental diversification of cortical inhibitory interneurons. Nature. 555, 457–462 (2018).

44. C. Neyt, M. Welch, A. Langston, J. Kohtz, G. Fishell, A short-range signal restricts cell movement between telencephalic proliferative zones. J. Neurosci. 17, 9194–9203 (1997).

45. F. García-Moreno, M. Pedraza, L. G. Di Giovannantonio, M. Di Salvio, L. López-Mascaraque, A. Simeone, J. A. De Carlos, A neuronal migratory pathway crossing from diencephalon to telencephalon populates amygdala nuclei. Nat. Neurosci. 13, 680–689 (2010).

46. A. J. Granger, M. L. Wallace, B. L. Sabatini, ScienceDirect Multi-transmitter neurons in the mammalian central nervous system. Curr. Opin. Neurobiol. 45, 85–91 (2017).

47. N. X. Tritsch, A. J. Granger, B. L. Sabatini, Mechanisms and functions of GABA co-release. Nat. Genet. 17, 139–145 (2016).

48. M. Bupesh, I. Legaz, A. Abellán, L. Medina, Multiple telencephalic and extratelencephalic embryonic domains contribute neurons to the medial extended amygdala. J. Comp. Neurol. 519, 1505–1525 (2011).

49. L. Zhang, V. S. Hernandez, C. R. Gerfen, S. Z. Jiang, L. Zavala, R. A. Barrio, L. E. Eiden, Behavioral role of PACAP signaling reflects its selective distribution in glutamatergic and GABAergic neuronal subpopulations. Elife. 10 (2021), doi:10.7554/eLife.61718.

50. N. Bozadjieva-Kramer, R. A. Ross, D. Q. Johnson, H. Fenselau, D. L. Haggerty, B. Atwood, B. Lowell, J. N. Flak, The Role of Mediobasal Hypothalamic PACAP in the Control of Body Weight and Metabolism. Endocrinology. 162 (2021), doi:10.1210/endocr/bqab012.

51. J.-H. Cho, K. Zushida, G. P. Shumyatsky, W. A. Carlezon Jr, E. G. Meloni, V. Y. Bolshakov, Pituitary adenylate cyclase-activating polypeptide induces postsynaptically expressed potentiation in the intra-amygdala circuit. J. Neurosci. 32, 14165–14177 (2012).

52. E. Y. Lee, L. C. Chan, H. Wang, J. Lieng, M. Hung, Y. Srinivasan, J. Wang, J. A. Waschek, A. L. Ferguson, K.-F. Lee, N. Y. Yount, M. R. Yeaman, G. C. L. Wong, PACAP is a pathogen-inducible resident antimicrobial neuropeptide affording rapid and contextual molecular host defense of the brain. Proc. Natl. Acad. Sci. U. S. A. 118 (2021), doi:10.1073/pnas.1917623117.

53. Y. Kita, H. Nishibe, Y. Wang, T. Hashikawa, S. S. Kikuchi, M. U A. C. Yoshida, C. Yoshida, T. Kawase, S. Ishii, H. Skibbe, T. Shimogori, Cellular-resolution gene expression profiling in the neonatal marmoset brain reveals dynamic species- and region-specific differences. Proc. Natl. Acad. Sci. U. S. A. 118 (2021), doi:10.1073/pnas.2020125118.

54. T. Shimogori, A. Abe, Y. Go, T. Hashikawa, N. Kishi, S. S. Kikuchi, Y. Kita, K. Niimi, H. Nishibe, M. Okuno, K. Saga, M. Sakurai, M. Sato, T. Serizawa, S. Suzuki, E. Takahashi, M. Tanaka, S. Tatsumoto, M. Toki, M. U Y. Wang, K. J. Windak, H. Yamagishi, K. Yamashita, T. Yoda, A. C. Yoshida, C. Yoshida, T. Yoshimoto, H. Okano, Digital gene atlas of neonate common marmoset brain. Neurosci. Res. 128, 1–13 (2018).

55. D. P. Orquera, M. B. Tavella, F. S. J. de Souza, S. Nasif, M. J. Low, M. Rubinstein, The Homeodomain Transcription Factor NKX2.1 Is Essential for the Early Specification of Melanocortin Neuron Identity and Activates Pomc Expression in the Developing Hypothalamus. J. Neurosci. 39, 4023–4035 (2019).

56. R. Murcia-Ramón, V. Company, I. Juárez-Leal, A. Andreu-Cervera, F. Almagro-García, S. Martínez, D. Echevarría, E. Puelles, Neuronal tangential migration from Nkx2.1-positive hypothalamus. Brain Struct. Funct. 225, 2857–2869 (2020).

57. C. E. Collins, D. C. Airey, N. A. Young, D. B. Leitch, J. H. Kaas, Neuron densities vary across and within cortical areas in primates. Proc. Natl. Acad. Sci. U. S. A. 107, 15927–15932 (2010).

58. B. Schuman, R. P. Machold, Y. Hashikawa, J. Fuzik, G. J. Fishell, B. Rudy, Four Unique Interneuron Populations Reside in Neocortical Layer 1. J. Neurosci. 39, 125–139 (2019).

59. M. Valero, T. J. Viney, R. Machold, S. Mederos, I. Zutshi, B. Schuman, Y. Senzai, B. Rudy, G. Buzsáki, Sleep down state-active ID2/Nkx2.1 interneurons in the neocortex. Nat. Neurosci. 24, 401– 411 (2021).

60. B. Zingg, H. Hintiryan, L. Gou, M. Y. Song, M. Bay, M. S. Bienkowski, N. N. Foster, S. Yamashita, I. Bowman, A. W. Toga, H.-W. Dong, Neural networks of the mouse neocortex. Cell. 156, 1096– 1111 (2014).

61. H. B. M. Uylings, H. J. Groenewegen, B. Kolb, Do rats have a prefrontal cortex? Behav. Brain Res. 146, 3–17 (2003).

62. R. L. Buckner, D. S. Margulies, Macroscale cortical organization and a default-like apex transmodal network in the marmoset monkey. Nat. Commun. 10, 1976 (2019).

63. M. K. Lin, Y. S. Takahashi, B.-X. Huo, M. Hanada, J. Nagashima, J. Hata, A. S. Tolpygo, K. Ram, B. C. Lee, M. I. Miller, M. G. Rosa, E. Sasaki, A. Iriki, H. Okano, P. Mitra, A high-throughput neurohistological pipeline for brain-wide mesoscale connectivity mapping of the common marmoset. Elife. 8 (2019), doi:10.7554/eLife.40042.

64. P. Majka, S. Bai, S. Bakola, S. Bednarek, J. M. Chan, N. Jermakow, L. Passarelli, D. H. Reser, P. Theodoni, K. H. Worthy, X.-J. Wang, D. K. Wójcik, P. P. Mitra, M. G. P. Rosa, Open access resource for cellular-resolution analyses of corticocortical connectivity in the marmoset monkey. Nat. Commun. 11, 1133 (2020).

65. A. Hladnik, D. Džaja, S. Darmopil, N. Jovanov-Milošević, Z. Petanjek, Spatio-temporal extension in site of origin for cortical calretinin neurons in primates. Front. Neuroanat. 8, 50 (2014).

66. E. Y. Choi, B. T. T. Yeo, R. L. Buckner, The organization of the human striatum estimated by intrinsic functional connectivity. J. Neurophysiol. 108, 2242–2263 (2012).

67. K. M. Anderson, F. M. Krienen, E. Y. Choi, J. M. Reinen, B. T. T. Yeo, A. J. Holmes, Gene expression links functional networks across cortex and striatum. Nat. Commun. 9, 1428 (2018).

68. M. Matamales, J. Götz, J. Bertran-Gonzalez, Quantitative Imaging of Cholinergic Interneurons Reveals a Distinctive Spatial Organization and a Functional Gradient across the Mouse Striatum. PLoS One. 11, e0157682 (2016).

69. E. Fino, M. Vandecasteele, S. Perez, F. Saudou, L. Venance, Region-specific and state-dependent action of striatal GABAergic interneurons. Nat. Commun. 9, 3339 (2018).

70. A. B. M. Manchado, C. B. Gonzales, A. Zeisel, H. Munguba, B. Bekkouche, N. G. Skene, P. Lonnerberg, J. Ryge, K. D. Harris, S. Linnarsson, J. Hjerling-Leffler, Diversity of Interneurons in the Dorsal Striatum Revealed by Single-Cell RNA Sequencing and PatchSeq. Cell Rep. 24, 2179– 2190.e7 (2018).

71. T. Zerucha, T. Stühmer, G. Hatch, B. K. Park, Q. Long, G. Yu, A. Gambarotta, J. R. Schultz, J. L. Rubenstein, M. Ekker, A highly conserved enhancer in the Dlx5/Dlx6 intergenic region is the site of cross-regulatory interactions between Dlx genes in the embryonic forebrain. J. Neurosci. 20, 709– 721 (2000).

72. T. Stuart, A. Srivastava, S. Madad, C. A. Lareau, R. Satija, Single-cell chromatin state analysis with Signac. Nat. Methods. 18, 1333–1341 (2021).

73. N. L. Jorstad, J. H. T. Song, D. Exposito-Alonso, H. Suresh, N. Castro, F. M. Krienen, A. M. Yanny, J. Close, E. Gelfand, K. J. Travaglini, S. Basu, M. Beaudin, D. Bertagnolli, M. Crow, S.-L. Ding, J. Eggermont, A. Glandon, J. Goldy, T. Kroes, B. Long, D. McMillen, T. Pham, C. Rimorin, K. Siletti, S. Somasundaram, M. Tieu, A. Torkelson, K. Ward, G. Feng, W. D. Hopkins, T. Höllt, C. Dirk Keene, S. Linnarsson, S. A. McCarroll, B. P. Lelieveldt, C. C. Sherwood, K. Smith, C. A. Walsh, A. Dobin, J. Gillis, E. S. Lein, R. D. Hodge, T. E. Bakken, Comparative transcriptomics reveals human-specific cortical features. *bioRxiv* (2022), p. 2022.09.19.508480.

74. D. Velmeshev, L. Schirmer, D. Jung, M. Haeussler, Y. Perez, S. Mayer, A. Bhaduri, N. Goyal, D. H. Rowitch, A. R. Kriegstein, Single-cell genomics identifies cell type–specific molecular changes in autism. Science. 364, 685–689 (2019).

75. W. B. Ruzicka, S. Mohammadi, J. F. Fullard, J. Davila-Velderrain, S. Subburaju, D. R. Tso, M. Hourihan, S. Jiang, H.-C. Lee, J. Bendl, G. Voloudakis, V. Haroutunian, G. E. Hoffman, P. Roussos, M. Kellis, PsychENCODE Consortium, Single-cell multi-cohort dissection of the schizophrenia transcriptome. bioRxiv (2022),, doi:10.1101/2022.08.31.22279406.

76. C. Nagy, M. Maitra, A. Tanti, M. Suderman, J.-F. Théroux, M. A. Davoli, K. Perlman, V. Yerko, Y. C. Wang, S. J. Tripathy, P. Pavlidis, N. Mechawar, J. Ragoussis, G. Turecki, Single-nucleus transcriptomics of the prefrontal cortex in major depressive disorder implicates oligodendrocyte precursor cells and excitatory neurons. Nat. Neurosci. 23, 771–781 (2020).

77. A. Sequeira, F. Mamdani, C. Ernst, M. P. Vawter, W. E. Bunney, V. Lebel, S. Rehal, T. Klempan, A. Gratton, C. Benkelfat, G. A. Rouleau, N. Mechawar, G. Turecki, Global brain gene expression analysis links glutamatergic and GABAergic alterations to suicide and major depression. PLoS One. 4, e6585 (2009).

78. D. Arendt, P. Y. Bertucci, K. Achim, J. M. Musser, Evolution of neuronal types and families. Curr. Opin. Neurobiol. 56, 144–152 (2019).

79. R. N. Delgado, D. E. Allen, M. G. Keefe, W. R. Mancia Leon, R. S. Ziffra, E. E. Crouch, A. Alvarez-Buylla, T. J. Nowakowski, Individual human cortical progenitors can produce excitatory and inhibitory neurons. Nature. 601, 397–403 (2022).

80. B. Martynoga, H. Morrison, D. J. Price, J. O. Mason, Foxg1 is required for specification of ventral telencephalon and region-specific regulation of dorsal telencephalic precursor proliferation and apoptosis. Dev. Biol. 283, 113–127 (2005).

81. J. M. Tepper, F. Tecuapetla, T. Koós, O. Ibáñez-Sandoval, Heterogeneity and diversity of striatal GABAergic interneurons. Front. Neuroanat. 4, 150 (2010).

82. J. DeFelipe, P. L. López-Cruz, R. Benavides-Piccione, C. Bielza, P. Larrañaga, S. Anderson, A. Burkhalter, B. Cauli, A. Fairén, D. Feldmeyer, G. Fishell, D. Fitzpatrick, T. F. Freund, G. González-Burgos, S. Hestrin, S. Hill, P. R. Hof, J. Huang, E. G. Jones, Y. Kawaguchi, Z. Kisvárday, Y. Kubota, D. A. Lewis, O. Marín, H. Markram, C. J. McBain, H. S. Meyer, H. Monyer, S. B. Nelson, K. Rockland, J. Rossier, J. L. R. Rubenstein, B. Rudy, M. Scanziani, G. M. Shepherd, C. C. Sherwood, J. F. Staiger, G. Tamás, A. Thomson, Y. Wang, R. Yuste, G. A. Ascoli, New insights into the classification and nomenclature of cortical GABAergic interneurons. Nat. Rev. Neurosci. 14, 202–216 (2013).

83. R. Tremblay, S. Lee, B. Rudy, GABAergic Interneurons in the Neocortex: From Cellular Properties to Circuits. Neuron. 91, 260–292 (2016).

84. E. Boldog, T. E. Bakken, R. D. Hodge, M. Novotny, B. D. Aevermann, J. Baka, S. Bordé, J. L. Close, F. Diez-Fuertes, S.-L. Ding, N. Faragó, Á. K. Kocsis, B. Kovács, Z. Maltzer, J. M. McCorrison, J. A. Miller, G. Molnár, G. Oláh, A. Ozsvár, M. Rózsa, S. I. Shehata, K. A. Smith, S. M. Sunkin, D. N. Tran, P. Venepally, A. Wall, L. G. Puskás, P. Barzó, F. J. Steemers, N. J. Schork, R. H. Scheuermann, R. S. Lasken, E. S. Lein, G. Tamás, Transcriptomic and morphophysiological evidence for a specialized human cortical GABAergic cell type. Nat. Neurosci. 21, 1185–1195 (2018).

85. A. M. M. Sousa, Y. Zhu, M. A. Raghanti, R. R. Kitchen, M. Onorati, A. T. N. Tebbenkamp, B. Stutz, K. A. Meyer, M. Li, Y. I. Kawasawa, F. Liu, R. G. Perez, M. Mele, T. Carvalho, M. Skarica, F. O. Gulden, M. Pletikos, A. Shibata, A. R. Stephenson, M. K. Edler, J. J. Ely, J. D. Elsworth, T. L. Horvath, P. R. Hof, T. M. Hyde, J. E. Kleinman, D. R. Weinberger, M. Reimers, R. P. Lifton, S. M. Mane, J. P. Noonan, State, Matthew W, E. S. Lein, J. A. Knowles, T. Marques-Bonet, C. C. Sherwood, M. B. Gerstein, N. Šestan, Molecular and cellular reorganization of neural circuits in the human lineage. Science. 358, 1027–1032 (2017).

86. G. Paxinos, C. Watson, M. Petrides, M. Rosa, H. Tokuno, The marmoset brain in stereotaxic coordinates (2012).

87. F. M. Krienen, M. Goldman, Q. Zhang, R. C H Del Rosario, M. Florio, R. Machold, A. Saunders, K. Levandowski, H. Zaniewski, B. Schuman, C. Wu, A. Lutservitz, C. D. Mullally, N. Reed, E. Bien, L. Bortolin, M. Fernandez-Otero, J. D. Lin, A. Wysoker, J. Nemesh, D. Kulp, M. Burns, V. Tkachev, R. Smith, C. A. Walsh, J. Dimidschstein, B. Rudy, L. S Kean, S. Berretta, G. Fishell, G. Feng, S. A. McCarroll, Innovations present in the primate interneuron repertoire. Nature. 586, 262–269 (2020).

88. S. J. Fleming, M. D. Chaffin, A. Arduini, A.-D. Akkad, E. Banks, J. C. Marioni, A. A. Philippakis, P. T. Ellinor, M. Babadi, Unsupervised removal of systematic background noise from droplet-based single-cell experiments using CellBender. *bioRxiv* (2019), doi:10.1101/791699.

89. C. S. McGinnis, L. M. Murrow, Z. J. Gartner, DoubletFinder: Doublet Detection in Single-Cell RNA Sequencing Data Using Artificial Nearest Neighbors. Cell Syst. 8, 329–337.e4 (2019).

90. L. Feng, T. Zhao, J. Kim, neuTube 1.0: A New Design for Efficient Neuron Reconstruction Software Based on the SWC Format. eNeuro. 2 (2015), doi:10.1523/ENEURO.0049-14.2014.

91. R. C. Challis, S. Ravindra Kumar, K. Y. Chan, C. Challis, K. Beadle, M. J. Jang, H. M. Kim, P. S. Rajendran, J. D. Tompkins, K. Shivkumar, B. E. Deverman, V. Gradinaru, Systemic AAV vectors for widespread and targeted gene delivery in rodents. Nat. Protoc. 14, 379–414 (2019).

