## Supplementary material for "A marmoset brain cell census reveals influence of developmental origin and functional class on neuronal identity"

**Supplementary Materials:**

Figs. S1-S11
Table S1-S7
Data S1-S2

### Supp S1

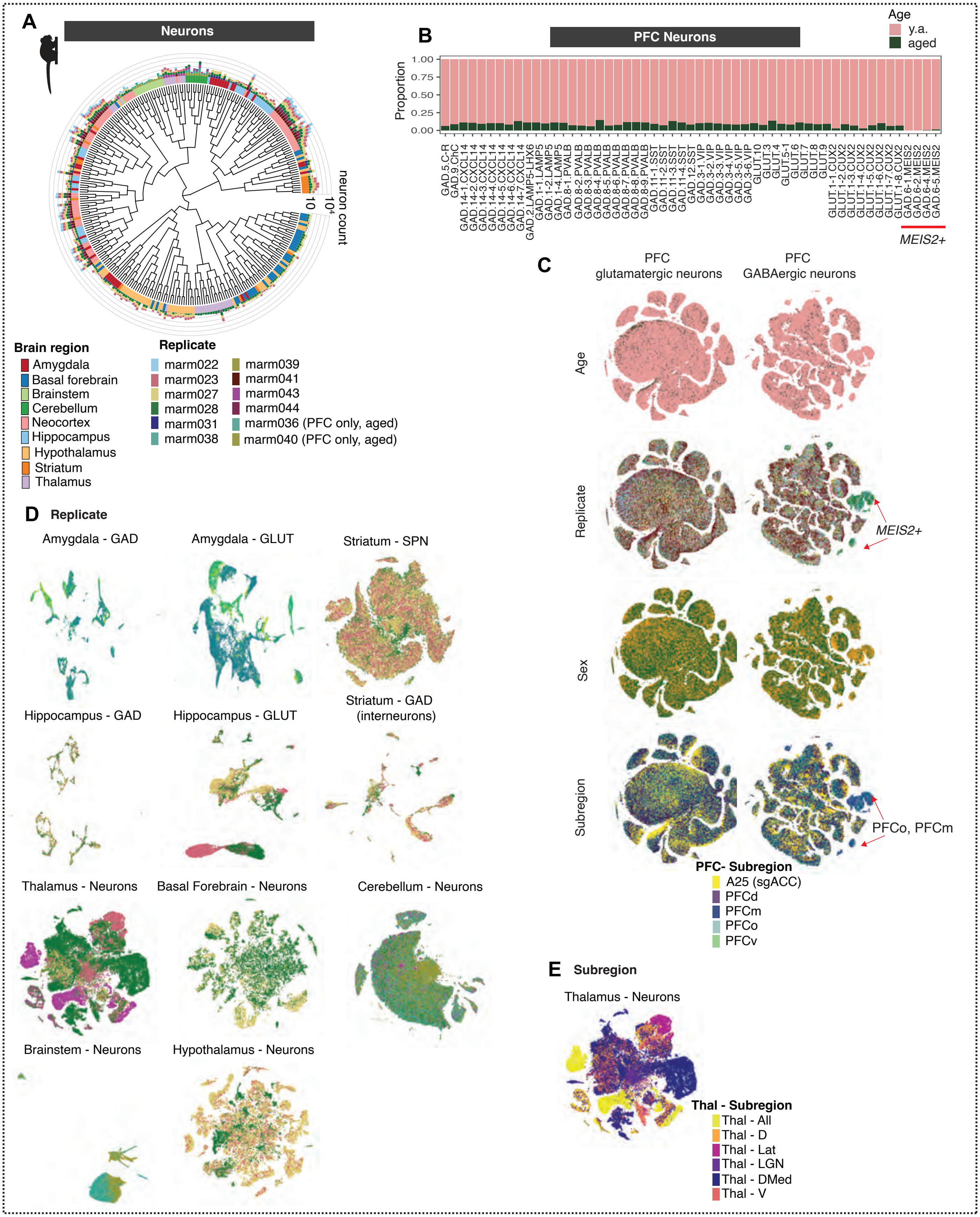

A
Neurons
neuron count
10
10^4
Brain region
Amygdala
Basal forebrain
Brainstem
Cerebellum
Neocortex
Hippocampus
Hypothalamus
Striatum
Thalamus
Replicate
marm022
marm023
marm027
marm028
marm031
marm038
marm039
marm041
marm043
marm044
marm036 (PFC only, aged)
marm040 (PFC only, aged)
B
PFC Neurons
Age
y.a.
aged
Proportion
1.00
0.75
0.50
0.25
0.00
GAD.5.C-R
GAD.9.ChC
GAD.14-1.CXCL14
GAD.14-2.CXCL14
GAD.14-3.CXCL14
GAD.14-4.CXCL14
GAD.14-5.CXCL14
GAD.14-6.CXCL14
GAD.14-7.CXCL14
GAD.2.LAMP5-LHX6
GAD.1-1.LAMP5
GAD.1-2.LAMP5
GAD.1-4.LAMP5
GAD.8-1.PVALB
GAD.8-2.PVALB
GAD.8-3.PVALB
GAD.8-4.PVALB
GAD.8-5.PVALB
GAD.8-6.PVALB
GAD.8-7.PVALB
GAD.8-8.PVALB
GAD.8-9.PVALB
GAD.11-1.SST
GAD.11-2.SST
GAD.11-3.SST
GAD.11-4.SST
GAD.12.SST
GAD.3-1.VIP
GAD.3-2.VIP
GAD.3-3.VIP
GAD.3-4.VIP
GAD.3-5.VIP
GAD.3-6.VIP
GLUT.10
GLUT.3
GLUT.4
GLUT.5-1
GLUT.6
GLUT.7
GLUT.8
GLUT.9
GLUT.1-1.CUX2
GLUT.1-2.CUX2
GLUT.1-3.CUX2
GLUT.1-4.CUX2
GLUT.1-5.CUX2
GLUT.1-6.CUX2
GLUT.1-7.CUX2
GLUT.1-8.CUX2
GAD.6-1.MEIS2
GAD.6-2.MEIS2
GAD.6-4.MEIS2
GAD.6-5.MEIS2
MEIS2+
C
PFC glutamatergic neurons
PFC GABAergic neurons
Age
Replicate
MEIS2+
Sex
Subregion
PFCo, PFCm
PFC- Subregion
A25 (sgACC)
PFCd
PFCm
PFCo
PFCv
D
Replicate
Amygdala - GAD
Amygdala - GLUT
Striatum - SPN
Hippocampus - GAD
Hippocampus - GLUT
Striatum - GAD (interneurons)
Thalamus - Neurons
Basal Forebrain - Neurons
Cerebellum - Neurons
Brainstem - Neurons
Hypothalamus - Neurons
E
Subregion
Thalamus - Neurons
Thal - Subregion
Thal - All
Thal - D
Thal - Lat
Thal - LGN
Thal - DMed
Thal - V

**Figure S1. Neuron counts by donor across brain regions. (A)** Neuronal dendrogram as in **Fig. 1C**, with outer barplots depicting number of nuclei per cell type and replicate. Ring colors are brain regions, colors in barplots correspond to replicates. **(B)** Proportional per-cluster representation of PFC neurons between young adult donors and aged (n=2) donors. While our snRNA-seq collection focused on post-sexual maturity young adults, we acquired an additional dataset of PFC sampled from 2 aged animals (1 M, 11y0m; 1F, 14y4m, 37,260 cells total; **Table S1**). Individual replicates contributed similar proportions of neurons to each prefrontal neuron subtype, and clusters generally had proportional representation across young adults and aged animals, as well as across males and females (**Fig. S1B-C**), suggesting that these variables do not dramatically impact neuronal ensembles and identities in prefrontal cortex. **(C)** t-SNE embeddings of PFC neurons (*top row*, GABAergic; *bottom row*, glutamatergic) with colors representing different metadata: age (young vs aged), replicate, PFC subregion. There was notable enrichment of *MEIS2*+ GABAergic neurons in medial prefrontal and orbital prefrontal dissections **(Fig. S1C)**. Based on their gene expression profiles, these cells likely correspond to the recently described population of LGE-derived *MEIS2*+ neurons that populate the olfactory bulb in mice, and which are instead directed to medial prefrontal cortex in macaques and humans (*33*). **(D)** t-SNEs of neurons in each brain structure, with cells colored by replicate (colors as in (A)). Telencephalic neurons are plotted separately by class: GABAergic and glutamatergic classes (neocortex, hippocampus, amygdala), or GABAergic interneurons and spiny projection neurons (striatum). Compared with neocortex, greater cross-donor variability was observed in some subcortical structures such as hypothalamus and thalamus, though this was likely driven more by dissection variability than by donor variability, as the donor-specific clusters tended to be from subregions that were only sampled in one individual (see E). **(E)** t-SNEs of thalamic neurons with cells colored by thalamic subdivision.

Supp S2

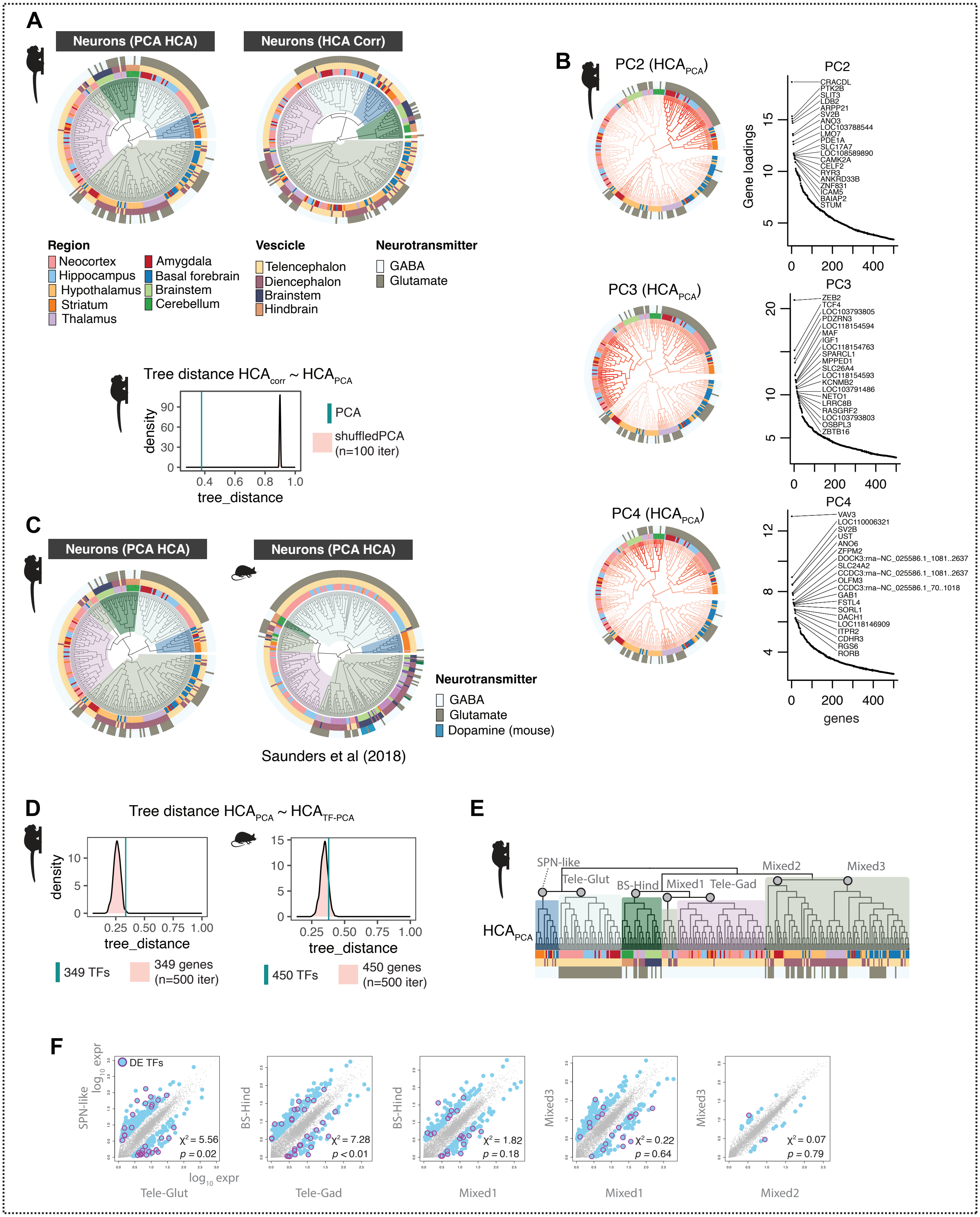
A
Neurons (PCA HCA)
Neurons (HCA Corr)
Region
Neocortex
Hippocampus
Hypothalamus
Striatum
Thalamus
Amygdala
Basal forebrain
Brainstem
Cerebellum
Vescicle
Telencephalon
Diencephalon
Brainstem
Hindbrain
Neurotransmitter
GABA
Glutamate
Tree distance HCA_corr ~ HCA_PCA
density
tree_distance
PCA
shuffledPCA (n=100 iter)
B
PC2 (HCA_PCA)
PC3 (HCA_PCA)
PC4 (HCA_PCA)
Gene loadings
genes
C
Neurons (PCA HCA)
Neurons (PCA HCA)
Neurotransmitter
GABA
Glutamate
Dopamine (mouse)
Saunders et al (2018)
D
Tree distance HCA_PCA ~ HCA_TF-PCA
density
tree_distance
349 TFs
349 genes (n=500 iter)
450 TFs
450 genes (n=500 iter)
E
HCA_PCA
SPN-like
Tele-Glut
BS-Hind
Mixed1
Tele-Gad
Mixed2
Mixed3
F
DE TFs
log10 expr
SPN-like
Tele-Glut
χ² = 5.56
p = 0.02
BS-Hind
Tele-Gad
χ² = 7.28
p < 0.01
BS-Hind
Mixed1
χ² = 1.82
p = 0.18
Mixed3
Mixed1
χ² = 0.22
p = 0.64
Mixed3
Mixed2
χ² = 0.07
p = 0.79

**Figure S2. Conservation of neuronal hierarchy across species and clustering methods. (A)** Comparison of marmoset and mouse neurons (mouse atlas data from (*6*)) using three different distance calculations for hierarchical clustering: $HCA_{corr}$ distance = gene expression correlations of top 5904 genes in marmoset or 3528 in mouse (mouse genes include all genes with 1:1 orthologs to the 5904 marmoset genes and that are expressed in at least 10 transcripts per 100,000 in at least one mouse neuron type); $HCA_{PCA}$ distance = top 100 PCA scores across the same genes; $HCA_{TF\text{-}PCA}$ distance = top 100 PCA scores using expressed transcription factors only (marmoset = 349 TFs, mouse = 450 TFs). **(B)** Distance between hierarchical clustering (HC) dendrogram trees computed using different methods. Cyan line = tree distance (R package TreeDist) between hierarchical clustering using distance = $HCA_{corr}$ and using distance = $HCA_{PCA}$. Pink distribution is tree distance scores between $HCA_{corr}$ and shuffled PCA scores (n = 100 shuffling iterations). Lower values of tree_distance (x-axis) mean higher agreement between dendrogram tree structures. **(C)** PCA loadings and top genes for PC2-PC4. PC scores are plotted on the $HCA_{PCA}$ dendrogram. Ranked gene loading plots show top 20 genes per PC. **(D)** Tree distances computed as in (B) between $HCA_{PCA}$ and $HCA_{TF\text{-}PCA}$. The tree distances between these two trees is low, but not different from distributions of random, same-sized sets of genes. **(E)** Marmoset dendrogram in **Fig. 1C** ($HCA_{PCA}$ ) indicating major clades compared in (F). **(F)** Ancestral reconstruction (AR; R package phytools) of gene expression profiles of major clades of marmoset neuron types from dendrogram in (E). Maximum likelihood estimates of gene expression (fastAnc) were computed for 7 major internal nodes (gray circles) of the $HCA_{PCA}$ dendrogram. Scatterplots show pairwise comparisons between AR of internal nodes of major clades. Blue dots = genes with >3 foldchange difference between the two ARs. Magenta circles = differentially expressed transcription factors (DE-TFs). Chi-square and p-values describe whether TFs are significantly differentially enriched between the AR pairs.

Supp S3

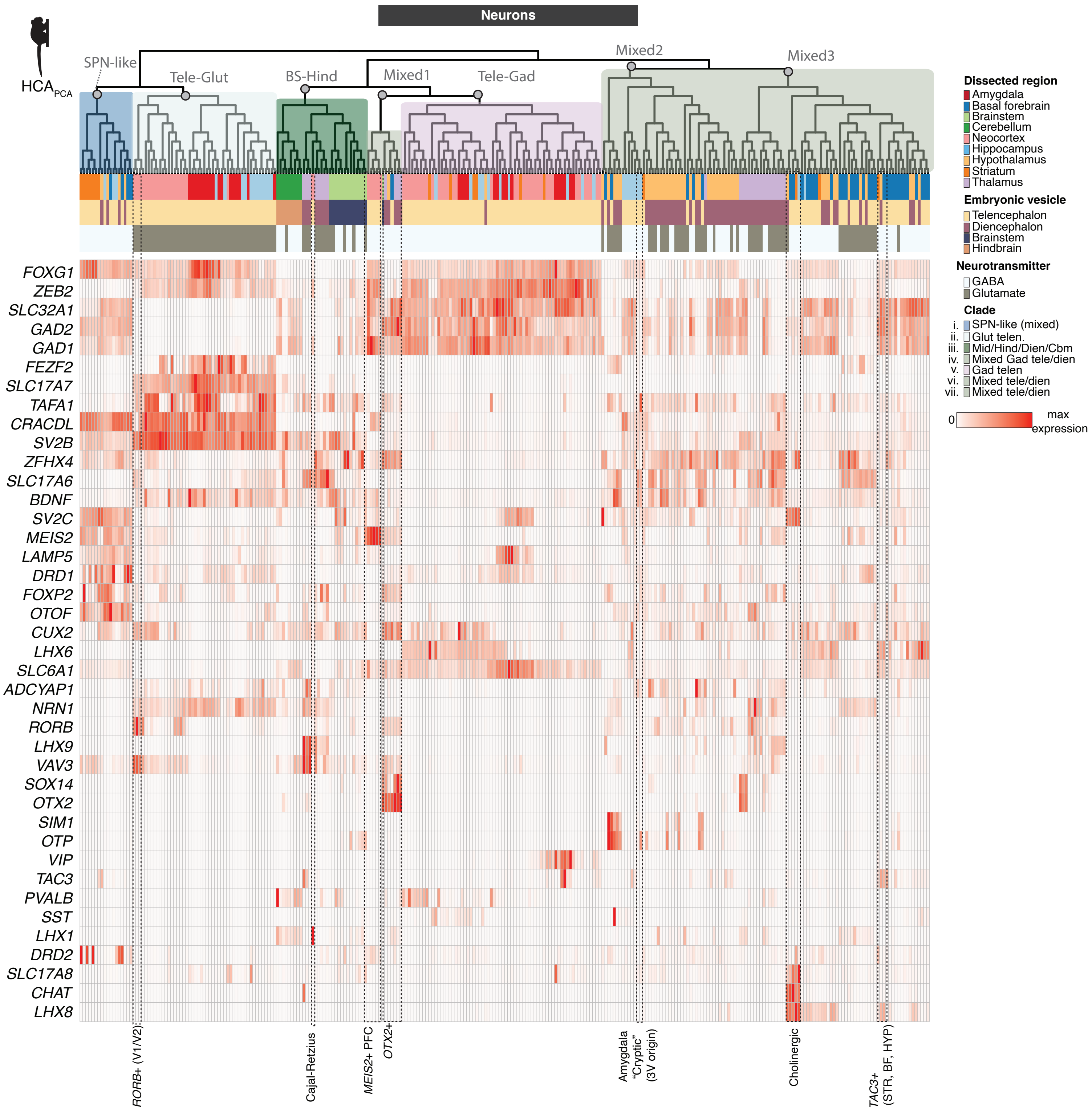
Neurons
HCA$_{PCA}$
SPN-like
Tele-Glut
BS-Hind
Mixed1
Tele-Gad
Mixed2
Mixed3
Dissected region
Amygdala
Basal forebrain
Brainstem
Cerebellum
Neocortex
Hippocampus
Hypothalamus
Striatum
Thalamus
Embryonic vesicle
Telencephalon
Diencephalon
Brainstem
Hindbrain
Neurotransmitter
GABA
Glutamate
Clade
i. SPN-like (mixed)
ii. Glut telen.
iii. Mid/Hind/Dien/Cbm
iv. Mixed Gad tele/dien
v. Gad telen
vi. Mixed tele/dien
vii. Mixed tele/dien
0
max expression
FOXG1
ZEB2
SLC32A1
GAD2
GAD1
FEZF2
SLC17A7
TAFA1
CRACDL
SV2B
ZFHX4
SLC17A6
BDNF
SV2C
MEIS2
LAMP5
DRD1
FOXP2
OTOF
CUX2
LHX6
SLC6A1
ADCYAP1
NRN1
RORB
LHX9
VAV3
SOX14
OTX2
SIM1
OTP
VIP
TAC3
PVALB
SST
LHX1
DRD2
SLC17A8
CHAT
LHX8
RORB+ (V1/V2)
Cajal-Retzius
MEIS2+ PFC
OTX2+
Amygdala "Cryptic" (3V origin)
Cholinergic
TAC3+ (STR, BF, HYP)

**Figure S3. Gene expression across neural populations.** Expression of broad class marker genes and other genes of interest across all neurons sampled by snRNA-seq. Heatmap colors are scaled to max normalized expression for each row (gene). Dendrogram ordering and metadata colors as in **Fig. 1C**. Cell types discussed in the main text are labeled at bottom.

Supp S4 related to Fig 2

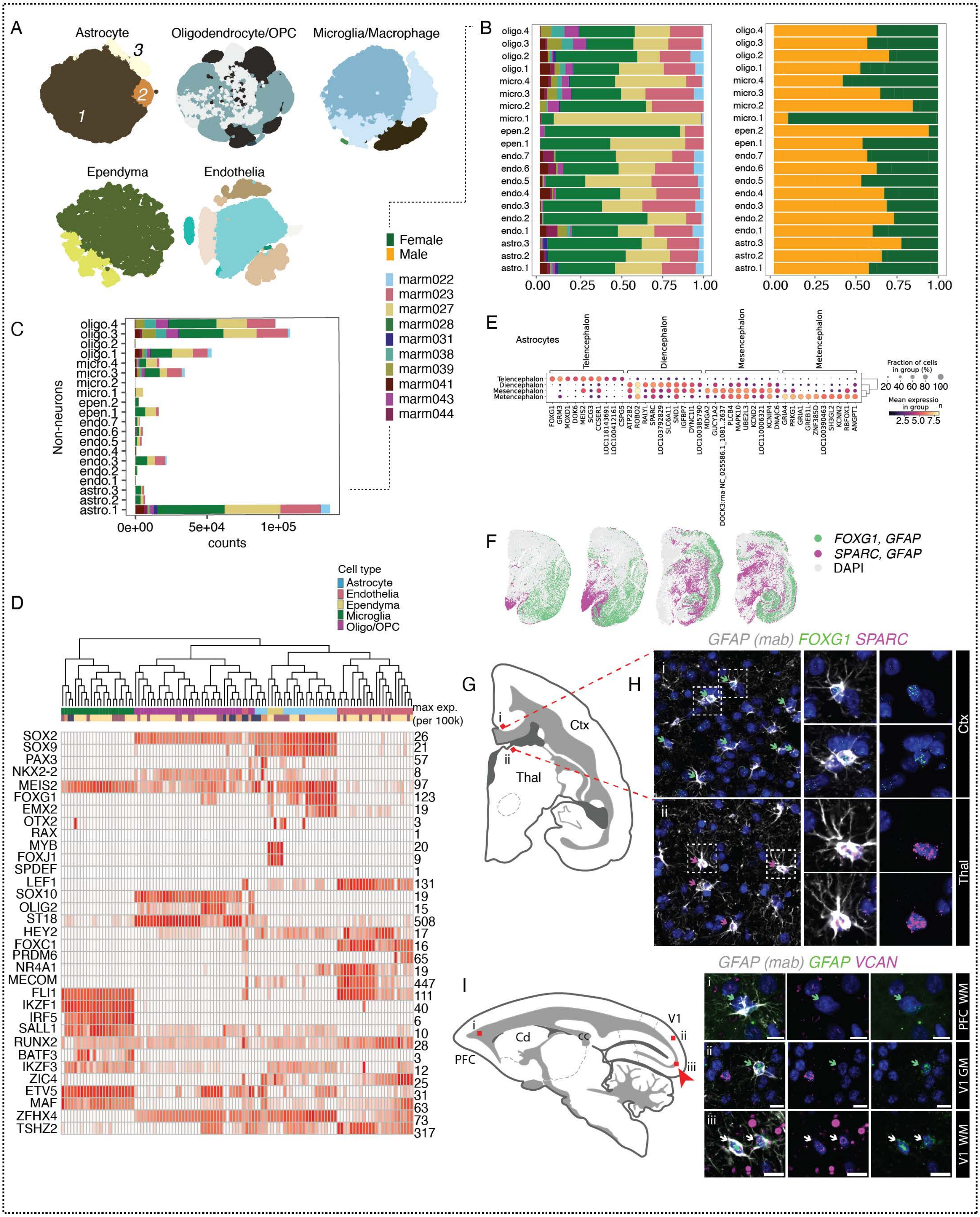

**Figure S4. Glia diversity across regions. (A)** t-SNE embeddings of major non-neuronal types colored by cluster. **(B)** Barplots of glial proportions colored by donor and by sex. **(C)** Non-neuronal nuclei counts. Colors indicate donor, same as (B). **(D)** Expression of marker genes in non-neurons. Genes as in (*8*). Heatmap colors are scaled to max normalized expression for each row (gene). Dendrogram and metadata colors as in **Fig. 1G**. **(E)** Differentially expressed genes in astrocytes across cephalic compartments. **(F)** Tissue validation for astrocyte differentially expressed genes (*FOXG1*, *SPARC*) in coronal sections of marmoset brain. Green dots indicate locations of cells that stain positive for *GFAP* (IHC, mAb **Table S4**) and *FOXG1* (smFISH). Magenta dots indicate cell positions for *GFAP* (IHC) and *SPARC* (smFISH). **(G)** Cartoon of coronal section imaged; Red boxes (*i*-*ii*) correspond to tissue validation in (H). **(H)** (*Left)* Fields of view from neocortex and thalamus stained for *GFAP* antibody (gray), *FOXG1* (green), and *SPARC* (magenta). Green arrows highlight *GFAP* cells colocalized with *FOXG1*, magenta arrows highlight *GFAP* cells colocalized with *SPARC.* (*Right*) Magnified examples of double positive cells in neocortex and thalamus. Ctx = cortex, Thal = thalamus. **(I)** (*Left*) Cartoon of sagittal section imaged; red boxes (*i*-*iii*) correspond to (*Right*) tissue validation of increased abundance of *VCAN*+ astrocytes in adult marmoset V1-adjacent white matter (*iii*) compared with PFC-adjacent white matter (*i*) and V1 gray matter (*ii*). GFAP antibody (gray) combined with smFISH probes against *VCAN* (magenta) and *GFAP* (green). Green arrows correspond to *GFAP*+ (antibody), *GFAP*+ (smFISH) cells. White arrows correspond to *GFAP*+ (antibody), *GFAP*+ (smFISH), and *VCAN*+ cells. V1 = visual cortex V1, PFC = prefrontal cortex, GM = gray matter, WM = white matter. Red arrow highlights locale of *VCAN*+ *GFAP*+ images. Scale bar = 10 µm.

### Supp S5 related to Fig 2

A

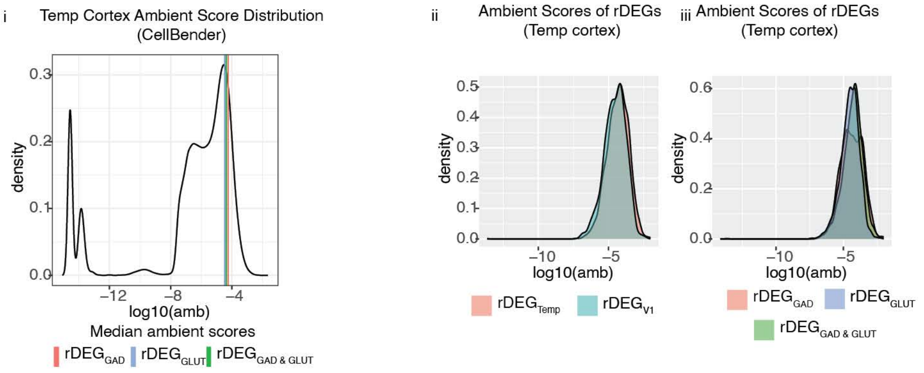


B

**Neuron$_{GAD}$**

| | pfc | m1 | temp | a1 | s1 | par | v2 | v1 |
|---|---|---|---|---|---|---|---|---|
| **pfc** | 0 | 27 | 2 | 6 | 86 | 34 | 101 | 231 |
| **m1** | 5 | 0 | 5 | 1 | 1 | 2 | 2 | 24 |
| **temp** | 25 | 33 | 0 | 86 | 102 | 117 | 155 | 352 |
| **a1** | 16 | 22 | 13 | 0 | 77 | 70 | 97 | 190 |
| **s1** | 57 | 6 | 36 | 25 | 0 | 171 | 108 | 189 |
| **par** | 19 | 9 | 15 | 10 | 89 | 0 | 33 | 181 |
| **v2** | 27 | 1 | 36 | 10 | 18 | 5 | 0 | 54 |
| **v1** | 72 | 14 | 109 | 27 | 53 | 62 | 19 | 0 |

**Neuron$_{GLUT}$**

| | pfc | m1 | temp | a1 | s1 | par | v2 | v1 |
|---|---|---|---|---|---|---|---|---|
| **pfc** | 0 | 118 | 65 | 136 | 202 | 292 | 401 | 413 |
| **m1** | 80 | 0 | 132 | 37 | 9 | 47 | 90 | 108 |
| **temp** | 96 | 242 | 0 | 28 | 96 | 173 | 321 | 364 |
| **a1** | 152 | 98 | 16 | 0 | 111 | 254 | 314 | 368 |
| **s1** | 154 | 35 | 55 | 44 | 0 | 272 | 308 | 358 |
| **par** | 131 | 36 | 87 | 53 | 105 | 0 | 74 | 280 |
| **v2** | 201 | 38 | 175 | 111 | 89 | 31 | 0 | 82 |
| **v1** | 211 | 57 | 189 | 120 | 150 | 141 | 58 | 0 |

**Oligodendrocyte lineage**

| | pfc | m1 | temp | a1 | s1 | par | v2 | v1 |
|---|---|---|---|---|---|---|---|---|
| **pfc** | 0 | 39 | 13 | 3 | 65 | 17 | 81 | 90 |
| **m1** | 15 | 0 | 7 | 9 | 12 | 3 | 5 | 3 |
| **temp** | 37 | 57 | 0 | 41 | 103 | 47 | 25 | 41 |
| **a1** | 201 | 10 | 2 | 0 | 121 | 62 | 76 | 128 |
| **s1** | 186 | 5 | 3 | 14 | 0 | 79 | 88 | 99 |
| **par** | 153 | 2 | 1 | 16 | 121 | 0 | 10 | 99 |
| **v2** | 191 | 13 | 4 | 40 | 96 | 34 | 0 | 83 |
| **v1** | 141 | 6 | 9 | 37 | 84 | 45 | 33 | 0 |

**Astrocytes**

| | pfc | m1 | temp | a1 | s1 | par | v2 | v1 |
|---|---|---|---|---|---|---|---|---|
| **pfc** | 0 | 71 | 23 | 29 | 149 | 42 | 72 | 151 |
| **m1** | 22 | 0 | 30 | 3 | 4 | 1 | 8 | 8 |
| **temp** | 44 | 38 | 0 | 12 | 32 | 24 | 51 | 50 |
| **a1** | 98 | 24 | 6 | 0 | 95 | 59 | 46 | 86 |
| **s1** | 104 | 3 | 12 | 21 | 0 | 86 | 55 | 83 |
| **par** | 221 | 19 | 11 | 57 | 204 | 0 | 95 | 248 |
| **v2** | 81 | 11 | 29 | 26 | 36 | 6 | 0 | 23 |
| **v1** | 134 | 14 | 43 | 44 | 68 | 53 | 14 | 0 |

**Figure S5. Cortical rDEGs are defined across cell types and do not reflect ambient RNA contamination. (A)** *(i)* rDEG ambient score distributions (from CellBender) in temporal cortex samples, which had amongst the highest numbers of rDEGs compared to other neocortical regions. Despite glutamatergic neurons being more numerous and having more expressed genes/transcripts per cell, the median ambient contamination scores for glutamatergic rDEGs were not higher than median contamination scores for GABAergic rDEGs. rDEGs shared between glutamatergic and GABAergic neurons had indistinguishable scores compared with rDEGs private to one neuronal class. *(ii)* Ambient scores in temporal cortex of temporal cortex rDEGs are indistinguishable from V1 rDEGs. *(iii)* Distributions of temporal cortex rDEG ambient scores by neuron class, again showing no difference between rDEGs that are shared or private to a neuronal class. **(B)** Numbers of regionally differentially expressed genes (rDEGs) between pairs of cortical regions for neurons, astrocytes, and oligodendrocyte lineage types.

FIGURE S6

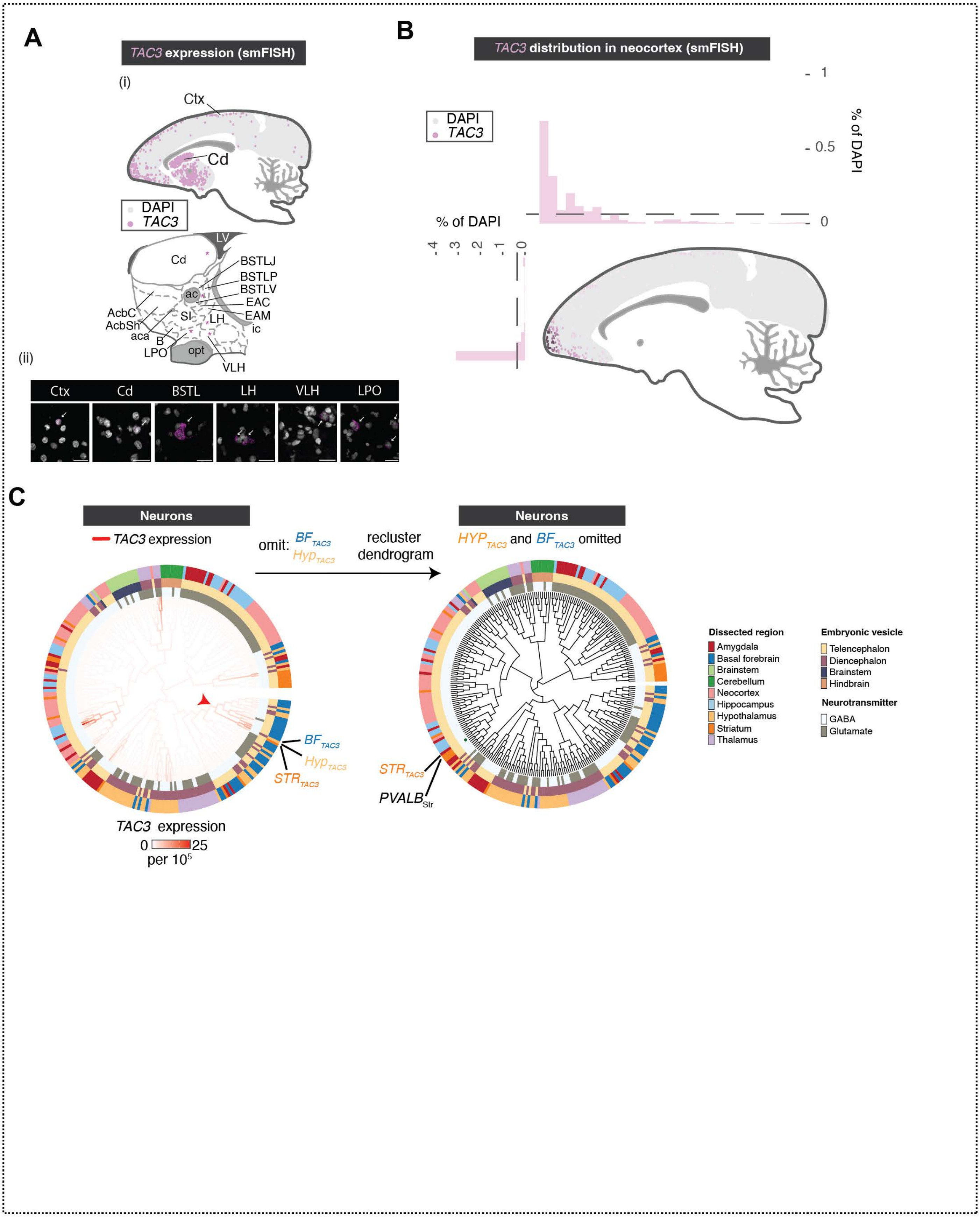

A
TAC3 expression (smFISH)
(i)
Ctx
Cd
DAPI
TAC3
LV
BSTLJ
BSTLP
BSTLV
ac
EAC
EAM
AcbC
AcbSh
SI
LH
ic
aca
B
LPO
opt
VLH
(ii)
Ctx
Cd
BSTL
LH
VLH
LPO
B
TAC3 distribution in neocortex (smFISH)
% of DAPI
DAPI
TAC3
C
Neurons
TAC3 expression
omit: BF TAC3 Hyp TAC3
recluster dendrogram
HYP TAC3 and BF TAC3 omitted
BF TAC3
Hyp TAC3
STR TAC3
PVALB Str
TAC3 expression
0 25
per 10^5
Dissected region
Amygdala
Basal forebrain
Brainstem
Cerebellum
Neocortex
Hippocampus
Hypothalamus
Striatum
Thalamus
Embryonic vesicle
Telencephalon
Diencephalon
Brainstem
Hindbrain
Neurotransmitter
GABA
Glutamate

**Figure S6. Locations of *TAC3*+ cells in marmoset forebrain. (A)** smFISH reveals anatomical locations and expression levels *TAC3+* types in different brain regions. *(i)* Schematic of *TAC3*+ cells imaged across cortex, dorsal striatum. Ctx = neocortex, Cd = Caudate. *(ii)* Cartoon close-up of nuclei in striatum, basal forebrain and hypothalamus. Magenta stars = locations of *TAC3*+ cells in lower image panel. Cd = Caudate, AcbC = nucleus accumbens core, AcbSh = nucleus accumbens shell, SI = Substantia innominata, B = basal nucleus of Meynert, EAM = extended amygdala, medial, EAC = extended amygdala, central, ac = anterior commissure, BSTLP = bed nuc st, lateral posterior, BSTLJ = bed nuc st, juxtacap, BSTLV = bed nuc st, lateral ventral, LH = lateral hypothalamus, VLH = ventrolateral hypothalamus, LPO = lateral preoptic area. **(B)** Density and location of *TAC3*+ cells as proportion of all DAPI+ cells. Barplots show percentages in bins (approximately 1,290 µm per bin ) taken across the anterior-posterior (top) and dorsal-ventral (left side) axes. **(C)** Effect on placement of the *TAC3+* striatal type on the neuronal dendrogram when omitting the two *TAC3*+ types in hypothalamus and basal forebrain. When these types are omitted and hierarchical clustering is repeated (using $HCA_{PCA}$), the *TAC3*+ striatal type is most similar to *PVALB*+ striatal interneurons, consistent with previous reports that only compared telencephalic interneurons (*3*).

FIGURE S7

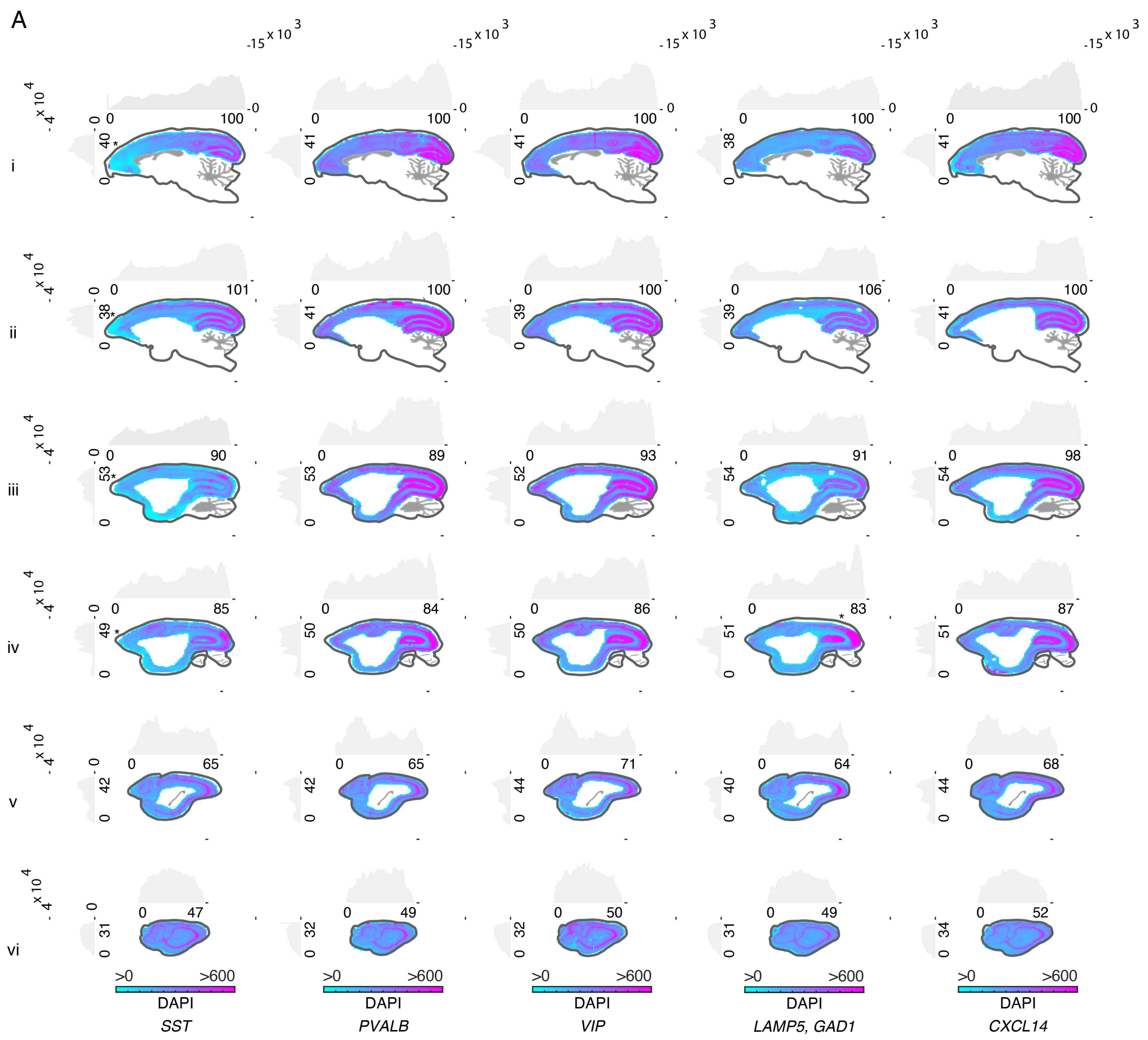

**Figure S7. Total cell numbers across marmoset neocortex. (A)** Total numbers of DAPI+ cells per unit area (approximately 387 μm per bin) for each of the sections shown in **Fig. 4C**.

FIGURE S8

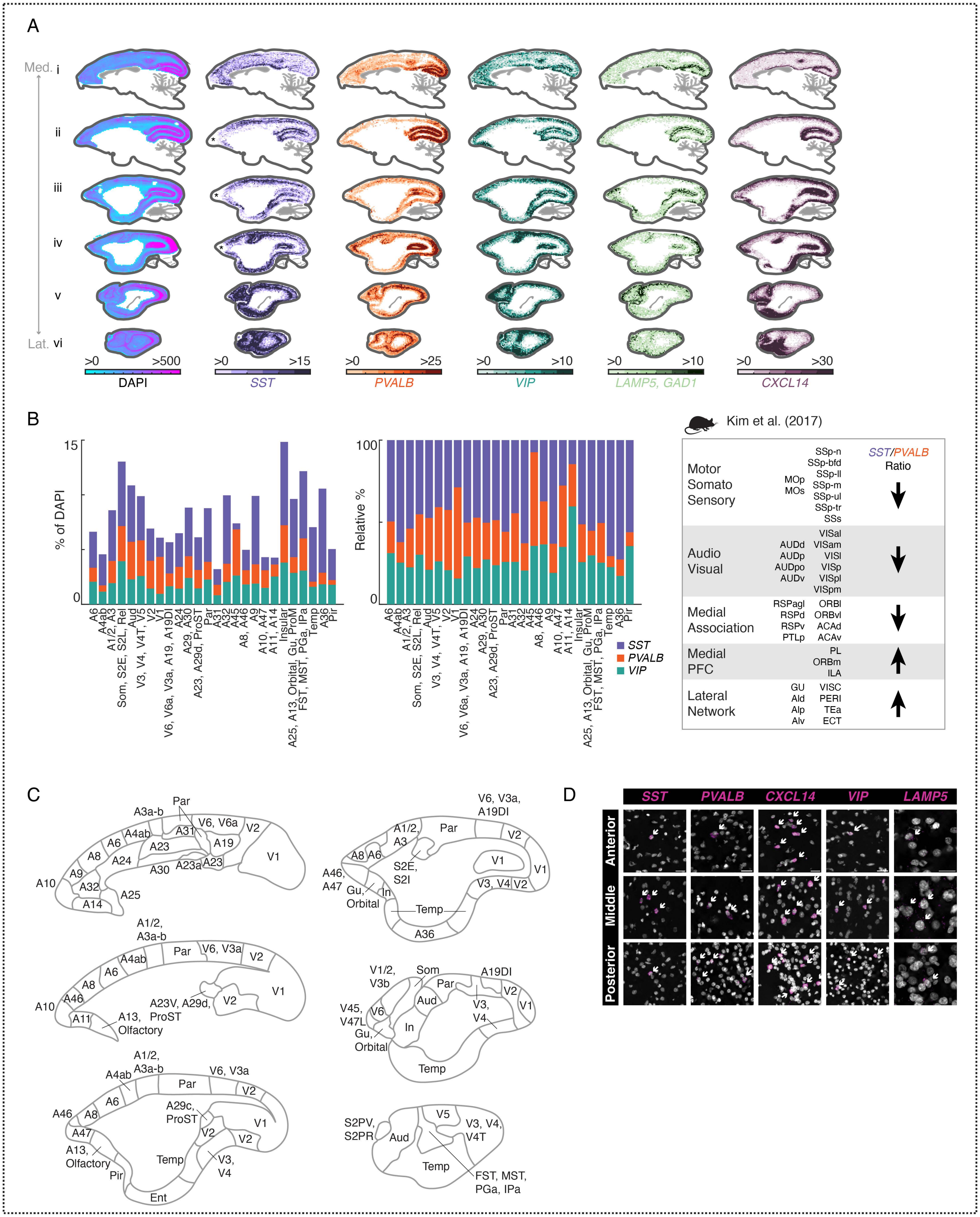

A
Med.
Lat.
i
ii
iii
iv
v
vi
>0 >500 DAPI
>0 >15 SST
>0 >25 PVALB
>0 >10 VIP
>0 >10 LAMP5, GAD1
>0 >30 CXCL14
B
% of DAPI
Relative %
SST
PVALB
VIP
A6
A4ab
A1/2, A3
Som, S2E, S2L, Rel
Aud
V3, V4, V4T, V5
V2
V1
V6, V6a, V3a, A19, A19DI
A24
A29, A30
A23, A29d, ProST
Par
A31
A32
A45
A8, A46
A9
A10, A47
A11, A14
Insular
A25, A13, Orbital, Gu, ProM
FST, MST, PGa, IPa
Temp
A36
Pir
Kim et al. (2017)
SST/PVALB Ratio
Motor Somato Sensory
MOp MOs
SSp-n SSp-bfd SSp-ll SSp-m SSp-ul SSp-tr SSs
Audio Visual
AUDd AUDp AUDpo AUDv
VISal VISam VISl VISp VISpl VISpm
Medial Association
RSPagl RSPd RSPv PTLp
ORBl ORBvl ACAd ACAv
Medial PFC
PL ORBm ILA
Lateral Network
GU AId AIp AIv
VISC PERI TEa ECT
C
D
SST
PVALB
CXCL14
VIP
LAMP5
Anterior
Middle
Posterior

**Figure S8. Interneuron numbers across marmoset neocortex. (A)** smFISH for neocortical interneuron subclass markers showing locations of cells positive for each marker across 6 sagittal sections of the marmoset neocortex. Heatmap scale shows absolute density per unit area (approximately 387 µm per bin). First column shows DAPI and area profiled. **(B)** (*Left, middle)* Quantitation of interneuron proportions by cortical area in marmoset parcellated according to Fig. S8C. *(Right)* Quantitation of interneuron proportions by cortical area in mouse reproduced from Kim et al. (2017). (*Left*) Absolute percentages of *SST, PVALB,* and *VIP* populations. (*Middle*) Same as (*left),* but scaled as proportions to 100%. (*Right*) Schematic describing ratios of major interneuron types (*Sst+, Pvalb+*) in mouse from Kim et al. (2017). **(C)** Cartoons of cortical areas and areal groupings used to bin smFISH neocortical interneuron proportions in (B) and **Fig. 4D-E**. Neocortical parcellation from https://doi.org/10.24475/bma.4520. **(D)** Examples of smFISH images quantitated in **Fig. 4B-E**. Panels for each marker show example positive cells in anterior, middle, and posterior locations across neocortex. White arrows indicate positive cells. Scale bar = 20 µm.

### FIGURE S9

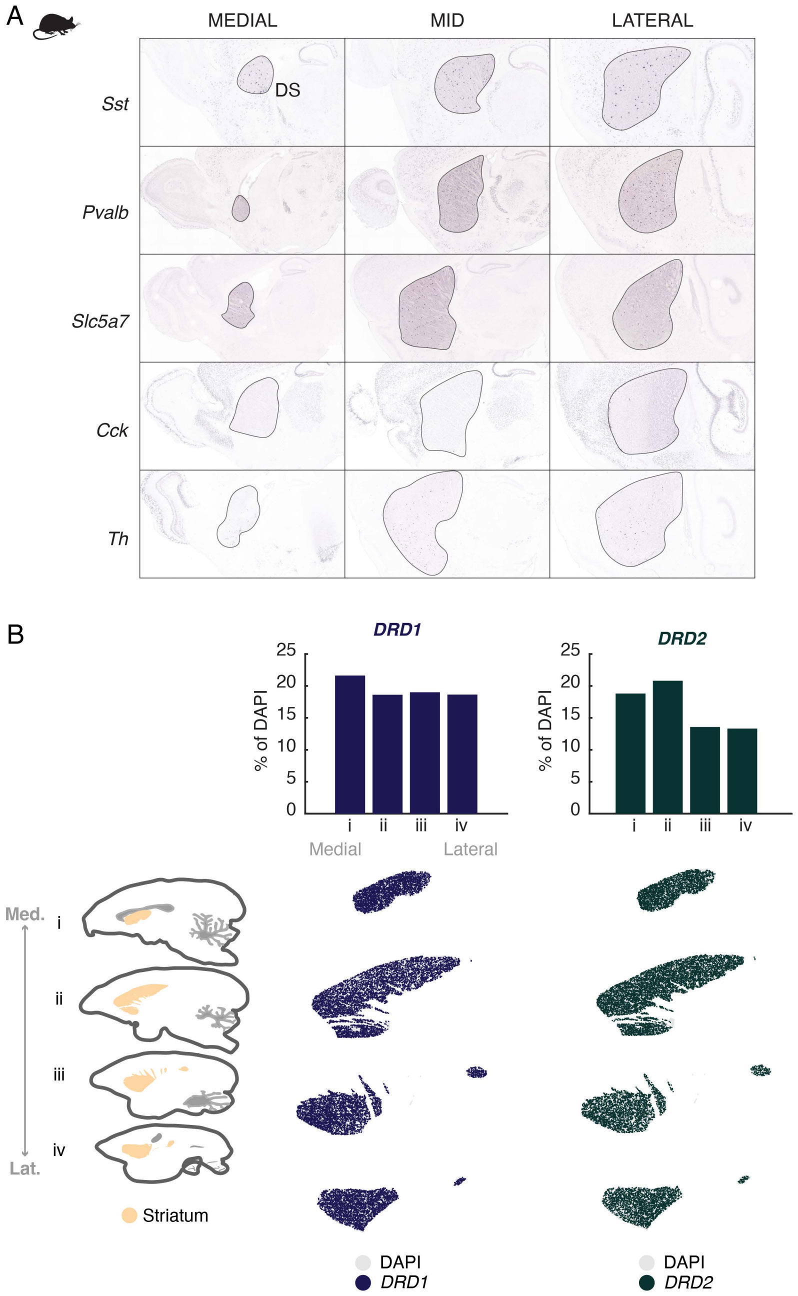

A
MEDIAL
MID
LATERAL
Sst
DS
Pvalb
Slc5a7
Cck
Th
B
DRD1
DRD2
% of DAPI
25
20
15
10
5
0
i
ii
iii
iv
Medial
Lateral
Med.
Lat.
Striatum
DAPI
DRD1
DAPI
DRD2

**Figure S9. Interneuron numbers across marmoset striatum. (A)** Mouse striatal *in situ* from Allen Brain Atlas for *Sst, Pvalb, Slc5a7, Cck, Th*. **(B)** Cartoon of marmoset striatum illustrates area profiled, medial to lateral. Proportions of *DRD1*+ (dark blue) and *DRD2*+ (dark green) cells across primate striatum calculated as in **Fig. 4H**.

FIGURE S10

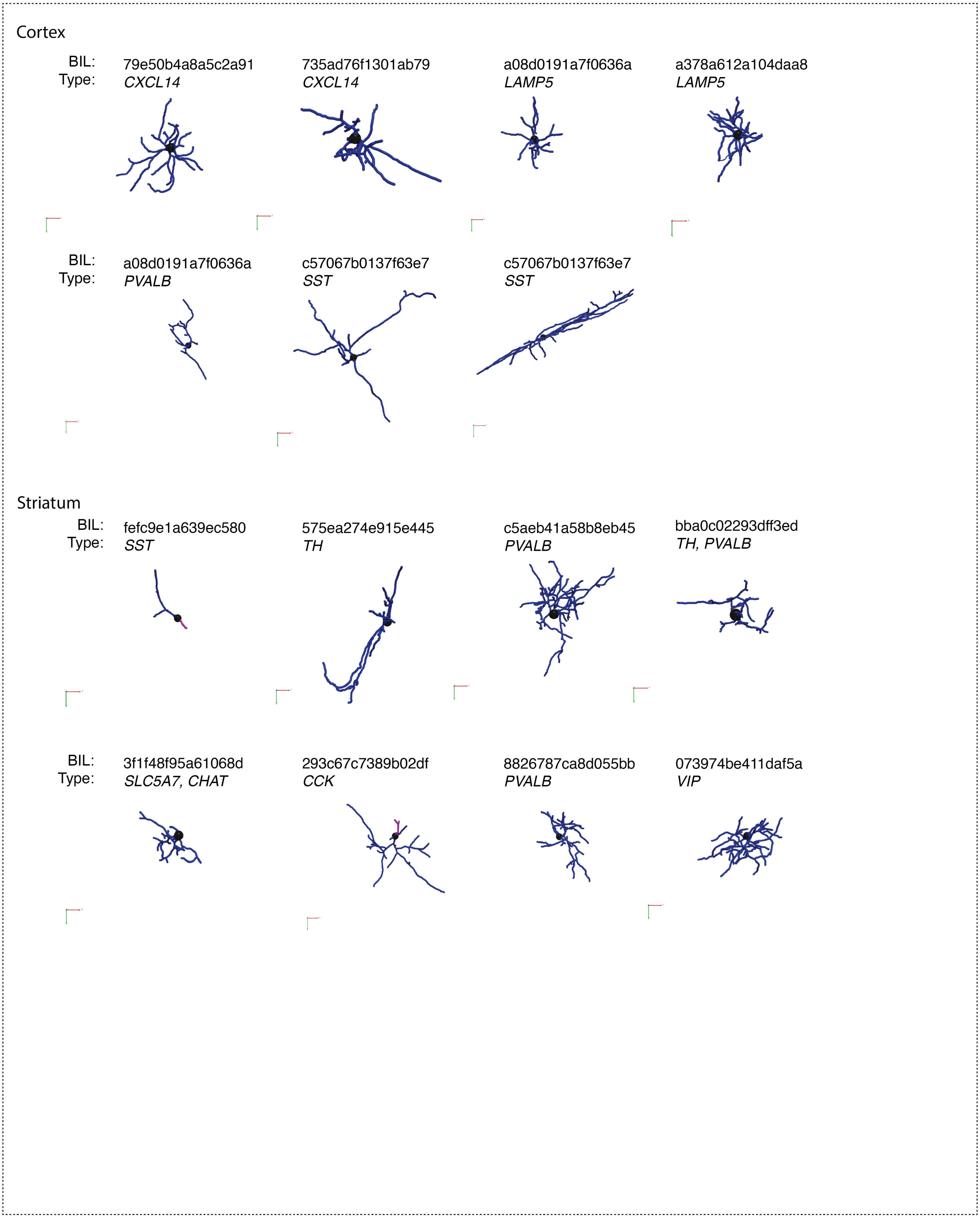

Cortex
BIL:
Type:
79e50b4a8a5c2a91
CXCL14
735ad76f1301ab79
CXCL14
a08d0191a7f0636a
LAMP5
a378a612a104daa8
LAMP5
BIL:
Type:
a08d0191a7f0636a
PVALB
c57067b0137f63e7
SST
c57067b0137f63e7
SST
Striatum
BIL:
Type:
fefc9e1a639ec580
SST
575ea274e915e445
TH
c5aeb41a58b8eb45
PVALB
bba0c02293dff3ed
TH, PVALB
BIL:
Type:
3f1f48f95a61068d
SLC5A7, CHAT
293c67c7389b02df
CCK
8826787ca8d055bb
PVALB
073974be411daf5a
VIP

**Figure S10. Morphology examples using NeuTube reconstructions. (A)** Example morphological reconstructions of striatal and neocortical interneurons using the NeuTube pipeline. Each cell, along with associated smFISH staining, is available for download at https://doi.org/10.35077/g.609.

Figure S11

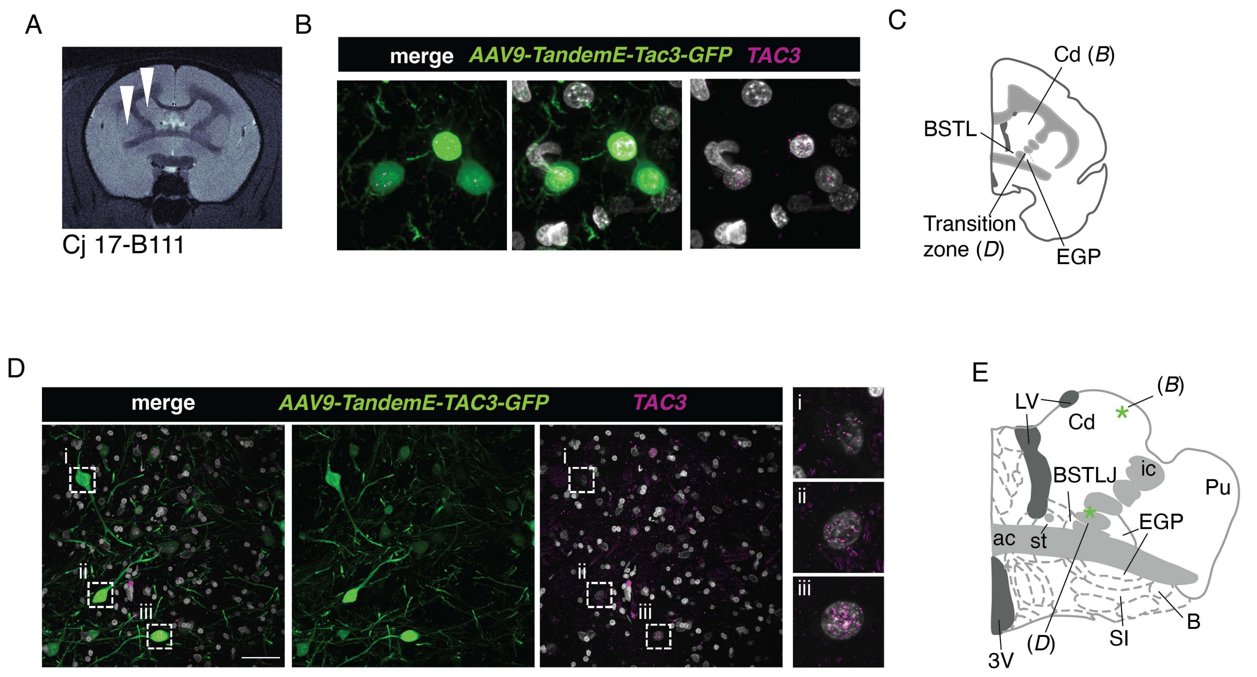
A
Cj 17-B111
B
merge AAV9-TandemE-Tac3-GFP TAC3
C
Cd (B)
BSTL
Transition zone (D)
EGP
D
merge AAV9-TandemE-TAC3-GFP TAC3
i
ii
iii
E
LV
Cd
(B)
BSTLJ
ic
Pu
EGP
ac
st
3V
(D)
SI
B

**Figure S11. Examples of striatal and peri-striatal *TAC3*+ neurons labeled by AAV-tandemE-TAC3-EGFP. (A)** MRI showing injection location (white arrowheads) of virus into bilateral dorsal striatum (caudate) in one animal (Cj 17-B111). **(B)** EGFP antibody-amplified confocal image of a labeled cell (position shown in (C) with smFISH for *TAC3* showing colocalization). **(C)** Cartoon showing location of positive cells in (B) as well as labeled cells in transition zone (D). **(D)** Examples of extra-striatal labeled cells from injections in (A). **(E)** Position of cells in (D). Cd = Caudate, SI = Substantia innominata, B = basal nucleus of Meynert, ac = anterior commissure, BSTLP = bed nuc st, lateral posterior, BSTLJ = bed nuc st, juxtacap, 3V = third ventricle, LV = lateral ventricle.

**Table S1. Marmoset sample information.** Table of animals, metadata, and experimental information.

**Table S2. snRNA-seq dataset by donor and brain area.** Tables of the number of cells per brain structure and samples per donor.

**Table S3. Neocortical rDEGs across three donors.** Neocortical rDEGs across three donors with pairwise comparisons between neocortical locations for major clusters of cortical excitatory neurons, inhibitory neurons, astrocytes and oligodendrocyte lineage types.

**Table S4. Fluorescent *in situ* hybridization probes and antibodies used.** A list of all FISH probes (RNA-Scope and Molecular Instruments) and antibodies used for validation of gene and protein expression *in situ*.

**Table S5. Morphological reconstructions performed with Neutube.** A list of all cells reconstructed with Neutube with their corresponding morphological measurements.

**Table S6. Morphological reconstructions performed with Imaris.** A list of all cells reconstructed with Imaris with their corresponding morphological measurements.

**Table S7. Links and DOIs.** A list of all links and DOIs referenced.

**Data S1. Striatal and cortical medio-lateral quantification.** Tabular data for striatal and cortical medio-lateral histograms/bar plots.

**Data S2. *TAC3* medio-lateral quantification.** Tabular data for *TAC3* medio-lateral histograms and bar plots.
